# Size- and stage-dependence in cause-specific mortality of migratory brown trout

**DOI:** 10.1101/544742

**Authors:** Chloé R. Nater, Yngvild Vindenes, Per Aass, Diana Cole, Øystein Langangen, S. Jannicke Moe, Atle Rustadbakken, Daniel Turek, L. Asbjørn Vøllestad, Torbjørn Ergon

**Affiliations:** Centre for Ecological and Evolutionary Synthesis (CEES), Department of Biosciences, University of Oslo, Oslo, Norway; Zoological Museum, The Natural History Museums and Botanical Garden, University of Oslo, Oslo, Norway; School of Mathematics, Statistics and Actuarial Science, University of Kent, Canterbury, England; Norwegian Institute for Water Research (NIVA), Oslo, Norway; Norconsult AS, Hamar, Norway; Department of Mathematics and Statistics, Williams College, Williamstown, Massachusetts, United States

**Keywords:** Bayesian statistics, dam, harvesting, hazard rate, mark-recapture, mortality, NIMBLE, trout

## Abstract

1. Evidence-based management of natural populations under strong human influence frequently requires not only estimates of survival but also knowledge about how much mortality is due to anthropogenic versus natural causes. This is the case particularly when individuals vary in their vulnerability to different causes of mortality due to traits, life-history stages, or locations.
2. Here, we estimated harvest and background (other cause) mortality of a landlocked migratory salmonid over half a century. In doing so, we quantified among-individual variation in vulnerability to cause-specific mortality resulting from differences in body size and spawning location relative to a hydropower dam.
3. We constructed a multistate mark-recapture model to estimate harvest and background mortality hazard rates as functions of a discrete state (spawning location) and an individual time-varying covariate (body size). We further accounted for among-year variation in mortality and migratory behavior and fit the model to a unique 50-year time-series of mark-recapture-recovery data on brown trout (*Salmo trutta*) in Norway.
4. Harvest mortality was highest for intermediate-sized trout, and outweighed background mortality for most of the observed size range. Background mortality decreased with body size for trout spawning below the dam and increased for those spawning above. All vital rates varied substantially over time, but a trend was evident only in estimates of fishers’ reporting rate, which decreased from over 50% to less than 10% throughout the study period.
5. We highlight the importance of body size for cause-specific mortality and demonstrate how this can be estimated using a novel hazard rate parameterisation for mark-recapture models. Our approach allows estimating effects of individual traits and environment on cause-specific mortality without confounding, and provides an intuitive way to estimate temporal patterns within and correlation among different mortality sources.

## Introduction

Population dynamics – particularly of long-lived species – are often highly sensitive to changes in mortality (Sæther and Bakke 2000). Mortality can have a wide variety of causes (e.g. starvation, predation, disease, harvest), and vulnerability to cause-specific mortality may depend on individual factors such as age or life stage (Ronget et al. 2017). As a consequence, population-level responses to changes in mortality may vary greatly depending on the underlying cause, and disentangling different causes of mortality may provide insights crucial for population management and conservation (Williams et al. 2002). This is particularly important in populations where a significant portion of mortality is linked to human activity; in such cases, knowledge about the relative impact of human-induced mortality and its effects on other mortality sources is crucial for developing sustainable and successful management strategies (Hilborn and Walters 2013, Koons et al. 2014).

Studies of marked individuals constitute a highly valuable source of demographic data for wild animal populations and are essential for estimating survival, as well as cause-specific mortality. The recovery of a dead marked animal often provides information on the cause of death. However, unless animals are marked with radio- or satellite transmitters, most dead individuals will not be found, and this imperfect detection needs to be accounted for when estimating mortality parameters. Moreover, when considering multiple mortality causes, detection probability frequently depends on the cause of mortality, and some causes of mortality may not be observable at all. This is usually the case for natural mortality when dead recoveries are exclusively based on the reports of hunters or fishers (e.g. Servanty et al. 2010, Koons et al. 2014).

Schaub and Pradel (2004) developed a multistate mark-recapture-recovery framework that allows separately estimating mortality from different causes while accounting for cause-dependent detection probabilities. Specifically, cause-specific mortalities are estimated as transitions from an “alive” state to several “dead from cause of interest” states. When this framework is extended to also include multiple “alive” states, it becomes possible to estimate differences in vulnerability to cause-specific mortality depending on, for example, an individual’s life-stage (e.g. juveniles vs. adults, Schaub and Pradel 2004) or location (Fernández-Chacón et al. 2015). Such group-level differences in mortality can be tremendous and accounting for them is crucial for modelling population dynamics (Ronget et al. 2017). However, in addition to that, vital rates and population dynamics are often strongly affected by individual differences in continuous, dynamic traits such as body size (De Roos et al. 2003, Vindenes and Langangen 2015). Particularly in species that are harvested and/or have indeterminate growth (e.g. fish species), cause-specific mortality is expected to depend strongly on body size. Fernández-Chacón et al. (2017) demonstrated this by estimating cause-specific mortalities for different sizes of Atlantic cod (*Gadus morhua*). However, they did so by lumping individuals into either of two size classes (“small” or “large”), thus foregoing the possibility of investigating the continuous relationship between body size and mortality from different causes. Knowledge about the relationships between continuous traits and vital rates is, however, invaluable for studying population-level trait dynamics (e.g. using integral projection models Ellner and Rees 2006).

Migratory salmonid fishes are extensively studied due to their ecological, cultural and economical value (Drenner et al. 2012). Nonetheless, studies at the population level are frequently hindered by a lack of knowledge about the mortality of adults residing in the sea or large lakes (Piccolo et al. 2012). Many salmonid populations are heavily impacted by human activity, not only in the form of harvesting, but also through pollution, fish farming, habitat fragmentation, and hydro-electrical power production (dams) in rivers (Aas et al. 2010), making the study of population-level consequences of such impacts a priority.

Here we study a population of migratory brown trout (*Salmo trutta*, hereafter “trout”) which inhabits a river-lake system in Eastern Norway and has been a popular target for fishing for decades due to its large body size. The spawning river is dammed, and trout migrating to spawning grounds above the dam face additional risks linked to dam passage on their up- and downriver migrations. Trout spawning below the dam, on the other hand, completely avoid these risks but may, in turn, incur costs related to poor river condition and crowding on the spawning grounds below the dam. Mortality risks are thus likely associated with spawning location in addition to individual body size and environmental conditions. To account for this heterogeneity, we re-parameterized mark-recapture models for cause-specific mortality in terms of mortality hazard rates (Cox 1972, Quinn 2003, Ergon et al. 2018) and extended the framework to include a continuous individual- and time-varying trait (body size) as a predictor of vulnerability within groups of individuals with different migration patterns. Fitting the resulting model to a unique 50-year time-series of recaptures and recoveries of marked trout enabled us to investigate the effects of individual (size, spawning location, origin) and environmental (river discharge) factors on, and temporal variation in, several key vital rates: the vulnerability of adult trout to mortality due to harvest, dam passage, and natural causes, and the probability of using a fish ladder within the dam to access upriver spawning areas.

## Materials and methods

### STUDY SYSTEM AND DATA

The studied population of landlocked migratory (potamodromous) brown trout inhabits the lake Mjøsa and its main inlet river, Gudbrandsdalslågen, in Eastern Norway. Eggs are deposited in the river in fall and develop over winter. After hatching in spring, juvenile trout remain in the river for an average of 4 years before smolting and migrating to the lake. They typically mature after 2 - 3 years of piscivorous diet and fast growth in the lake, and from that point on migrate up the river to spawn every other year (usually in August/September, Figure 1). See Aass et al. (1989) for a more detailed description of the life history. The population consists of wild-hatched trout and stocked (first-generation hatchery-reared) trout, which are recognizable by their clipped adipose fin. Stocked trout are released into the river and lake as smolts but then follow the same general life history as wild-hatched individuals (Aass 1993).

**Figure 1:**
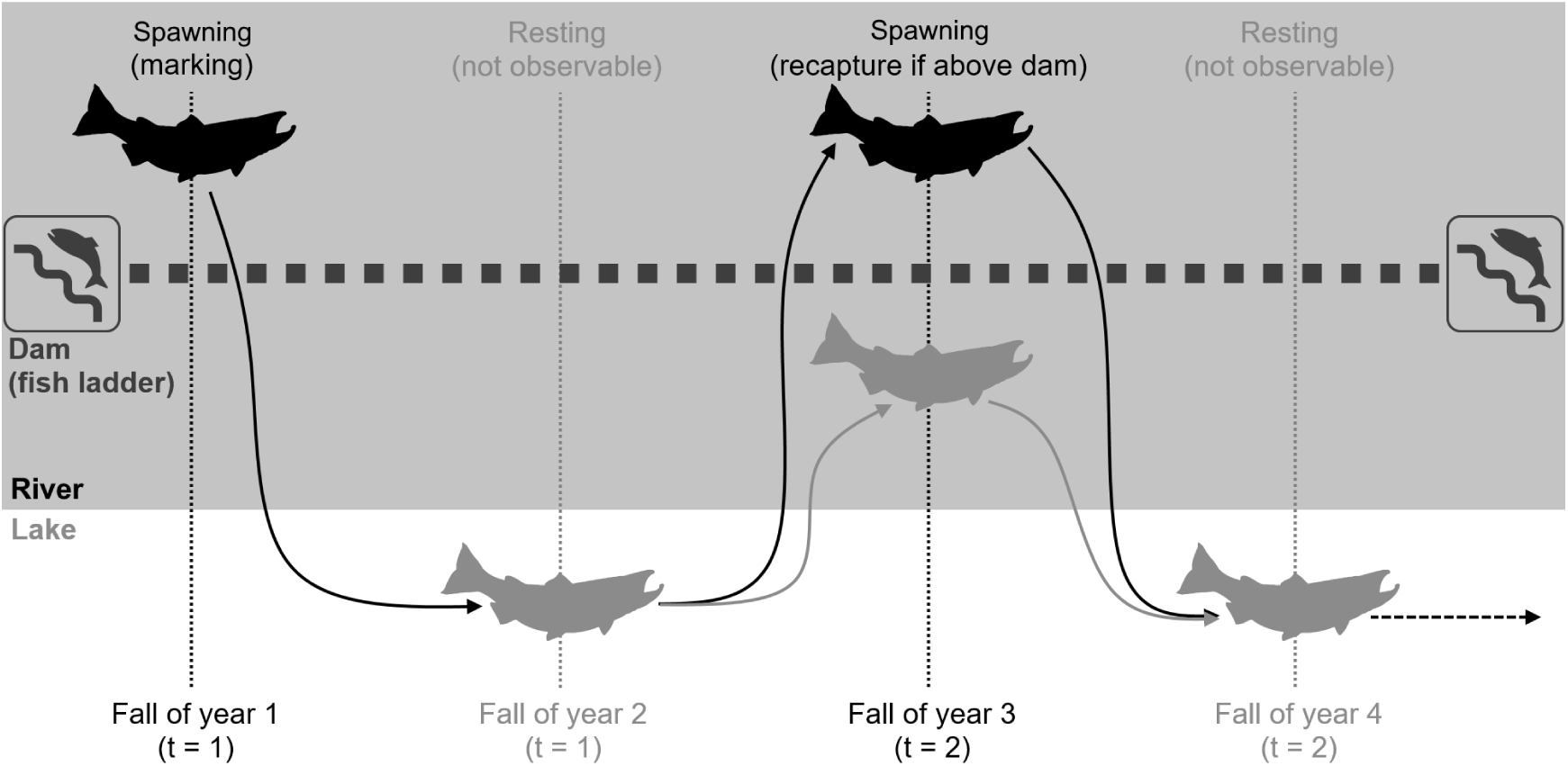
Illustration of the biennial spawning cycle and mark-recapture scheme of the studied trout population. All individuals are marked in the fish ladder while passing the dam on an upriver spawning migration. Two years later they may be recaptured on the next spawning migration, but only if they pass the fish ladder to spawn above the dam (if they spawn below the dam, they are unobservable). Trout remain in the lake and are unobservable during non-spawning years.

Shortly after the river was dammed in the 1960’s, a fish ladder was installed to restore connectivity to the spawning grounds above the dam. Depending on body size and hydrological conditions, trout may either pass the dam by using the fish ladder on their upriver spawning migration, or reproduce below the dam (Aass et al. 1989, Haugen et al. 2008). Trout spawning above the dam have to pass the dam again on their return migration to the lake (in October/November or in the following spring). Since the fish ladder cannot be used for moving downriver, these trout must pass either through the floodgates or the turbine shaft. Whether or not an individual uses the fish ladder thus determines not only its spawning location, but also the potential risks it encounters during the return to the lake.

From 1966 to 2016 a trap was operated within the fish ladder, allowing for all trout passing the ladder to be captured, measured, and individually marked. Thus, all adult trout were marked with Carlin tags (Carlin 1955) when they used the fish ladder on an upriver spawning migration for the first time, and could be recaptured on subsequent spawning migrations given that they passed the ladder again. Subsequent spawning runs occur two years later for the majority of fish (98.5%), which adhered to a strictly biennial spawning cycle (Figure 1). Over the 50-year time period, 13,975 adult trout were marked and 2,106 of these were recaptured in the ladder later. Since the population has been exposed to fishing over the entire time period, an additional 2,322 marked trout were reported dead by fishers. For more details on the marking scheme, sampling protocol, and resulting data from the mark-recapture-recovery study, see Moe et al. (2019).

In the present study we performed mark-recapture analyses over intervals of two years, as estimating parameters for spawning and non-spawning years separately was not possible (due to trout being unobservable in non-spawning years, Figure 1). We thus summarised the data into individual capture histories *y_i,t_*, in which each time index *t* corresponds to a two-year time step (interval from current spawning year to next spawning year). For each time step, we coded three types of observations: 1 = alive and captured in the ladder, 2 = dead from harvest and reported, and 3 = not observed. We set *y_i,t_* = 1 when an individual was captured in the fish ladder in any month during time interval *t*. Harvest of trout happens year-round (Figure S1.1) and if an individual was harvested and reported at any point during interval *t* we set *y_i,t_* = 2, unless (a) the individual had also been caught in the fish ladder during interval *t* or (b) the harvest happened after August of the second year within the interval *t*. If either (a) or (b) was the case, we moved the harvest observation to the next interval such that *y_i,t_*_+1_ = 2. Furthermore, we excluded all individuals that did not follow a strictly biennial spawning cycle (1.5% of all individuals), did not have a single size measurement taken (<1%), or were of unknown origin (wild vs. stocked, <1%). The analyses presented here are based on the remaining 13,003 capture histories containing 1,498 trap recaptures and 2,252 harvest recoveries from both wild-hatched and stocked (hatchery-reared) trout.

### MODEL FORMULATION

#### General model structure

We analysed the trout mark-recapture-recovery data in a multistate mark-recapture framework (Lebreton et al. 1999) with both “alive” and cause-specific “newly dead” states (Figure 2). Since trout are marked in the fish ladder while passing the dam on an upriver spawning migration, all individuals are in state 1, “spawning upriver”, at the start of their first 2-year time interval. State 1 individuals *i* may survive from the current (*t*) to the next (*t* + 1) spawning migration with probability *S*_1_*_,i,t_* and will then either use the fish ladder (probability *p_i,t_*_+1_) to spawn above the dam again, or remain below the dam for spawning (probability 1 *− p_i,t_*_+1_). Individuals using the ladder and thus remaining in state 1 are guaranteed to be observed, while individuals not using the ladder transition to state 2, “spawning downriver”, and are unobservable. Since spawning location may have a considerable effect on mortality, state 2 individuals have their own survival probability *S*_2_*_,i,t_*, but we assume that their probability of using the fish ladder during the next spawning run (*p_i,t_*_+1_) does not differ from that of state 1 individuals.

**Figure 2:**
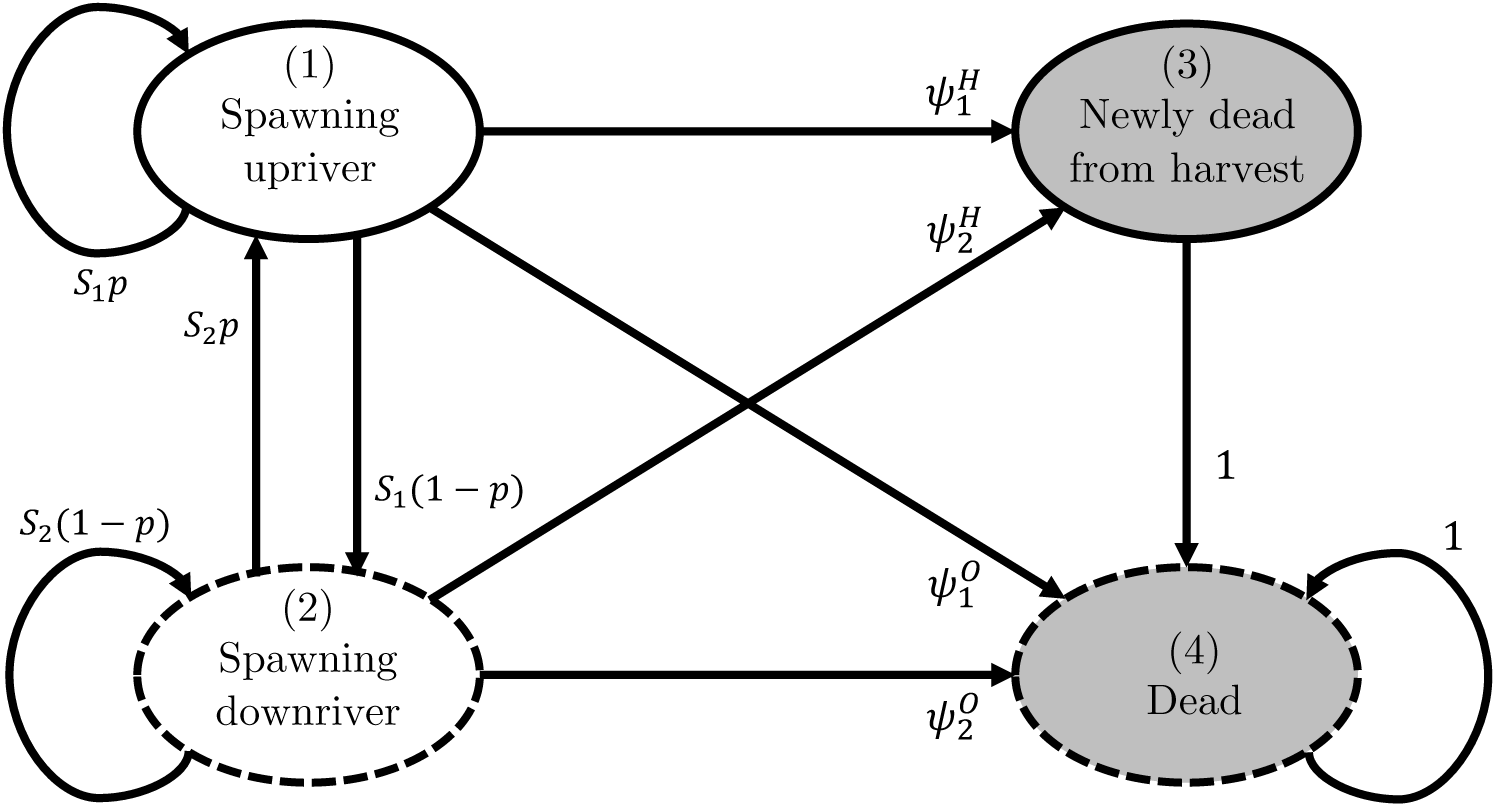
Design of the trout mark-recapture-recovery model (transitions on two-year intervals). White states are alive, grey states are dead. Solid borders indicate states that are at least partially observable, whereas dashed borders indicate unobservable states. *S_n_* = survival probabilities. 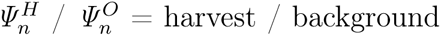 mortality probabilities (where *n* indicates the state). *p* = ladder usage probability. Indices for individual *i* and time *t* are omitted here for simplicity.

When deaths of marked individuals can be observed and attributed to a cause, multistate mark-recapture models can be used to estimate the probability of dying from cause X as the transition from an “alive” state to “newly dead from cause X” state (Schaub and Pradel 2004, Servanty et al. 2010). For the studied trout population, deaths due to harvest are clearly distinguishable from deaths due to other causes since fishers may report catching marked trout. Extending the model with the state “newly dead from harvest” (state 3) thus allows us to include the probability of an individual *i* in state *n* (*n* = 1 for above-dam spawners, *n* = 2 for below-dam spawners) dying due to harvest, 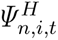, and dying due to other causes 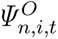 over the time interval *t* to *t* + 1. Individuals that have recently died due to harvest (state 3) may be reported by fishers with reporting rate *r_t_*. Individuals that die due to other causes are not observable and therefore transition directly to the “dead” state (state 4; see Figure 2).

The resulting multistate model for the trout mark-recapture-recovery data can be expressed with the following state transition matrix and associated observation probabilities:

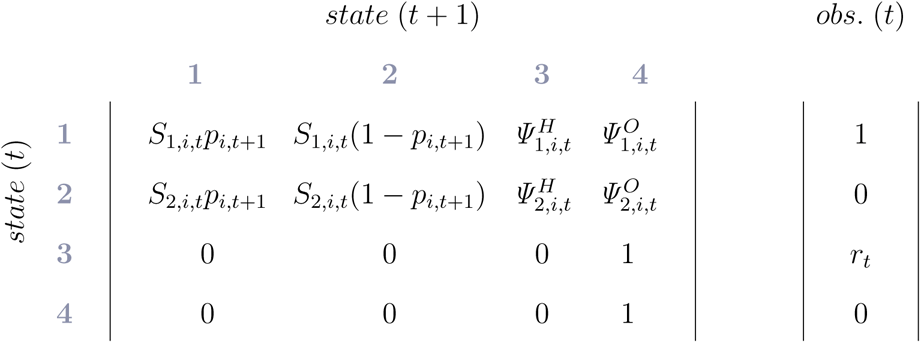

#### Parameterisation by mortality hazard rates

Different cause-specific mortality probabilities (*Ψ*) are not independent of one another; if a certain cause of mortality becomes more prevalent (e.g. due to some event or change in the environment), not only will the probability of dying from that cause increase, but the probability of dying from any other cause will decrease at the same time. This confounding complicates inference (e.g. Cooch et al. 2014), but Ergon et al. (2018) have recently re-emphasized that this can be avoided – also in the context of discrete-time mark-recapture analyses – by parameterising with mortality hazard rates instead of probabilities (Cox 1972, Quinn 2003). Assuming that the intensities of mortality from different causes remain proportional within time intervals, we can define the survival- and mortality probabilities in the trout model using harvest (*m^H^* ) and other-cause (hereafter “background”) mortality hazard rates (*m^O^* ):

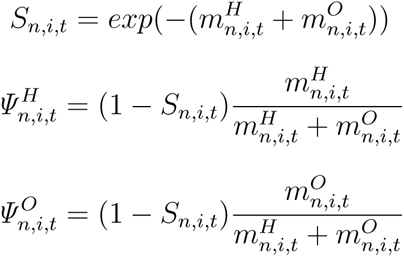

(see derivation in Ergon et al. (2018))

In the present implementation, we further constrained harvest mortality to be the same for trout spawning above and below the dam: 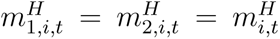 This constraint was necessary to obtain an identifiable model, but also biologically reasonable because most harvest happens in the lake and fishing in the river is restricted during the spawning season (which is also short relative to the two-year interval of analysis).

### MODEL IMPLEMENTATION

#### Individual and temporal variation in vital rate parameters

Body size and hydrological conditions are often key determinants of vital rate variation in freshwater fish, including our study population (e.g. Carlson et al. 2008, Letcher et al. 2015, Haugen et al. 2008). We thus used individual body size (length; mm) at the beginning of the time-interval and average river discharge during the relevant season as covariates for mortality and ladder usage in our model. We further accounted for potential effects of hatchery origin and additional among-year variation in all parameters *x* using intercept offsets for stocked individuals 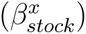 and temporal random effects 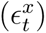, respectively. Random effects on all parameters were assumed to be independently normally distributed on the link scale (but see Supporting Information (SI) S6 for a model extension with correlated random effects).

Harvest in our study system has been done mostly using fishing rods or gillnets; the selectivity of the former is often positively correlated with body size (Lewin et al. 2006) while the latter typically have bell-shaped selectivity curves (Hamley 1975). Since we here pooled harvest by all gear types, we modelled linear and quadratic effects of size on harvest hazard rate on the log-scale:

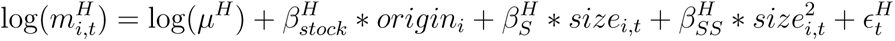

where *µ^H^* is the median harvest hazard rate, 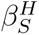 and 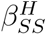 are slope parameters for linear and quadratic size effects respectively. *size_i,t_* is the individual length at spawning and *origin_i_* is a binary variable taking values of 1 for stocked fish and 0 for wild-hatched fish.

Background mortality, is expected to depend not just on body size but also on spawning location and river discharge, as above- and below-dam spawners encounter different hydrological conditions during/after spawning and only the former need to pass the dam on their downriver migration. Mortality associated with the spawning migration in general, and passing of the dam in particular, may also depend on body size. We thus modelled background mortality hazard rate as:

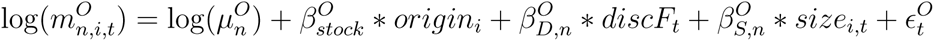

Here 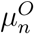 is the median background mortality hazard rate of state *n*, *discF_t_* is the average discharge during the fall when many post-spawned trout are expected to migrate downriver (Oct - Nov), and 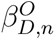 and 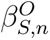 are slope parameters for discharge and size effects respectively. Stocking effects 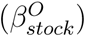 and temporal random effects 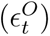 for background mortality are assumed to be shared across states *n*.

The probability of using the fish ladder and thus spawning above the dam was previously found to depend on a complex interplay of individual body size and river discharge (Haugen et al. 2008). We adopted the basic model structure from this earlier analysis and extended it by allowing for stocking effects and random among-year variation such that

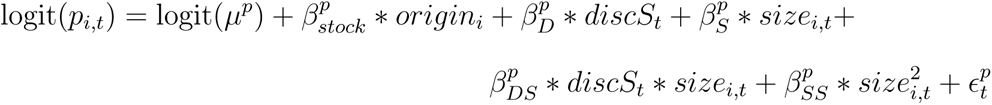

The discharge covariate used here, *discS_t_*, differs from the one used above and represents the average discharge over the summer season when trout undertake their upriver spawning migration (Jul-Oct).

#### Size imputation under imperfect detection

Using continuous, time-varying individual traits such as body size as covariates in mark-recapture models is problematic due to imperfect detection: information on body size will be missing for sampling occasions when an individual is not captured (Pollock 2002). There are several ways to approach this problem, including integrated growth models (e.g. Bonner et al. 2010) and inter-/extrapolation using other available data and/or separate models. Due to the prohibitively large computational demands of an integrated analysis, we here adopted the latter approach and used a detailed growth model previously developed for the study population of brown trout (Nater et al. 2018) to impute missing values in the individual size covariate. Specifically, we r e-fitted th e gr owth mo del of Na ter et al . (2 018) to an extended set of growth data from 6,843 individuals spanning the years 1952 to 2003 and used the resulting parameter estimates, plus a correction factor, to calculate all missing entries in the body size covariate. The imputation procedure, as well as implementation and results of the growth analysis, are described in detail in SI S5.

#### Autoregressive reporting rate model

Time-dependent reporting rate *r_t_* can be expected to vary considerably over a period of 50 years. To accommodate this, we followed the example of Zhao et al. (2018) and used a flexible, autoregressive model for time-dependent reporting rates:

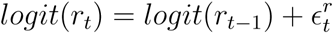

where 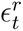 are independently normally distributed random effects. For details on the implementation of the autoregressive model in the context of the overlapping 2-year time-intervals in our model, we refer readers to the model code (supplementary file `nimbleDHMM.R`).

#### Implementation with NIMBLE

We implemented the model in a Bayesian framework in NIMBLE (de Valpine et al. 2017). Building on the work of Turek et al. (2016), we developed a highly efficient custom likelihood function to greatly reduce MCMC runtimes and memory load of our analysis (detailed description/evaluation of the custom implementation and code are provided in SI S2 and `nimbleDHMM.R`). To accommodate the 2-year interval of our analysis, we split the data into two sets containing only individuals spawning in even years and in odd years respectively. We then formulated the likelihood for both datasets separately, but analysed them jointly under the assumption of shared intercept-, slope-, and variance parameters. We used non-informative priors for all parameters, and made use of NIMBLE’s default set of samplers. The MCMC algorithm was run for 4 chains of 35,000 iterations, discarding the first 5,000 samples as burn-in. Analyses were run in `R 3.5.0` (R Core Team 2018) using version `0.6-13` of the `nimble` package (NIMBLE Development Team 2018).

### MODEL IDENTIFIABILITY AND FIT

With increasing model complexity, and particularly when unobserved states are included, it is not obvious whether all parameters within a multi-state mark-recapture model can be estimated (Lebreton and Pradel 2002, Gimenez et al. 2003). Using an extended (hybrid) symbolic method (Cole et al. 2010, Cole 2012, Choquet and Cole 2012) implemented in the computer algebra package Maple, we looked at intrinsic parameter redundancy in the above described model including different covariate- and random effect structures. Analyses of instrinsic parameter redundancy, as well as investigation of potential near-redundancy using prior-posterior overlap (Garrett and Zeger 2000, Gimenez et al. 2009), are described in detail in SI S3. Maple code is provided as supplementary material.

Subsequently, we tested the fit of our model to the data using posterior predictive checks (PPCs, Conn et al. 2018). Specifically, we selected 500 evenly spaced samples from our posterior distributions and used them to simulate 10 replicate mark-recapture-recovery datasets per sample. From each simulated dataset, we then extracted several test statistics representing numbers and size distributions of recaptured/harvested trout and compared them to the same quantities obtained from the real data using visual tools and Bayesian p-values. Methodology and results of the PPCs are described in detail in SI S4.

## Results

### MODEL IDENTIFIABILITY AND FIT

We found that in the absence of random effects, the only model structures that were intrinsically identifiable were those where harvest mortality depended on an individual time-varying covariate (e.g. body size) and background mortality was either constant or dependent on an environmental covariate (Table S3.1). However, all models (irrespective of covariate structure) became identifiable when random year effects were included on at least harvest hazard or reporting rates (Table S3.1). Prior-posterior overlaps were below 35% for all parameters except *r*_1_, indicating no major problems with near-identifiability (SI S3.3).

PPCs indicated that overall, the model produced a decent fit to the data, with Bayesian p-values for the majority of considered data properties falling into an acceptable range (0.10 - 0.90 for the whole dataset, 0.37 - 0.59 for averages across marking cohorts, SI S4.3). We found some evidence for lack of fit for a subset of data properties: mean/median size of individuals recaptured two years after marking and the number of individuals harvested two to four years after marking. In both cases, lack of fit was most pronounced in the beginning the time series (Figures S4.3 & S4.7). Graphical tools illustrated that the model’s predictions of whole size distributions were generally realistic despite Bayesian p-values for size mean, median, and standard deviation sometimes indicating some degree of lack of fit (Figure S4.4). For detailed PPC results, refer to SI S4.4.

### SIZE-DEPENDENT FISH LADDER USAGE

Posterior distributions for all estimated parameters are plotted in Figures S1.2 to S1.10. Numerical results in the following text are displayed as median [95% credibility interval].

The probability of using the fish ladder – and thus spawning above the dam – depended strongly on individual size and, to a lesser degree, on river discharge (Figure 3). Intermediate-sized trout (600-700 mm) were most likely to pass the dam under any discharge conditions. Small to intermediate-sized trout had a higher probability of using the ladder when river discharge was high, whereas the probability decreased markedly with size for larger trout irrespective of hydrological conditions. Ladder usage probability fluctuated considerably over time (Figure 4c) and was predicted to be lower for stocked (0.476 [0.414, 0.546]) than wild-hatched (0.533 [0.477, 0.592]) trout (Figure S1.11).

**Figure 3:**
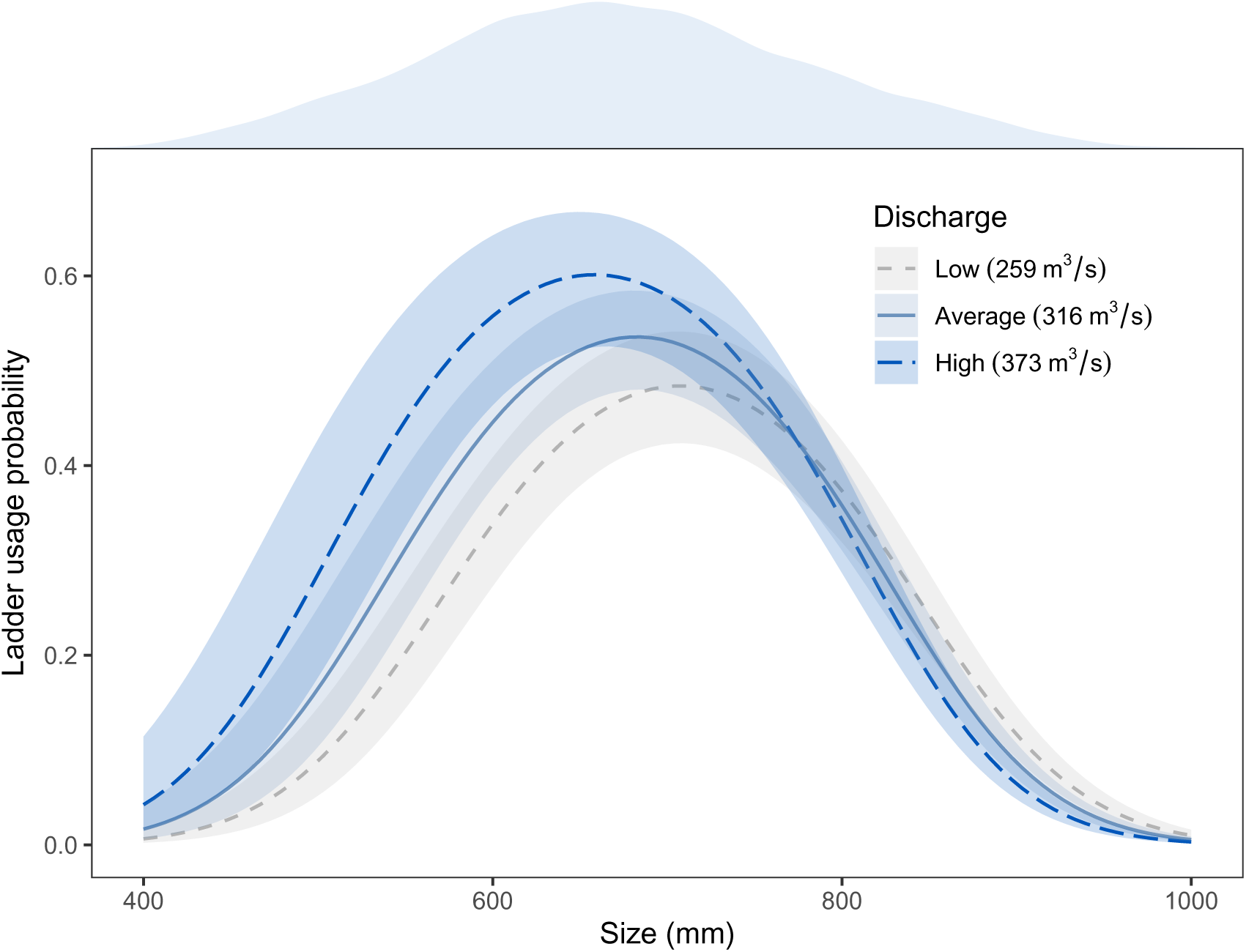
Predictions of the effects of body size on ladder usage probability at different levels of river discharge. Grey, dashed = low discharge (mean *−* SD). Grey-blue, solid = average discharge (mean). Blue, longdashed = high discharge (mean + SD). Lines represent median prediction, ribbons indicate 95% credibility intervals. The blue density kernel above the plot visualizes the size distribution of trout caught in the ladder (data).

**Figure 4:**
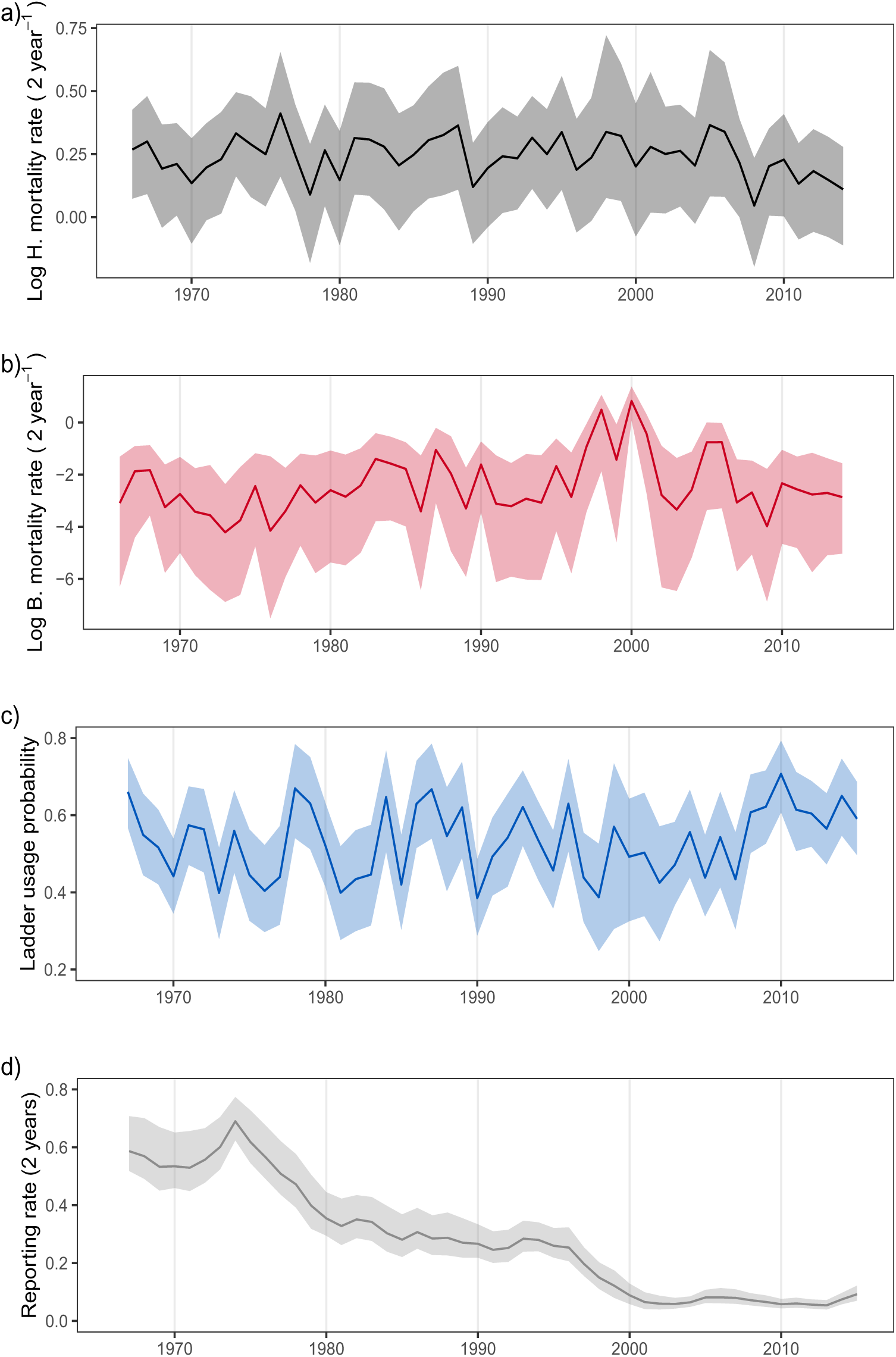
Estimates for time-dependent a) log harvest hazard rate, b) log background mortality hazard rate (above-dam spawners), c) ladder usage probability, and d) reporting rate (calculated using random variation and discharge effects). Lines represent median predictions, ribbons indicate 95% credibility intervals.

### CAUSE- AND SIZE-DEPENDENT MORTALITY

Median mortality hazard rates were estimated at 1.285 [1.090, 1.437] (harvest), 0.084 [0.021, 0.320] (background above-dam), and 0.115 [0.024, 0.540] (background below-dam) per two years for average-sized trout (670 mm). The resulting probabilities of dying during a 2-year interval due to harvest 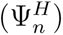 and due to other causes 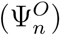 were 0.700 [0.600, 0.752] and 0.045 [0.011, 0.173] for above-dam spawners and 0.692 [0.561, 0.751] and 0.063 [0.013, 0.324] for below-dam spawners. Harvest hazard rate was predicted to be highest for individuals with a size around 550 mm (Figure 5a). Background mortality hazard rate, while mostly lower than harvest hazard rate, decreased with size for above-dam spawners and increased with size for below-dam spawners (Figure 5a). Consequently, total survival probability increased with size for all trout up to 870 mm, but flattened out for larger below-dam spawners (Figure 5b). River discharge was predicted to increase back-ground mortality of above-dam spawners only (Figure S1.2). Residual among-year random variation was substantial in harvest and especially background mortality, with hazard rates at the 97.5 percentile being 1.28- and 69.67-fold higher than at the 2.5 percentile respectively, but no temporal trends were evident in either mortality cause (Figures 4a & 4b).

**Figure 5:**
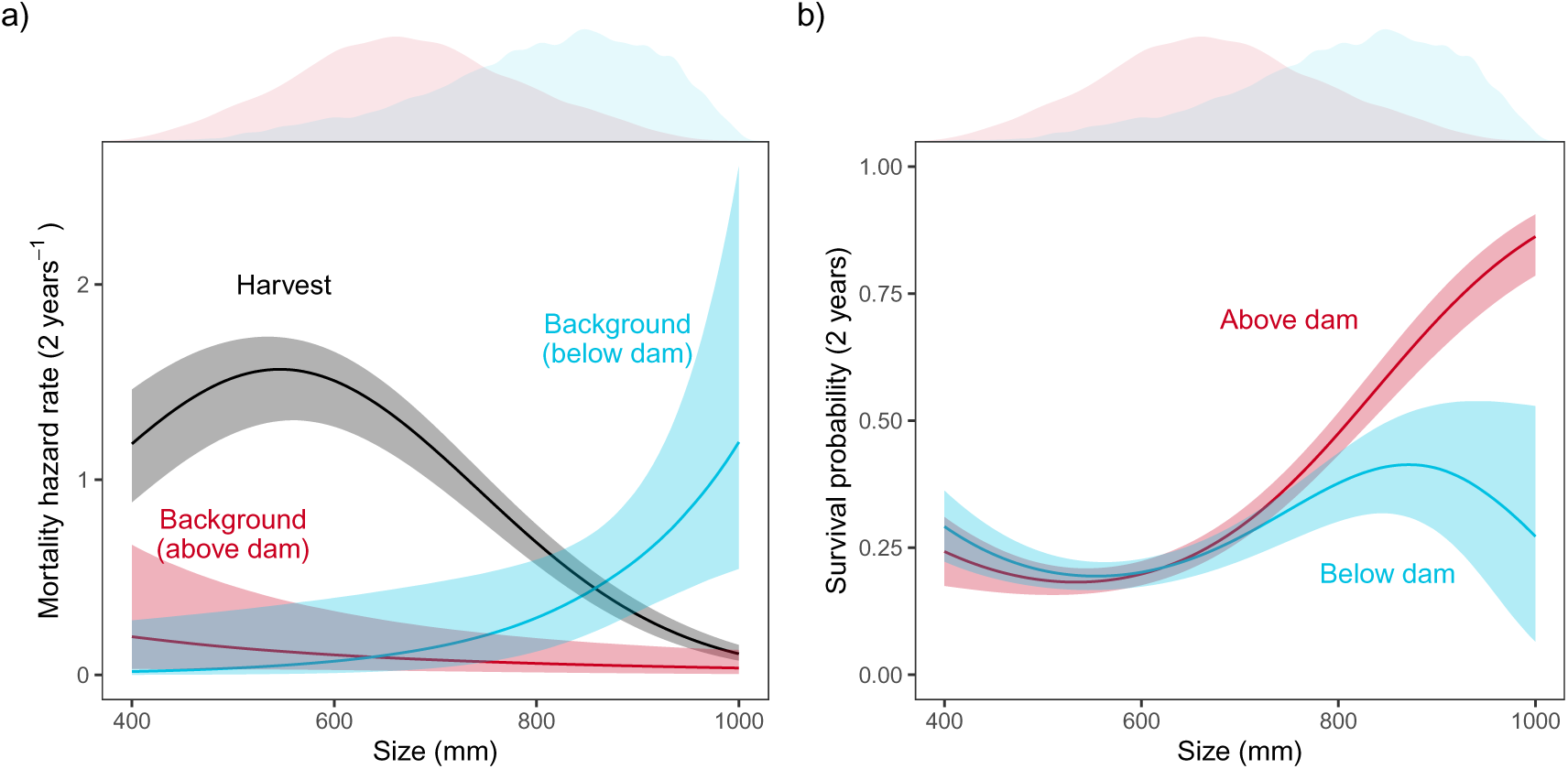
Predictions of the effects of body size on a) harvest and background mortality hazard rates and b) survival probabilities (under consideration of both mortality sources). Red and blue curves apply to individuals that have last spawned above and below the dam respectively. The black curve (harvest) applies to all individuals irrespective of their last spawning location. Lines represent median predictions, ribbons indicate 95% credibility intervals. Density kernels above the panels illustrate the informative data range: red = size distribution of individuals captured in the fish ladder (above-dam spawners, raw data), blue = simulated size distribution of unobservable below-dam spawners.

Model results did not support differences in harvest- or background mortality due to trout origin: hazard ratios of stocked and wild trout were 0.988 [0.886, 1.081] and 0.991 [0.617, 1.601] for harvest and background mortality respectively (Figure S1.11).

### TEMPORAL PATTERNS IN REPORTING RATE

A clear decrease in estimates of reporting rate over the 50-year time-period was evident (Figure 4d), with values exceeding 50% in early years but dropping below 10% towards the end of the time series.

## Discussion

Individuals can differ greatly in their vulnerability to mortality from different causes depending on traits like body size and variation in exposure to mortality risk (e.g. as a consequence of reproductive state or location). Particularly when some mortality causes are directly linked to human activity, understanding and accounting for such individual differences in vulnerability can be crucial for management and conservation. In this study, we combined recent advances in mark-recapture methodology and Bayesian modelling to investigate factors determining vulnerability of large migratory brown trout to harvest- and background mortality in a system heavily impacted by fishing and hydropower production.

### SIZE-DEPENDENCE OF CAUSE-SPECIFIC MORTALITY

Size-dependent survival is well documented for salmonid fishes like brown trout, but direction and strength of size effects vary widely across habitats, populations, years, and life history stages (Carlson et al. 2008, Drenner et al. 2012). Here, we were able to not only reproduce previous findings of positively size-dependent survival for the studied trout population (Figure 5b, Haugen et al. 2008), but to disentangle the underlying contributions from mortality due to harvest and other causes.

Model results supported our initial expectation of non-linear dependence of vulnerability to harvest and body size : harvest mortality was highest for trout with sizes of around 550 mm and decreased for both smaller and larger individuals (Figure 5a). Bell-shaped selectivity curves such as this are typical for gillnets (Hamley 1975), which have been commonly used in our study area. The low harvest mortality of large trout, however, may seem surprising given that 44% of the reported harvests were due to angling, which often targets larger fish (Lennox et al. 2017). This may indicate that large trout escape harvest either through their individual behavior (e.g. different foraging habitats and prey preferences, learning, Lewin et al. 2006, Arlinghaus et al. 2008) or because cohort selection favours more cautious fish, allowing them to survive and grow to large sizes (Lennox et al. 2017). Effects of body size on background mortality were predicted to be markedly different for trout spawning above and below the dam, in particular for larger trout (Figure 5a). Trout spawning above the dam generally had low background mortality, possibly indicating limited mortality risk associated with dam passage for adult fish. Nonetheless, smaller individuals were slightly more vulnerable to dying from non-harvest causes than larger ones (92% of posterior samples indicated a negative effect of size on background mortality, Figure S1.2). Two mechanisms that may be (partially) responsible for this are turbine mortality and energetic costs of dam passage. During downriver migration after spawning above the dam, trout have to pass through the floodgates or the turbine shaft to return to the lake. As on many hydroelectric dams, racks are installed in front of the Hunderfossen power plant’s turbine intake to prevent fish from entering, but small fish may slip through the grid and subsequently suffer severe injury and die passing the turbine (e.g. Fjeld-stad et al. 2018). Alternatively, smaller fish may have reduced survival following dam passage due to large energy expenditures resulting from dam passage (on up-and/or downriver migration) itself (e.g. Roscoe et al. 2011) or as a consequence of migration delays, particularly if these force individuals to overwinter in the river (Fjeldstad et al. 2018). Both of these mechanisms are plausible here when also considering that background mortality of above-dam spawners was predicted to increase at higher levels of river discharge (Figure S1.2): stronger water flow could increase both the risk of being swept into the turbine shaft and the energetic costs of passage.

Unlike trout spawning above the dam, trout spawning below the dam were predicted more vulnerable to background mortality at larger sizes (Figure 5a). Many mechanisms may be responsible for this; one possibility is related to trout density downriver of the dam, which can be very high during the spawning season (Kraabøl 2006) and likely results in elevated levels of stress, aggressive interaction, and disease transmission. Mortality below the dam could increase with body size if larger individuals (due to their size, age, or other traits correlated with large body size) were less able to cope with these challenges and/or increased their investment into reproduction at the cost of survival under adverse conditions. At the same time as having higher background mortality below the dam, large trout were also much more likely to spawn below the dam in the first place (Figure 3), and thus incur the resulting higher mortality. The hydropower dam therefore has the potential to function as an ecological trap (Schlaepfer et al. 2002) via its size-selective fish ladder and adverse conditions on downriver spawning grounds, particularly when considering that the reproductive output of large fish is often central to the viability of salmonid populations (Jonsson and Jonsson 2011).

A second, more practical consequence of the selectivity of the fish ladder is that it substantially limited comparisons of background mortality of above- and below-dam spawning trout of the same size in the present study. With small and large trout predominantly spawning above and below the dam respectively, direct comparisons are only informative for a relatively narrow size range (*∼* 700 – 850 mm). Within this range, predictions for above- and below-dam spawners mostly overlap, with the exception of the largest sizes (Figure 5). Additional data – particularly on the fates of individuals spawning below the dam – would be necessary for a more detailed assessment of the interactive effects of hydropower production and spawning location on mortality and for investigating potential mechanisms explaining higher mortality large fish below relative to above the dam. What our approach did allow, however, was an unbiased quantitative comparison of size-dependent harvest and background mortality: the risk of dying due to fishing was higher than the risk of dying due to any other cause for almost the entire size range, suggesting fishing as the main source of adult mortality in this population (see Kleiven et al. 2016, for a similar result on Atlantic cod).

### TEMPORAL VARIATION OVER 50 YEARS

The present analysis extended over half a century, in which the river-lake system experienced variation in abiotic and biotic factors due to river regulation, lake restoration, and changes in climate and human activities (Hobæk et al. 2012). It is therefore unsurprising that we found high among-year variation in cause-specific mortality and fish ladder usage over the course of the 50-year study period (Figure 4a-c). Background mortality in particular was subject to large fluctuations and displayed a marked increase during the period 1997-2001 (also visible in overall mortality and survival, Figure S1.12). This coincides with a documented outbreak of a fungal disease in the study population (*Saprolegnia* spp. infections, possibly in combination with ulcerative dermal necrosis, Johnsen and Ugedal 2001). This suggests that disease may be a key driver of changes in adult trout mortality and has the potential to substantially affect population viability (Hudson et al. 2002). Since freshwater ecosystems are particularly vulnerable to infectious diseases (Okamura and Feist 2011), studying fungal disease dynamics and how are affected by harvest, river regulation, and other environmental factors (e.g. temperature, Letcher et al. 2015) represents an important venue for future research.

Unlike cause-specific mortality and ladder usage, which displayed strong fluctuations but no obvious trends, fisher’s reporting rate decreased clearly and rapidly over time: from over 50% of catches being reported in the beginning of the study period to less than 10% in the last two decades (Figure 4d). Declining fisher engagement over time is a known problem in tagging studies without reward tags (Piccolo et al. 2012), and highlights the importance of maintaining volunteer participation in long-term studies by providing appropriate feedback and keeping up with technological development of tools and platforms for reporting (Dickinson et al. 2012).

### MODEL LIMITATIONS

When analysing long-term ecological data even complex hierarchical models, like the ones used here, can fail to sufficiently capture heterogeneity (overdispersion) in the data, resulting in lack of model fit (Richards 2008). PPCs (Conn et al. 2018) showed that overall our final model fit the data reasonably well, but also revealed that goodness-of-fit varied substantially across the study period. Particularly the early years in the data, which correspond to the first two decades following dam construction, were characterized by relatively poorer model fit (Figures S4.3 & S4.7) Many individuals present during this period were hatched while the river was still free flowing and prior to implementation of the stocking programme. They may have experienced environmental conditions vastly different from individuals later in the time series, possibly resulting in long-lasting cohort effects not uncommon for salmonid fishes (e.g. Vincenzi et al. 2016). Furthermore, given the profound changes in harvest practices (gradual shift from gillnet to rod fishing, Aass and Kraabøl 1999), river regulation (flow regimes, turbine intake grid sizes, etc.), and disease prevalence during the 50-year study period, it is also not unlikely that size-dependence of mortality and migratory behavior itself has changed over time. Overdispersion in our data could thus be related to changes in selection pressures, something that may warrant attention in future studies.

Both parameter estimates and resulting model fit were sensitive to the way we imputed body size, illustrating that covariate imputation remains the main challenge of mark-recapture models with continuous individual time-varying covariates like body size (Pollock 2002, Bonner et al. 2010). Imputing body size using mean estimates from an externally run growth model, as we have done here, comes with several limitations. First, data used to estimate growth may not be representative of the individuals contained in the mark-recapture data. In our case, most data on growth in the lake pertains to the subadult life stage (prior to maturation) and resulting growth estimates may thus be less well suited for the mature, spawning trout that make up the mark-recapture data. Second, growth data is only available for 53% of individuals and 74% of years (only up to 2003) contained in the mark-recapture data. Size imputation for a non-random sample of individuals was thus lacking estimates of year and individual random effects. Finally, and per-haps most importantly, by directly imputing size using mean estimates of growth model parameters, we omitted all uncertainty in size estimates arising from residual variation in growth (stochasticity) and parameter uncertainty. Since the reduced growth model we used matched well with observations (Figure S5.1) and fit of the mark-recapture-recovery model was overall decent, it is unlikely that the results we present here are biased to a degree as to invalidate any of the main conclusions. However, as a result of direct size imputation and likely related lack of model fit, some of the patterns and effects may be estimated with inflated precision and this has to be considered when interpreting the presented relationships.

### OUTLOOK: DATA INTEGRATION AND POPULATION PERSPECTIVE

The fundamental issues arising from imputing missing individual covariate values can be addressed through integrated analysis of growth and survival/state transition processes (Bonner et al. 2010, Letcher et al. 2015), which allows imputation of the “true” latent body size and estimation of its effects on vital rates without bias and under full consideration of uncertainty. In our case, not just one but two distinct data sources provide information on growth: length measurements from trout captured in the fish ladder (mark-recapture data) and lengths back-calculated from scale year rings of a subset of marked individuals. This provides a unique opportunity for integrated analysis of multiple data sets which is likely to result in more precise estimates of vital rates, more comprehensive understanding of variation therein, and insights into potential discrepancies among different types of data (Plard et al. 2019b, Saunders et al. 2019).

The large drawback of Bayesian integrated analysis is its high computational costs, and in the case of the present data and model, computational demands precluded a fully integrated analysis. However, in SI S2 we have shown how implementing the mark-recapture-recovery model with a custom distribution in NIMBLE can lead to dramatic increases in computational efficiency (32-times faster MCMC than with standard JAGS). With the continuing development of both computational power and flexible, user-friendly MCMC software, large integrated analyses will likely become more feasible in the future.

More efficient computational solutions are also becoming invaluable when looking beyond single vital rates (growth, survival) and towards more holistic models of population dynamics. Several of the results presented here may have important implications for brown trout management but questions such as whether the high harvest mortality of adult trout has consequences for population via- bility or whether the dam does indeed function as an ecological trap, can only be addressed by adopting a population perspective. The framework of integrated population models (Plard et al. 2019b) in general, and recent extensions for populations structured by continuous traits in particular (Plard et al. 2019a), lend themselves well to the study of these questions for our system and will follow naturally from the integration of growth and survival estimation. Fully integrated, size-structured population models will further provide new opportunities to study the joint impacts of harvesting, stocking, habitat alteration, climate change, and disease dynamics (Plard et al. 2019b) and are thus highly relevant for future studies aiming to improve understanding and inform management of the trout in lake Mjøsa and of animal population inhabiting ecosystems heavily impacted by human activity in general.

## CONCLUSION

Multi-state mark-recapture models are powerful tools for estimating and understanding survival in animal populations that experience mortality from both natural and anthropogenic causes (Schaub and Pradel 2004). We used such a model to disentangle harvest- and background mortality of adult brown trout and showed that (1) harvest generally outweighed all other sources of mortality and (2) that vulnerability to both mortality causes was determined by individual differences in body size and migration pattern (dam passage). The use of a novel hazard rate parameterization (Ergon et al. 2018) and data from both recaptures and harvest recoveries allowed to estimate size-dependence and among-year variation in cause-specific mortality, state transition probabilities, and reporting rate without confounding. This framework, including the computationally efficient implementation of it, is highly applicable to other studies of cause-specific mortality in populations whose vital rates are strongly affected by continuous traits, and may prove particularly valuable also in the context of estimating correlation among different sources of mortality. Finally, we illustrated that the use of an appropriate year random effects structure can be a prerequisite to establishing identifiability of complex mark-recapture models and is therefore crucial to obtain reliable estimates of vital rate parameters. In practice, such random effects can only be estimated when data are collected over a sufficient number of years, emphasizing the importance of investing in the (continued) collection of individual-based data over long time periods (Clutton-Brock and Sheldon 2010).

## Supporting information

Full supporting information (S1-s6)

Maple code for intrinsic identifiability analyses

Code for custom model implementation with NIMBLE

## Acknowledgements

We thank David Koons and two anonymous reviewers for constructive feedback and suggestions during the review process. This work was supported by the Research Council of Norway (project SUSTAIN, 244647/E10). We thank all individuals and institutions that have been involved in collecting, maintaining, and organizing the trout data (contributions detailed in Moe et al. 2019), and the Norwegian Water and Energy Directorate (NVE) for providing river discharge data. Model fitting was performed on the Abel Cluster (University of Oslo and UNINETT Sigma2 AS).

## Authors’ contributions

CRN, TE, ØL, YV, and LAV conceived the ideas; CRN and TE designed methodology; CRN, PA, SJM, and AR prepared the data for analysis; CRN analysed the data and led the writing of the manuscript; DT developed and tested the custom likelihood and drafted SI S2. DC designed identifiability analyses and drafted SI S3. All authors contributed critically to the drafts and gave final approval for publication.

## Data accessibility

The complete mark-recapture-recovery and growth data sets will be made available on the Dryad Digital Repository (DOI to be added) and are documented in Moe et al. (2019).

## Supporting information

The following supporting information is available for this publication: Appendices S1 - S6.

## References

Aas, Ø., A. Klemetsen, S. Einum, and J. Skurdal, 2010. Atlantic salmon ecology. John Wiley & Sons.

Aass, P., 1993. Stocking strategy for the rehabilitation of a regulated brown trout (*Salmo trutta L*.) river. Regulated Rivers: Research & Management 8:135–144.

Aass, P. and M. Kraabøl, 1999. The exploitation of a migrating brown trout (*Salmo trutta L*.) population; change of fishing methods due to river regulation. River Research and Applications 15:211–219.

Aass, P., P. S. Nielsen, and Å. Brabrand, 1989. Effects of river regulation on the structure of a fast-growing brown trout (*Salmo trutta L*.) population. Regulated Rivers: Research & Management 3:255–266.

Arlinghaus, R., T. Klefoth, A. Kobler, and S. J. Cooke, 2008. Size selectivity, injury, handling time, and determinants of initial hooking mortality in recreational angling for northern pike: the influence of type and size of bait. North American Journal of Fisheries Management 28:123–134.

Bonner, S. J., B. J. Morgan, and R. King, 2010. Continuous covariates in mark-recapture-recovery analysis: a comparison of methods. Biometrics 66:1256–1265.

Carlin, B., 1955. Tagging of salmon smolts in the river lagan. *Rep. Inst Freshwat. Res.*, Drottningholm 36:57–74.

Carlson, S. M., E. M. Olsen, and L. A. Vøllestad, 2008. Seasonal mortality and the effect of body size: a review and an empirical test using individual data on brown trout. Functional Ecology 22:663–673.

Choquet, R. and D. J. Cole, 2012. A hybrid symbolic-numerical method for determinig model structure. Mathematical Biosciences 236:117–125.

Clutton-Brock, T. and B. C. Sheldon, 2010. Individuals and populations: the role of long-term, individual-based studies of animals in ecology and evolutionary biology. Trends in Ecology & Evolution 25:562–573.

Cole, D. J., 2012. Determining parameter redundancy of multi-state mark– recapture models for sea birds. Journal of Ornithology 152:305–315.

Cole, D. J., B. J. Morgan, and D. Titterington, 2010. Determining the parametric structure of models. Mathematical Biosciences 228:16–30.

Conn, P. B., D. S. Johnson, P. J. Williams, S. R. Melin, and M. B. Hooten, 2018. A guide to bayesian model checking for ecologists. Ecological Monographs 88:526– 542.

Cooch, E. G., M. Guillemain, G. S. Boomer, J.-D. Lebreton, and J. D. Nichols, 2014. The effects of harvest on waterfowl populations. Wildfowl pages 220–276.

Cox, D. R., 1972. Regression models and life-tables. Journal of the Royal Statistical Society: Series B (Methodological) 34:187–202.

De Roos, A. M., L. Persson, and E. McCauley, 2003. The influence of size-dependent life-history traits on the structure and dynamics of populations and communities. Ecology Letters 6:473–487.

de Valpine, P., D. Turek, C. J. Paciorek, C. Anderson-Bergman, D. T. Lang, and R. Bodik, 2017. Programming with models: writing statistical algorithms for general model structures with nimble. Journal of Computational and Graphical Statistics 26:403–413.

Dickinson, J. L., J. Shirk, D. Bonter, R. Bonney, R. L. Crain, J. Martin, T. Phillips, and K. Purcell, 2012. The current state of citizen science as a tool for ecological research and public engagement. Frontiers in Ecology and the Environment 10:291–297.

Drenner, S. M., T. D. Clark, C. K. Whitney, E. G. Martins, S. J. Cooke, and S. G. Hinch, 2012. A synthesis of tagging studies examining the behaviour and survival of anadromous salmonids in marine environments. PloS One 7:e31311.

Ellner, S. P. and M. Rees, 2006. Integral projection models for species with complex demography. The American Naturalist 167:410–428.

Ergon, T., Ø. Borgan, C. R. Nater, and Y. Vindenes, 2018. The utility of mortality hazard rates in population analyses. Methods in Ecology and Evolution 9:2046– 2056.

Fernández-Chacón, A., E. Moland, S. H. Espeland, A. R. Kleiven, and E. M. Olsen, 2017. Causes of mortality in depleted populations of atlantic cod estimated from multi-event modelling of mark–recapture and recovery data. Canadian Journal of Fisheries and Aquatic Sciences 74:116–126.

Fernández-Chacón, A., E. Moland, S. H. Espeland, and E. M. Olsen, 2015. Demographic effects of full vs. partial protection from harvesting: inference from an empirical before–after control-impact study on atlantic cod. Journal of Applied Ecology 52:1206–1215.

Fjeldstad, H.-P., U. Pulg, and T. Forseth, 2018. Safe two-way migration for salmonids and eel past hydropower structures in Europe: a review and recommendations for best-practice solutions. Marine and Freshwater Research 69:1834–1847.

Garrett, E. S. and S. L. Zeger, 2000. Latent class model diagnosis. Biometrics 56:1055–1067.

Gimenez, O., R. Choquet, and J.-D. Lebreton, 2003. Parameter redundancy in multistate capture-recapture models. Biometrical Journal: Journal of Mathematical Methods in Biosciences 45:704–722.

Gimenez, O., B. J. Morgan, and S. P. Brooks, 2009. Weak identifiability in models for mark-recapture-recovery data. In Modeling demographic processes in marked populations, pages 1055–1067. Springer.

Hamley, J. M., 1975. Review of gillnet selectivity. Journal of the Fisheries Board of Canada 32:1943–1969.

Haugen, T. O., P. Aass, N. C. Stenseth, and L. A. Vøllestad, 2008. Changes in selection and evolutionary responses in migratory brown trout following the construction of a fish ladder. Evolutionary Applications 1:319–335.

Hilborn, R. and C. J. Walters, 2013. Quantitative fisheries stock assessment: choice, dynamics and uncertainty. Springer Science & Business Media.

Hobæk, A., J. E. Løvik, T. Rohrlack, S. J. Moe, M. Grung, H. Bennion, G. Clarke, and G. T. Piliposyan, 2012. Eutrophication, recovery and temperature in lake Mjøsa: detecting trends with monitoring data and sediment records. Freshwater Biology 57:1998–2014.

Hudson, P. J., A. Rizzoli, B. T. Grenfell, J. Heesterbeek, and A. P. Dobson, 2002. Ecology of wildlife diseases.

Johnsen, B. U. and O. Ugedal, 2001. Soppinfeksjoner (*Saprolegnia* spp.) på lakse-fisk i norge - statusrapport. NINA Oppdragsmelding 716:1–34.

Jonsson, B. and N. Jonsson, 2011. Maturation and Spawning. In *Ecology of Atlantic Salmon and Brown Trout*, pages 327–414. Springer Netherlands, Dordrecht.

Kleiven, A. R., A. Fernandez-Chacon, J.-H. Nordahl, E. Moland, S. H. Espeland, H. Knutsen, and E. M. Olsen, 2016. Harvest pressure on coastal Atlantic cod (*Gadus morhua*) from recreational fishing relative to commercial fishing assessed from tag-recovery data. PLoS One 11:e0149595.

Koons, D. N., R. F. Rockwell, and L. M. Aubry, 2014. Effects of exploitation on an overabundant species: the lesser snow goose predicament. Journal of Animal Ecology 83:365–374.

Kraabøl, M., 2006. Gytebiologi hos Hunderørret i Gudbrandsdalslågen nedenfor Hunderfossen kraftverk. NINA rapport 217 pages 1–34.

Lebreton, J.-D., T. Almeras, and R. Pradel, 1999. Competing events, mixtures of information and multistratum recapture models. Bird Study 46:S39–S46.

Lebreton, J.-D. and R. Pradel, 2002. Multistate recapture models: modelling in-complete individual histories. Journal of Applied Statistics 29:353–369.

Lennox, R. J., J. Alós, R. Arlinghaus, A. Horodysky, T. Klefoth, C. T. Monk, and S. J. Cooke, 2017. What makes fish vulnerable to capture by hooks? a conceptual framework and a review of key determinants. Fish and Fisheries 18:986–1010.

Letcher, B. H., P. Schueller, R. D. Bassar, K. H. Nislow, J. A. Coombs, K. Sakrejda, M. Morrissey, D. B. Sigourney, A. R. Whiteley, M. J. O’donnell, et al., 2015. Robust estimates of environmental effects on population vital rates: an integrated capture–recapture model of seasonal brook trout growth, survival and movement in a stream network. Journal of Animal Ecology 84:337–352.

Lewin, W.-C., R. Arlinghaus, and T. Mehner, 2006. Documented and potential biological impacts of recreational fishing: insights for management and conservation. Reviews in Fisheries Science 14:305–367.

Moe, S. J., C. R. Nater, A. Rustadbakken, L. A. Vøllestad, E. Lund, T. Qve-nild, O. Hegge, and P. Aass, 2019. A 50-year series of mark-recapture data of large-sized brown trout (*Salmo trutta*) from Lake Mjøsa, Norway. bioRxiv doi:10.1101/544825.

Nater, C. R., A. Rustadbakken, T. Ergon, Ø. Langangen, S. J. Moe, Y. Vindenes, L. A. Vøllestad, and P. Aass, 2018. Individual heterogeneity and early life conditions shape growth in a freshwater top predator. Ecology 99:1011–1017.

NIMBLE Development Team, 2018. NIMBLE: MCMC, particle filtering, and programmable hierarchical modeling. R package version 0.6–13.

Okamura, B. and S. W. Feist, 2011. Emerging diseases in freshwater systems. Freshwater Biology 56:627–637.

Piccolo, J. J., J. R. Norrgård, L. A. Greenberg, M. Schmitz, and E. Bergman, 2012. Conservation of endemic landlocked salmonids in regulated rivers: a case-study from lake Vänern, Sweden. Fish and Fisheries 13:418–433.

Plard, F., R. Fay, M. Kéry, A. Cohas, and M. Schaub, 2019b. Integrated population models: powerful methods to embed individual processes in population dynamics models. Ecology page e02715.

Plard, F., D. Turek, M. U. Grüebler, and M. Schaub, 2019a. IPM^2^: Toward better understanding and forecasting of population dynamics. Ecological Monographs 89:e01364.

Pollock, K. H., 2002. The use of auxiliary variables in capture-recapture modelling: an overview. Journal of Applied Statistics 29:85–102.

Quinn, T. J., 2003. Ruminations on the development and future of population dynamics models in fisheries. Natural Resource Modeling 16:341–392.

R Core Team, 2018. R: A Language and Environment for Statistical Computing. R Foundation for Statistical Computing, Vienna, Austria.

Richards, S. A., 2008. Dealing with overdispersed count data in applied ecology. Journal of Applied Ecology 45:218–227.

Ronget, V., M. Garratt, J.-F. Lemaître, and J.-M. Gaillard, 2017. The ‘Evo-Demo’ implications of condition-dependent mortality. Trends in Ecology & Evolution 32:909–921.

Roscoe, D., S. Hinch, S. Cooke, and D. Patterson, 2011. Fishway passage and post-passage mortality of up-river migrating sockeye salmon in the Seton River, British Columbia. River Research and Applications 27:693–705.

Sæther, B.-E. and Ø. Bakke, 2000. Avian life history variation and contribution of demographic traits to the population growth rate. Ecology 81:642–653.

Saunders, S. P., M. T. Farr, A. D. Wright, C. A. Bahlai, J. W. Ribeiro Jr, S. Rossman, A. L. Sussman, T. W. Arnold, and E. F. Zipkin, 2019. Disentangling data discrepancies with integrated population models. Ecology page e02714.

Schaub, M. and R. Pradel, 2004. Assessing the relative importance of different sources of mortality from recoveries of marked animals. Ecology 85:930–938.

Schlaepfer, M. A., M. C. Runge, and P. W. Sherman, 2002. Ecological and evolutionary traps. Trends in Ecology & Evolution 17:474–480.

Servanty, S., R. Choquet, É. Baubet, S. Brandt, J.-M. Gaillard, M. Schaub, C. Toïgo, J.-D. Lebreton, M. Buoro, and O. Gimenez, 2010. Assessing whether mortality is additive using marked animals: a Bayesian state–space modeling approach. Ecology 91:1916–1923.

Turek, D., P. de Valpine, and C. J. Paciorek, 2016. Efficient Markov chain Monte Carlo sampling for hierarchical hidden Markov models. Environmental and Ecological Statistics 23:549–564.

Vincenzi, S., M. Mangel, D. Jesensek, J. C. Garza, and A. J. Crivelli, 2016. Within- and among-population variation in vital rates and population dynamics in a variable environment. Ecological Applications 26:2086–2102.

Vindenes, Y. and Ø. Langangen, 2015. Individual heterogeneity in life histories and eco-evolutionary dynamics. Ecology Letters 18:417–432.

Williams, B. K., J. D. Nichols, and M. J. Conroy, 2002. Analysis and management of animal populations. Academic Press.

Zhao, Q., G. S. Boomer, and W. L. Kendall, 2018. The non-linear, interactive effects of population density and climate drive the geographical patterns of waterfowl survival. Biological Conservation 221:1–9.

