## Supplementary material for "Size- and stage-dependence in cause-specific mortality of migratory brown trout": Full supporting information (S1-s6)

**S6 Extended Model with Correlated Random Effects** **44**

*

#### S1 Supplementary Figures and Tables

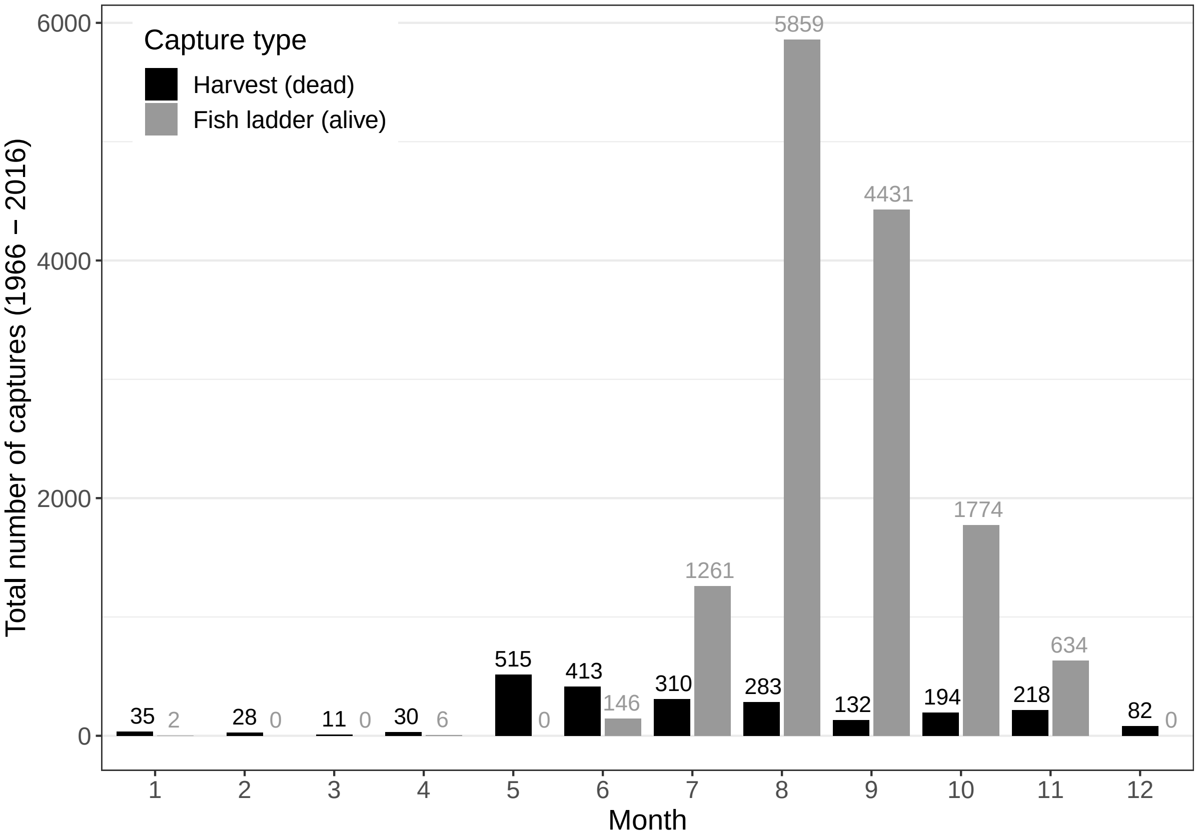

Figure S1.1: Distribution of trout captures over calendar months for the time period 1966 - 2016. Grey = trout captured alive in the fish ladder (marking and recapture events), black = trout captured and reported dead by fishers.

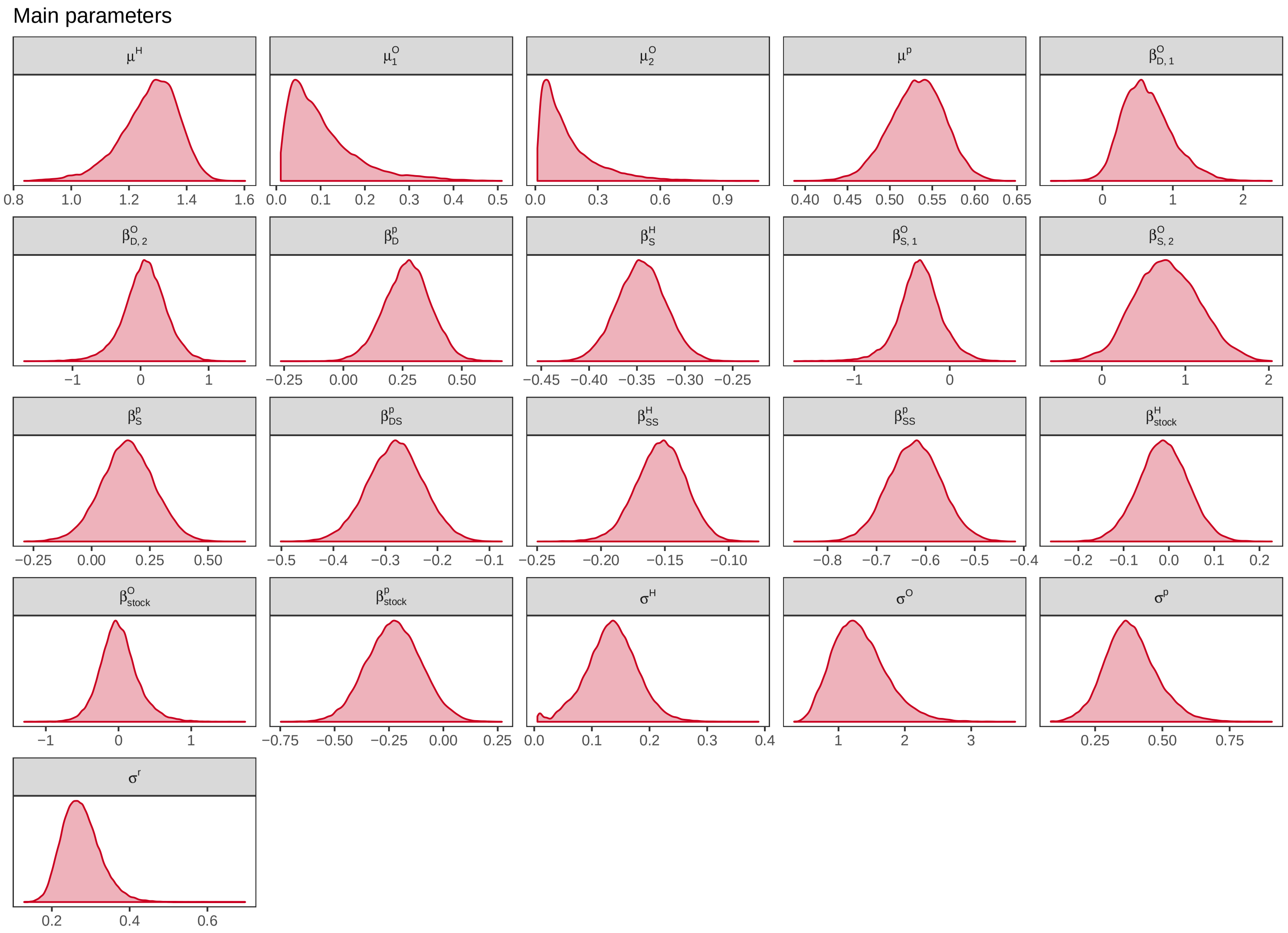

Figure S1.2: Posterior distributions for intercept, slope, and variance parameters.

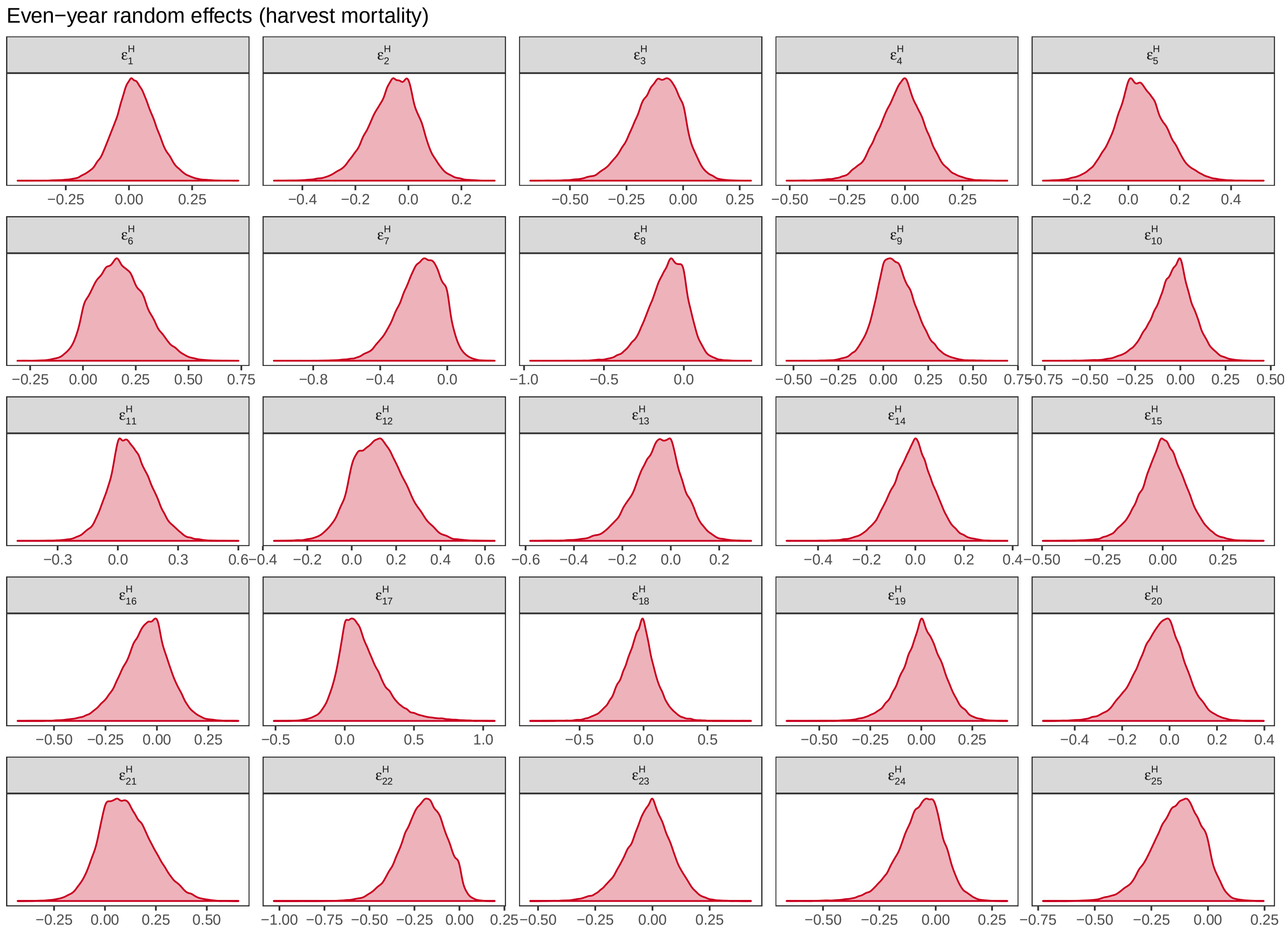

Figure S1.3: Posterior distributions for even-year harvest mortality random effect levels.

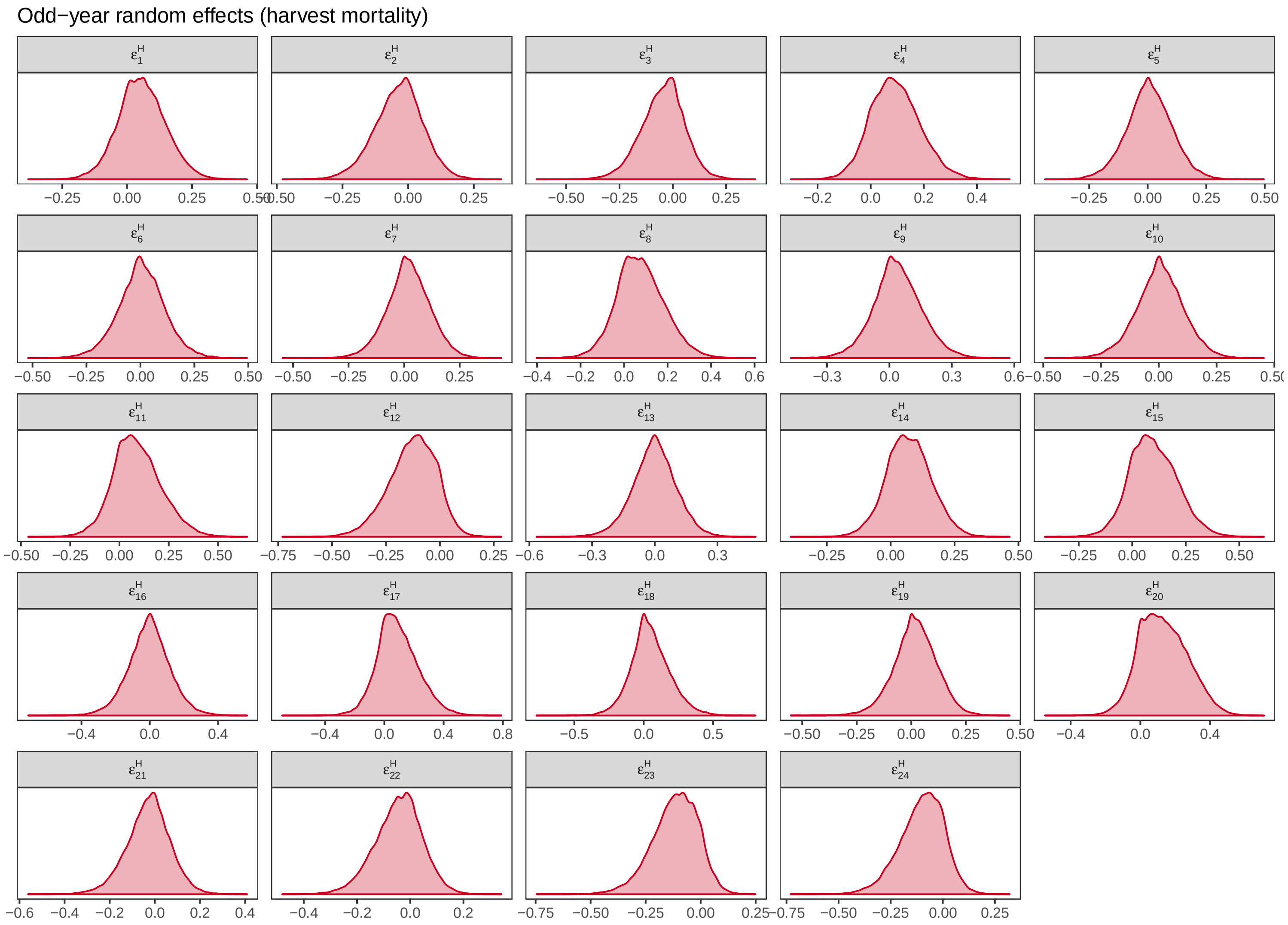

Figure S1.4: Posterior distributions for odd-year harvest mortality random effect levels.

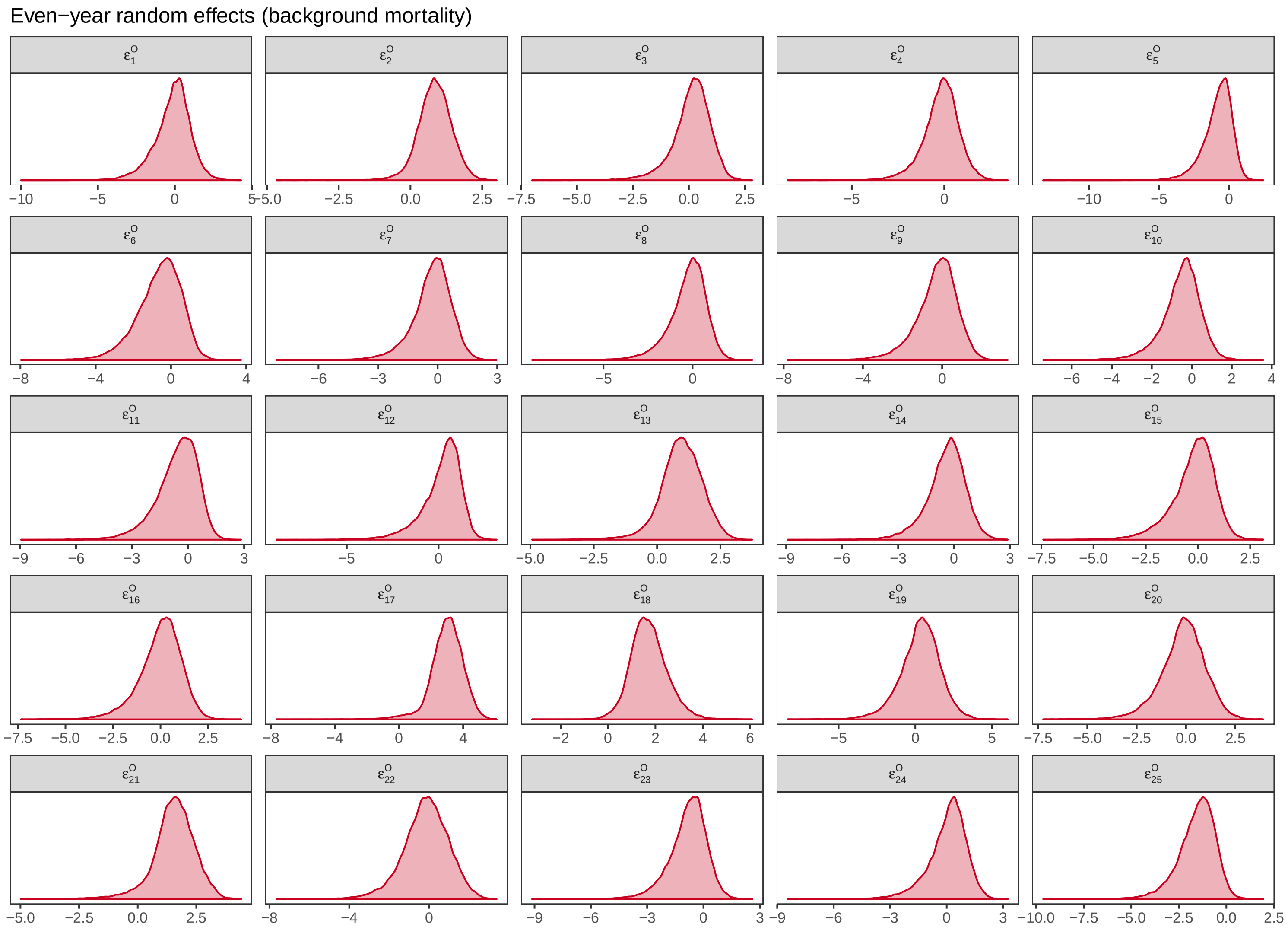

Figure S1.5: Posterior distributions for even-year background mortality random effect levels.

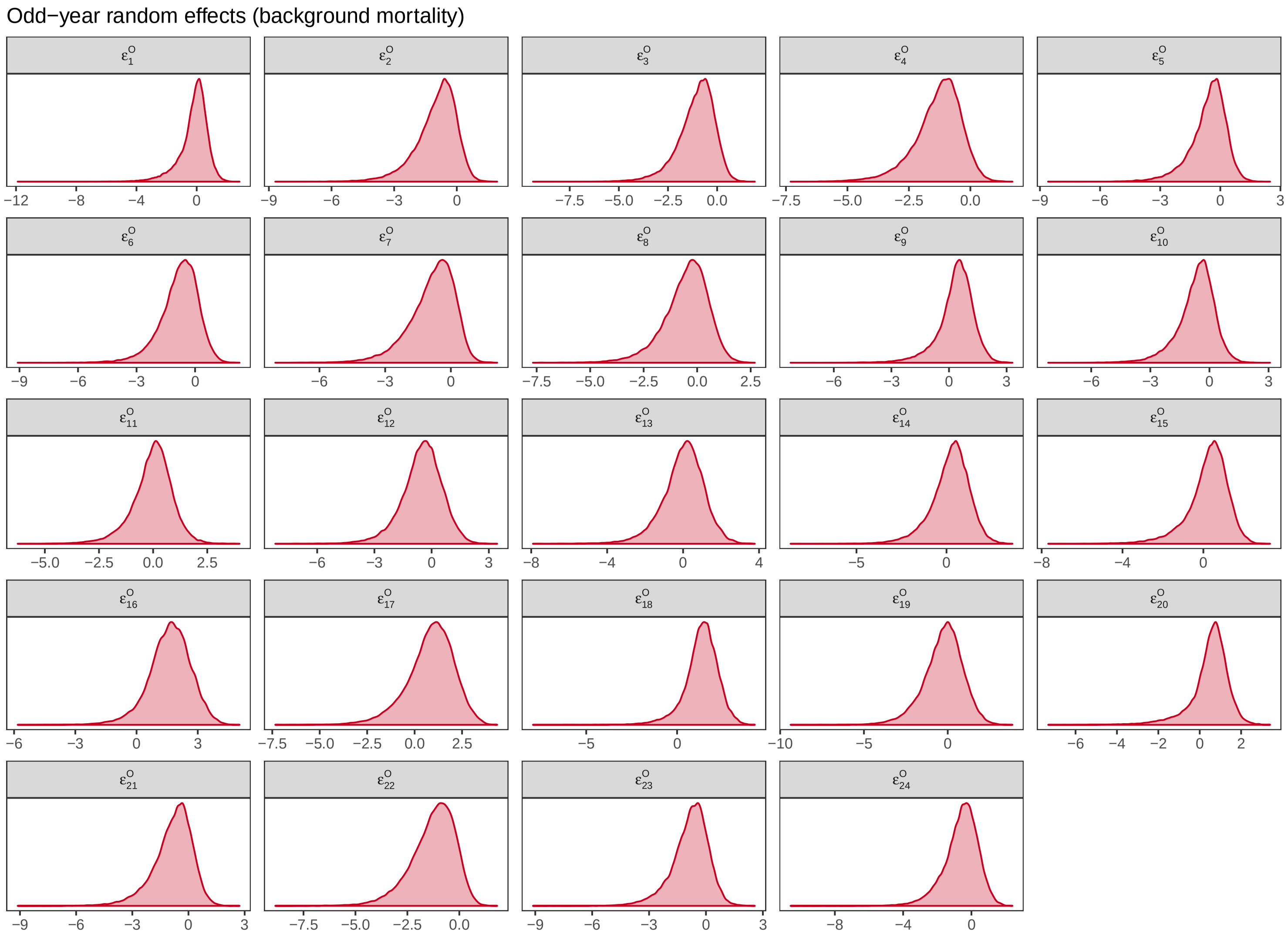

Figure S1.6: Posterior distributions for odd-year background mortality random effect levels.

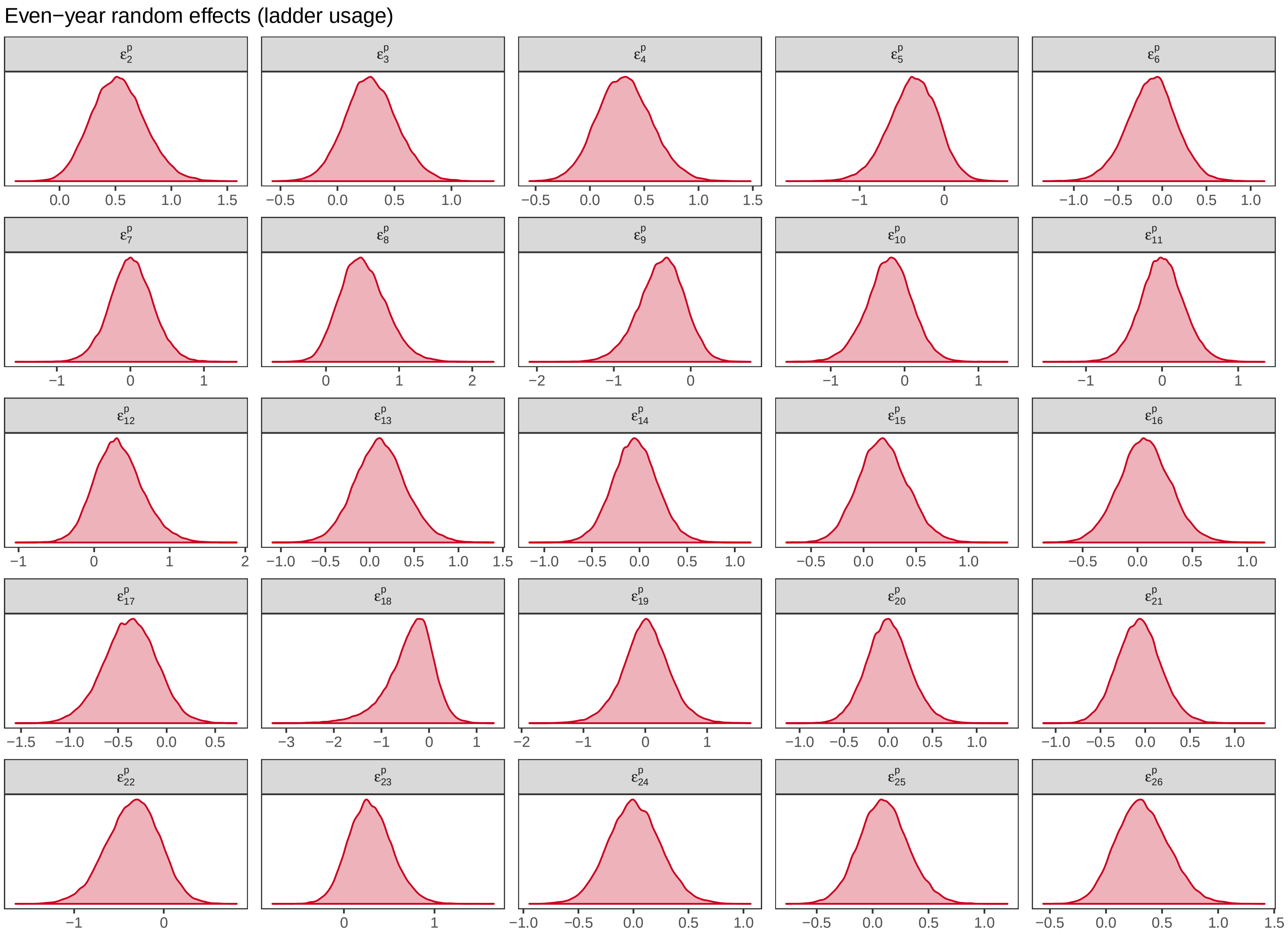

Figure S1.7: Posterior distributions for even-year ladder usage probability random effect levels.

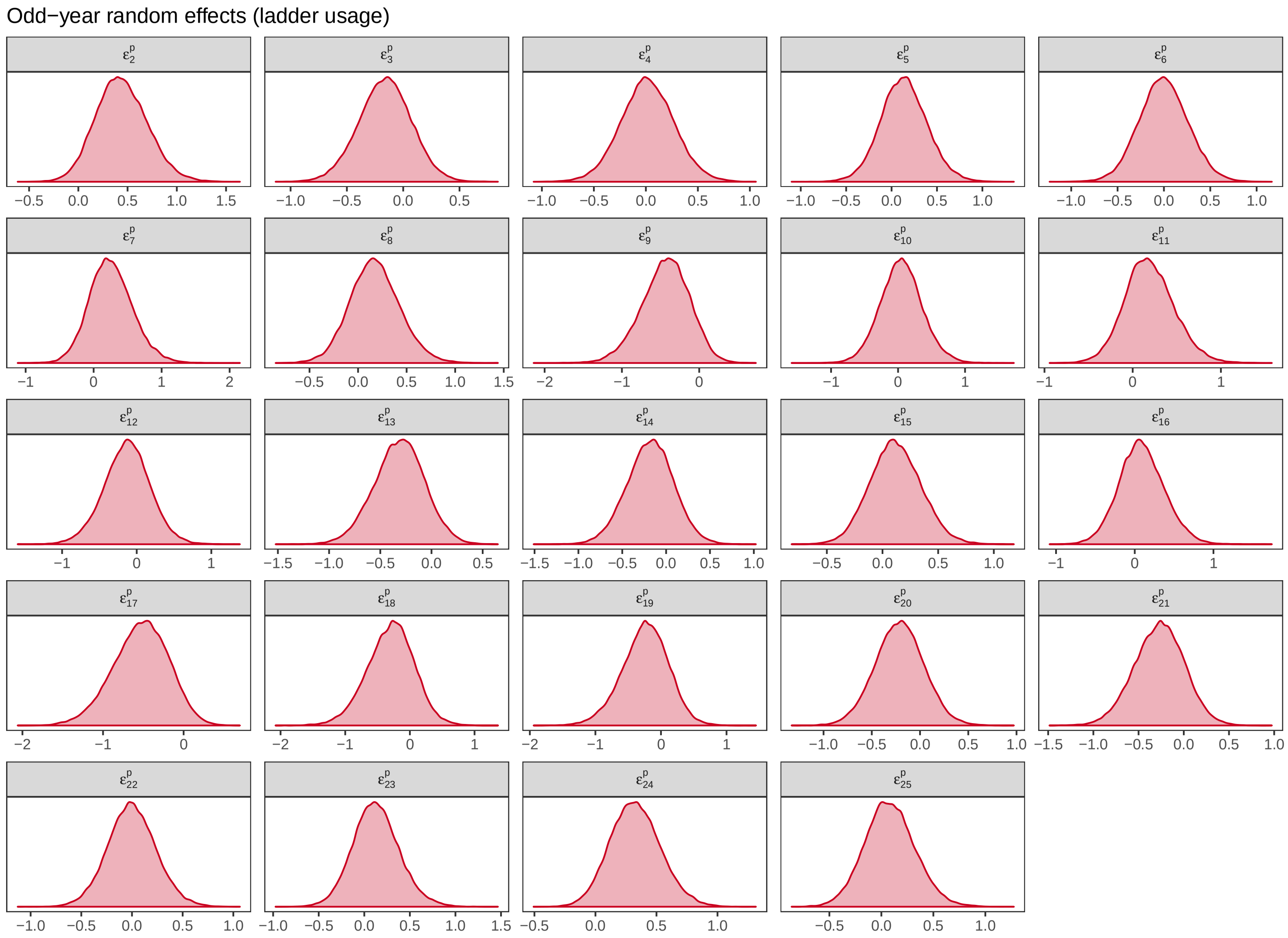

Figure S1.8: Posterior distributions for odd-year ladder usage probability random effect levels estimated from real data.

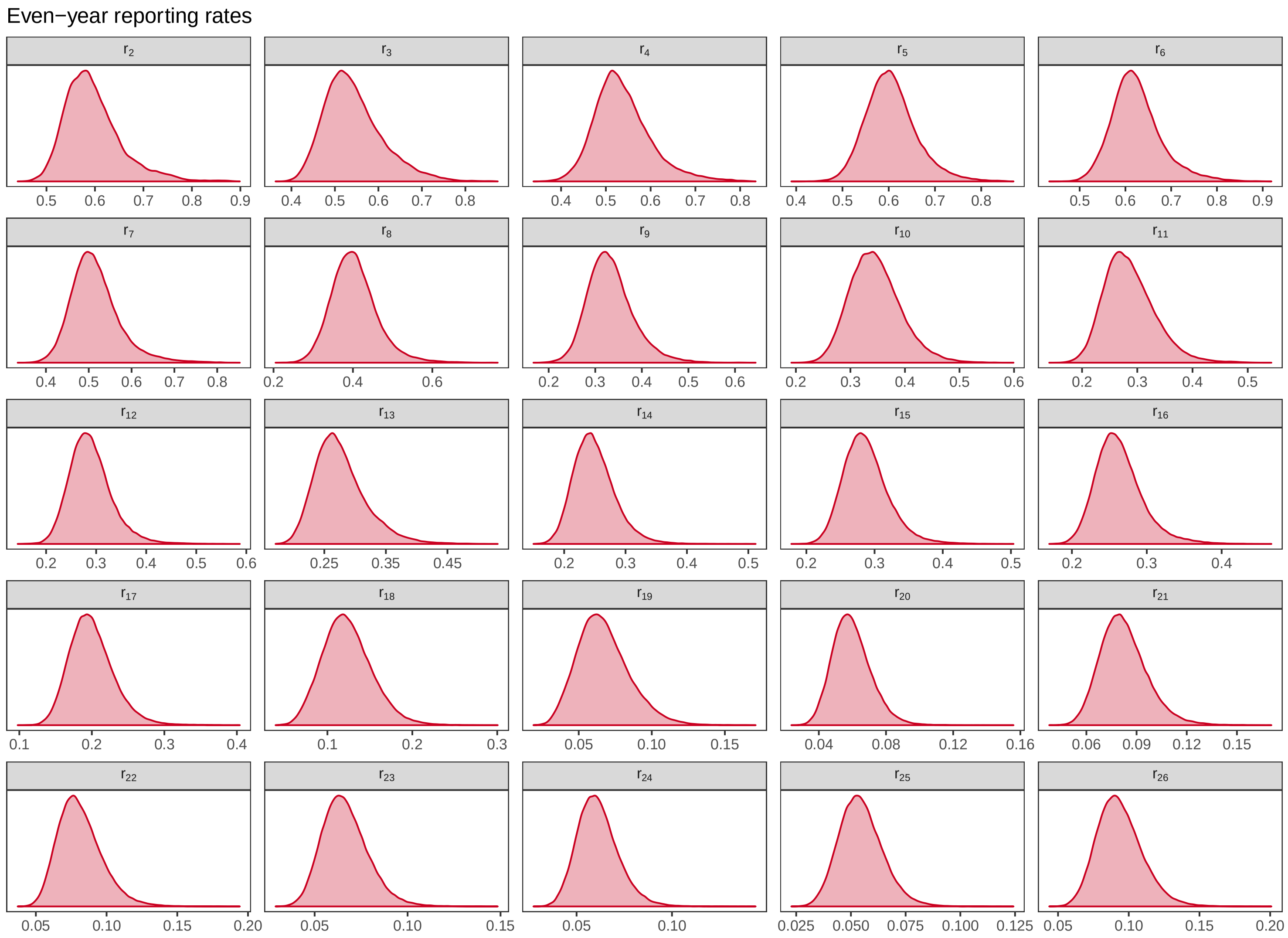

Figure S1.9: Posterior distributions for even-year reporting rates.

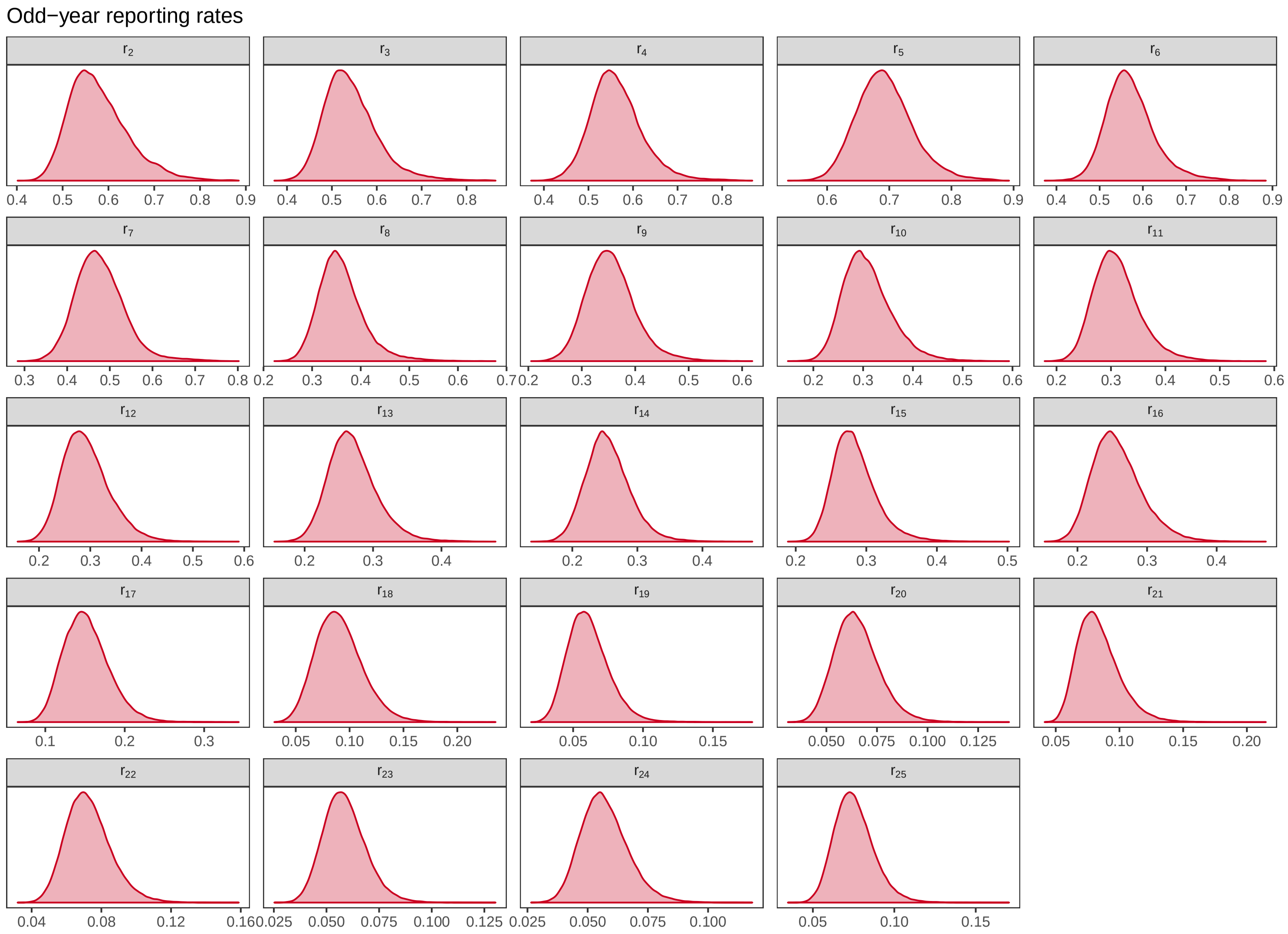

Figure S1.10: Posterior distributions for odd-year reporting rates.

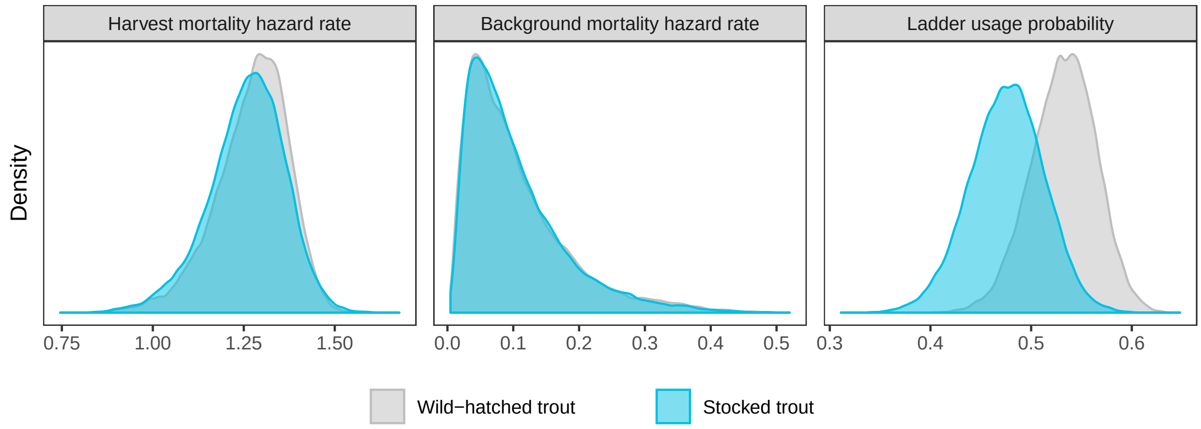

Figure S1.11: Posterior distributions of baseline harvest / background mortality hazard rates (above-dam spawners) and ladder usage probabilities of wild-hatched (grey) and stocked (blue) trout.

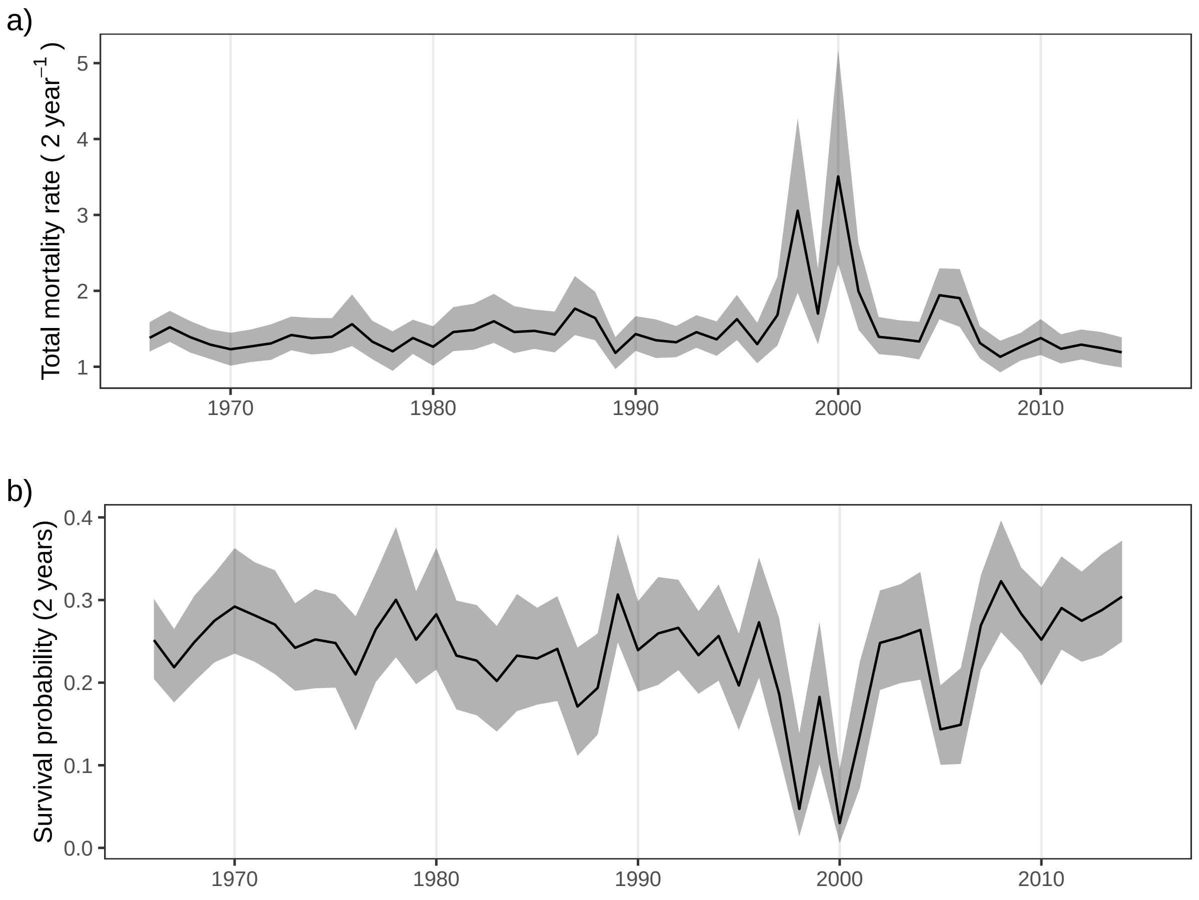

Figure S1.12: Time variation in a) survival probability and b) total mortality hazard rate of above-dam spawners. Lines represent the median predictions, ribbons indicate 95% credibility intervals.

#### S2 Hierarchical Model using Custom Likelihood Distribution

We used the `nimble` package (de Valpine et al. 2017, NIMBLE Development Team 2018) for R for specifying the hierarchical model, and model fitting via MCMC. NIMBLE provides a great deal of algorithmic and modeling flexibility, including a possibility to define custom distributions for use in the hierarchical model specifications. Here, we take advantage of this possibility to greatly reduce the the prohibitively long MCMC runtimes of analysis of our large mark-recapture-recovery dataset (which can not be summarized into reduced data representations such as "m-arrays" due to the individual size covariate in the model).

In particular, we define a custom distribution `dtroutDHMM`, the name suggesting its use in a *discrete hidden Markov model*, to exactly calculate the likelihood of each capture history, conditional on values of the model parameters. Doing so allows removing a over 6,000 latent states (representing the true state of individuals at each sampling occasion) from the hierarchical model, and thus reduces the dimension of the posterior distribution (and equivalently the MCMC sampling problem) by that same number. This also serves to improve the MCMC mixing of the remaining posterior dimensions, as it no longer relies on MCMC integration over the nuisance dimensions. Specifically, for model parameters $\theta$ and capture histories $y = \{y_1, \ldots, y_n\}$, the posterior distribution is updated according to:

$$p(\theta|y) \propto p(\theta) \prod_{i=1}^{n} p(y_i|\theta),$$

where the likelihood $p(y_i|\theta)$ of capture history $y_i$ is calculated using the custom likelihood function, and $p(\theta)$ is the prior specification. Our implementation extends that of Turek et al. (2016) by incorporating individual-specific covariates (in this case, body length) into the likelihood calculation. In addition, to further speed up computation time, our custom implementation strictly uses one-dimensional linear calculations in lieu of the matrix operations used in Turek et al. (2016). This forgoes the need to construct multi-dimensional arrays for storing state transition and observation probabilities, which were found to be prohibitively large.

The reductions in the model structure made by the `dtroutDHMM` distribution come with added computational expense of the exact likelihood calculation, which involves a discrete summation over the possible latent state values. But as we see, this added computation is dwarfed by the gains realized from smaller memory demands and improved mixing of the resulting MCMC.

In this section, we first present the gains realized through use of this custom distribution, and then provide the definition and details of the custom `dtroutDHMM` distribution.

#### S2.1 Comparison of Hierarchical Model Performance

We first present performance comparisons between three different implementations of the hierarchical model. The three implementations differ in the underlying software and representation of the model, but are identical in their inferences for the biological parameters of interest.

The first model, denoted "jags LS" is a latent state formulation of the model specified using the `jags` package (Plummer 2003) for R. This formulation contains over 60,000 latent variables representing the true, unknown state of each individual on each relevant time period.

The second model, denoted "nimble LS", is a porting of this same latent state representation into NIMBLE, a more flexible and faster platform for hierarchical models and MCMC.

The third and final version, "nimble DHMM", is constructed using NIMBLE, *and* makes use of the custom `dtroutDHMM` distribution to directly calculate model likelihoods, thus removing the over 60,000 latent states from the model. This version of the model leaves only the top-level model parameters (intercept, slope, and variance parameters) to undergo MCMC sampling. We note that this represents a 99.9% reduction in the dimension of the MCMC sampling problem, albeit incurring additional computational costs for the exact evaluation of the model likelihood.

##### MCMC sampling efficiency

Use of the custom `dtroutDHMM` distribution dramatically speeds up MCMC mixing of the remaining model parameters. This is because the mixing of the top-level parameters is no longer hampered by potentially slow mixing of the massive number of latent states. Removing the latent states from the model, the top-level parameters are able to mix freely, as we are "only" exploring the $n$-dimensional space of the posterior distribution (where $n$ = number of top parameters).

As discussed, these gains in mixing come at the cost of a higher computational burden of the likelihood evaluations. Thus, our measure of MCMC "efficiency" takes into account both MCMC mixing as well as the associated computational requirement. We define the efficiency of MCMC sampling as the *rate of generating effectively independent posterior samples* (Turek et al. 2017). Thus, efficiency is calculated as the posterior effective sample size (ESS) divided by the MCMC runtime (measured in seconds), and has units of effectively independent posterior samples per second.

This efficiency can be calculated independently for each posterior dimension. We generally consider an MCMC to only be as good as its slowest mixing dimension, and thus consider the minimum efficiency across all model parameters. That said, the mean efficiency (across model parameters) is also an interesting metric to consider, as well as the distribution of sampling efficiencies across all model parameters.

Figure S2.1 compares the sampling efficiencies across all top-level parameters for each model formulation (run on simulated data[1]). Focusing on the minimum efficiency across parameters, we see a 6-fold increase using the "nimble LS" model as compared to jags, and a 32-fold increase through use of the custom distribution "nimble DHMM" formulation of the model.

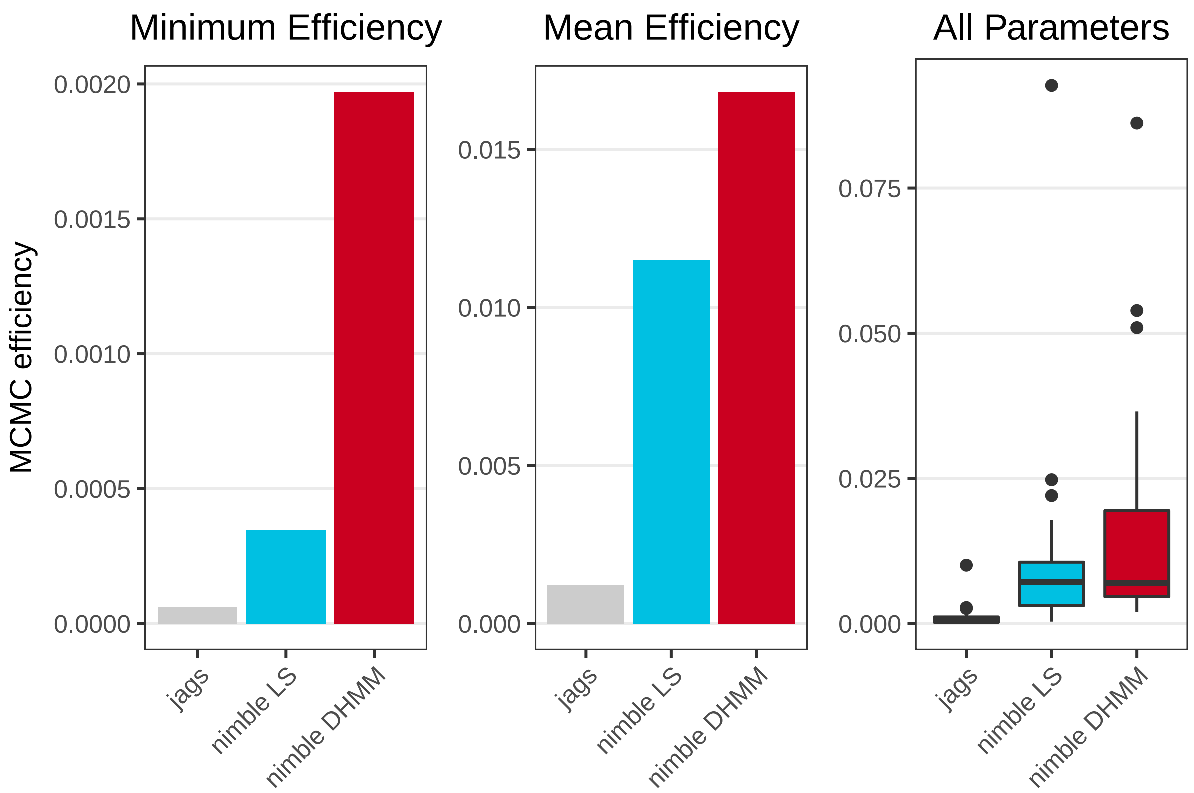

Figure S2.1: Sampling efficiencies for top-level model parameters resulting from the three different model formulations (run on simulated data). Minimum efficiency across all model parameters (left), mean efficiency across model parameters (center), and boxplots showing the distribution of efficiency values for all parameters (right).

[1] We simulated capture histories assuming similar top-level parameter values as obtained from model runs on real data and ignoring effects of hatchery origin. We further used a simpler model for reporting rate in the simulations. This model assumed five time periods, in each of which $r$ was allowed to vary around a period-specific mean: $logit(r_t) = logit(\mu^r_{period}) + \epsilon^r_t$. We chose this model for testing the custom distribution for convenience, because runtimes are considerably longer when using the autocorrelated reporting rate model instead.

As an alternate view of the same data, Figure S2.2 separates the model parameters into natural groupings of structurally-similar parameters, and presents the progression of sampling efficiency of each parameter resulting from each model formulation.

We universally observe increases in sampling efficiency through use of NIMBLE and the custom `dtroutDHMM` distribution, the improvement of the latter being most pronounced for the covariate effects on mortality and ladder usage.

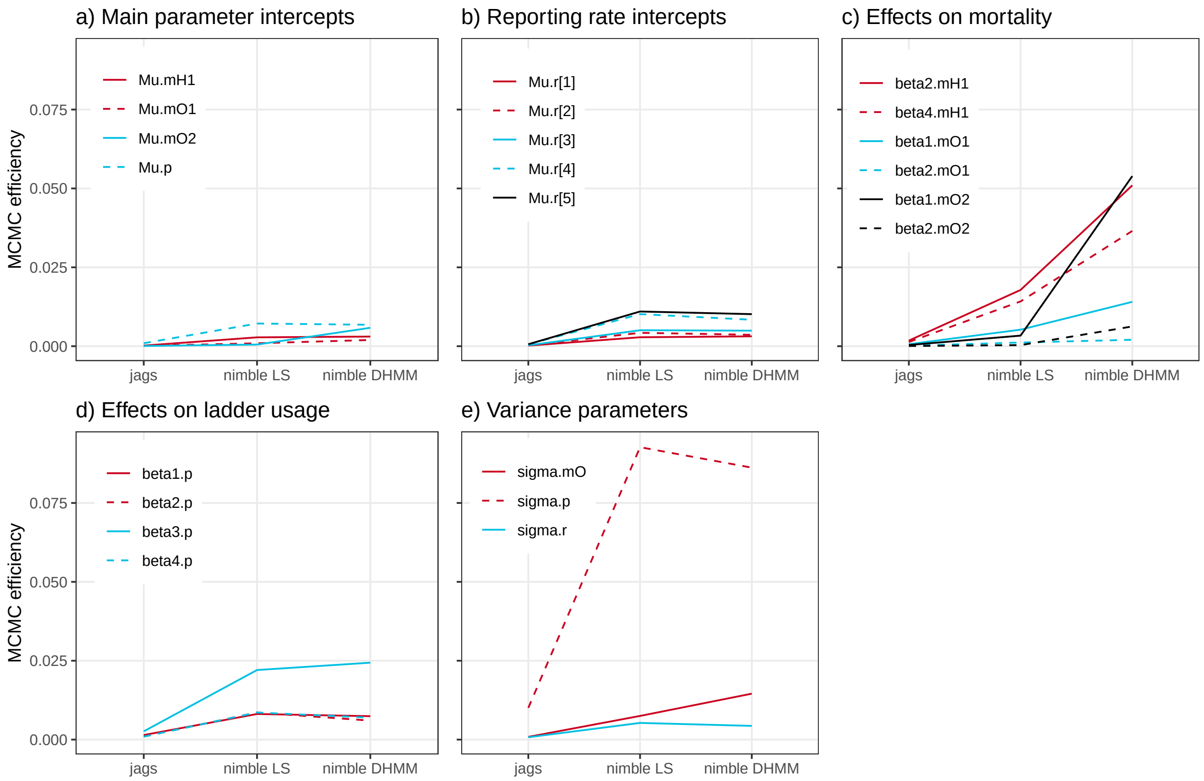

Figure S2.2: MCMC sampling efficiencies for model parameters under each model formulation (run on simulated data), grouped by the structural nature of the parameters. Parameters are named as written in code and defined in section S2.2. There are five period-specific mean reporting rate parameters resulting from the use of an alternative reporting rate model for simulating data and testing MCMC efficiency[1].

#### Model build and compilation time

Unlike jags, NIMBLE separates the steps of (a) building the hierarchical model object, (b) compilation to C++, and (c) MCMC execution. For the comparisons of MCMC efficiency presented above, we only considered the final step of MCMC execution. However, for the NIMBLE versions of the model we can also monitor the model build and compilation times, and report the additional savings form using the custom `dtroutDHMM` distribution form of the model.

Broadly, “model building” consists of parsing and processing the BUGS code representation of the model, constructing an equivalent directed acyclic graph object with functions for node density calculations and random simulation, and instantiating and populating data structures for storing the values of model parameters, latent states, and data. The "compilation" step consists of dynamically generating equivalent C++ code to represent the functions and data structures within the model and MCMC, invoking a C++ compiler to (on-the-fly) compile this dynamically generated code, linking this compiled library into the current R session, and instantiating compiled objects using the linked library. For large models, as here, these steps are non-trivial.

Table S2.1 presents the wall-clock time requirements for model building and compilation, using the two different NIMBLE formulations of the model: the "nimble LS" model including latent states, and the "nimble DHMM" model making use of the custom `dtroutDHMM` distribution. Through use of the custom distribution DHMM model, we observe a 370-fold reduction in model building time, and a 12-fold reduction in compilation time. These savings are noteworthy, as this represents a reduction by more than 7 hours in terms of actual wall-clock time.

Table S2.1: Absolute and relative amounts of time required by `nimble` to build (with and without checking calculations) and compile the model with either the latent state or custom hidden Markov distribution (DHMM) formulation.

| | **Latent State** | | **DHMM** | | **Fold-increase with DHMM** |
|---|---|---|---|---|---|
| | hours | minutes | hours | minutes | |
| **Model Building** | 7.252 | 435.131 | 0.020 | 1.178 | 369.519 |
| **Compilation** | 0.187 | 11.237 | 0.016 | 0.946 | 11.879 |

##### Memory usage

Finally, use of the custom `dtroutDHMM` distribution also provides a huge reduction in the memory demands of the MCMC. This is a result of the large multi-dimensional data structures required by the latent state formulation of the model, which are not needed or constructed in the DHMM formulation.

The maximal memory usage during the latent state MCMC reached 13.17 GB. In contrast, the MCMC using the DHMM formulation of the model had a maximal memory usage of 1.23 GB, representing a 93% reduction in (maximal) memory usage. Most importantly, this reduction brings the MCMC sampling problem into the realm of tractability for many local computing platforms, as 13 GB of RAM is in excess of the memory available on many laptop or desktop computers.

All analyses presented in this section were run under OSX (version 10.14.2), using a 3.3 GHz Intel Core i5 processor, and with 32 GB of memory.

#### S2.2 Specification of Custom `dtroutDHMM` Likelihood Distribution

Now we present the definition of the custom `dtroutDHMM` distribution, as used in the "nimble DHMM" formulation of the model.

This distribution is defined as a `nimbleFunction`, following particular conventions to allow the distribution to be used in NIMBLE model code, and to pass through the NIMBLE compiler. See the NIMBLE User Manual[2] for further details of writing custom distributions and other `nimbleFunction`s.

Following the `nimbleFunction` definition of the custom `dtroutDHMM` distribution, we provide an itemized description of the distribution's arguments, and intermediate variables used in the calculation of the model likelihood.

---

```
dtroutDHMM <- nimble::nimbleFunction(
    run = function(
        ## argument type declarations
        x = double(1), length = double(), origin = double(),
        mu.mH1 = double(), mu.m01 = double(), mu.m02 = double(), mu.p = double(),
        beta1.m01 = double(), beta1.m02 = double(), beta2.m01 = double(), beta2.m02 = double(),
        betaS.m0 = double(), beta2.mH1 = double(), beta4.mH1 = double(), betaS.mH = double(),
        beta1.p = double(), beta2.p = double(),
        beta3.p = double(), beta4.p = double(), betaS.p = double(), size = double(1),
        epsilon.mH = double(1), epsilon.m0 = double(1), epsilon.p = double(1),
        discF = double(1), discS = double(1), r = double(1), log = double()) {

        ## calculate background and harvest mortality hazard rates,
        ## ladder usage probability, and survival probabilities
        mH1 <- exp(mu.mH1 + betaS.mH*origin + beta2.mH1*size + beta4.mH1*size^2 + epsilon.mH)
        m01 <- exp(mu.m01 + betaS.m0*origin + beta1.m01*discF + beta2.m01*size + epsilon.m0)
        m02 <- exp(mu.m02 + betaS.m0*origin + beta1.m02*discF + beta2.m02*size + epsilon.m0)
        p <- expit(mu.p + betaS.p*origin + beta1.p*discS + beta2.p*size + beta3.p*discS*size +
        beta4.p*size^2 + epsilon.p)
        S1 <- exp(-(mH1 + m01))
        alpha1 <- mH1 / (mH1 + m01)
        S2 <- exp(-(mH1 + m02))
        alpha2 <- mH1 / (mH1 + m02)

        logL <- 0                              ## initialize log-likelihood
        pi <- numeric(4, init = FALSE)
        pi[1] <- S1[1]*p[2]                    ## calculate probability of each
        pi[2] <- S1[1]*(1-p[2])                ## unobserved state, conditional
        pi[3] <- (1-S1[1])*alpha1[1]           ## on first observation of spawning
        pi[4] <- (1-S1[1])*(1-alpha1[1])
        for(t in 2:length) {                   ## iterate over remaining observations
            Zpi <- pi
            if(x[t] == 1) { Zpi[2] <- 0        ## observed and seen alive:
                            Zpi[3] <- 0        ## update conditional distribution of unobserved
                            Zpi[4] <- 0 }      ## states, given this observation
            if(x[t] == 2) { Zpi[1] <- 0        ## harvested and reported:
```

[2] https://r-nimble.org/html_manual/cha-welcome-nimble.html

```
                        Zpi[2] <- 0       ## update conditional distribution
                        Zpi[3] <- pi[3] * r[t]
                        Zpi[4] <- 0 }
        if(x[t] == 3) { Zpi[1] <- 0       ## not seen or reported
                        Zpi[3] <- pi[3] * (1-r[t]) }
        sumZpi <- sum(Zpi)
        logL <- logL + log(sumZpi)        ## log-likelihood contribution of observed state
        if(t != length) {                 ## distribution of unobserved state for next time period
            pi[1] <- Zpi[1]*S1[t]*p[t+1]            + Zpi[2]*S2[t]*p[t+1]
            pi[2] <- Zpi[1]*S1[t]*(1-p[t+1])        + Zpi[2]*S2[t]*(1-p[t+1])
            pi[3] <- Zpi[1]*(1-S1[t])*alpha1[t]     + Zpi[2]*(1-S2[t])*alpha2[t]
            pi[4] <- Zpi[1]*(1-S1[t])*(1-alpha1[t]) + Zpi[2]*(1-S2[t])*(1-alpha2[t])+Zpi[3]+Zpi[4]
            pi <- pi / sumZpi             ## normalize
        }
    }
    returnType(double())
    if(log) return(logL) else return(exp(logL))    ## return log-likelihood
  }
)
```

Description of the `dtroutDHMM` distribution's arguments and intermediate variables with all vectors and vector arguments pertaining to a given individual:

| | |
|---|---|
| **x** | Vector argument containing the observed capture history data |
| **length** | The length of the capture history |
| **mu.mH1** | Baseline harvest mortality hazard rate on the log scale |
| **mu.mO1** | Baseline above-dam background mortality hazard rate on the log scale |
| **mu.mO2** | Baseline below-dam background mortality hazard rateon the log scale |
| **mu.p** | Baseline ladder usage probability on the logit scale |
| **beta1.mO1** | Log-linear discharge effect on above-dam background mortality hazard rate |
| **beta1.mO2** | Log-linear discharge effect on below-dam background mortality hazard rate |
| **beta2.mO1** | Log-linear size effect on above-dam background mortality hazard rate |
| **betaS.mO** | Stocking effect on log background mortality hazard rate |
| **beta2.mO2** | Log-linear size effect on below-dam, background mortality hazard rate |
| **beta2.mH1** | Log-linear size effect on harvest mortality hazard rate |
| **beta4.mH1** | Log-quadratic size effect on harvest mortality hazard rate |
| **betaS.mH** | Stocking effect on log harvest mortality hazard rate |
| **beta1.p** | Logit-linear discharge effect on ladder usage probability |
| **beta2.p** | Logit-linear size effect on ladder usage probability |
| **beta3.p** | Logit-interactive discharge and size effect on ladder usage probability |
| **beta4.p** | Logit-quadratic size effect on ladder usage probability |
| **betaS.p** | Stocking effect on logit ladder usage probability |
| **size** | Vector argument containing standardized body sizes |
| **epsilon.mH** | Vector argument containing the random effects on harvest mortality hazard rates (log scale) |
| **epsilon.mO** | Vector argument containing the random effects on background mortality hazard rates (log scale) |
| **epsilon.p** | Vector argument containing the random effects on ladder usage probabilities (logit scale) |
| **discF** | Vector argument containing standardized fall discharge values |
| **discS** | Vector argument containing standardized summer discharge values |
| **r** | Vector argument containing reporting rates |
| **log** | Logical argument specifying whether the log of the likelihood should be returned |
| **mH1** | Vector containing harvest mortality hazard ratesl |
| **mO1** | Vector containing above-dam background mortality hazard rates |
| **mO2** | Vector containing below-dam background mortality hazard rates |
| **p** | Vector containing ladder usage probabilities |
| **S1** | Vector containing above-dam survival probabilities |
| **alpha1** | Vector containing above-dam ratio of harvest- to total mortality hazard rate |
| **S2** | Vector containing below-dam survival probabilities |
| **alpha2** | Vector containing below-dam ratio of harvest- to total mortality hazard rate |
| **logL** | Variable to store the log-likelihood of the observed capture history |
| **pi** | Probabilities of each latent state, conditioned on preceding observations |
| **Zpi** | Probability of current observed capture, conditioned on each possible latent state |
| **sumZpi** | Marginal probability of current observed capture, or the likelihood of one observation |

##### S2.3 Model Specification with Custom `dtroutDHMM` Likelihood Distribution

The complete NIMBLE code giving the hierarchical model specification is provided in the supplementary file `nimbleDHMM.R`. Below, we provide an itemized description of additional parameters and data objects appearing in the model specification within the complete code. Names flagged with *even* or *odd* in the code indicate parameters and data objects belonging to the analysis of even- and odd-year spawners respectively and are not described separately.

| | |
|---|---|
| **Mu.mH1** | Baseline harvest mortality hazard rate |
| **Mu.mO1** | Baseline above-dam background mortality hazard rate |
| **Mu.mO2** | Baseline below-dam background mortality hazard rate |
| **Mu.p** | Baseline ladder usage probability |
| **Mu.r1** | Reporting rate over the first time period |
| **sigma.mH** | Standard deviation of random time effects on harvest mortality hazard rates (log scale) |
| **sigma.mO** | Standard deviation of random time effects on background mortality hazard rates (log scale) |
| **sigma.p** | Standard deviation of random time effects on ladder usage probability (logit scale) |
| **sigma.r** | Standard deviation of random time effects on reporting rate (logit scale) |

| | |
|---|---|
| **y** | Matrix of capture histories |
| **period** | Vector of time periods (1-5) for all time intervals |
| **nind** | Total number of individuals |
| **n.occasions** | Total number of capture occasions (= time steps) |
| **first** | Vector of first capture occasions (marking events) of all individuals |
| **last** | Vector of last analysis occasion of all individuals |

### S3 Model Identifiability

In some models it is not possible to uniquely estimate every parameter regardless of the amount and quality of data. This issue is due to intrinsic non-identifiability of the parameters. In some models it may be obvious that parameters are non-identifiable. For example, in an occupancy model with a single visit per season parameters representing detection and occupancy only ever appear in the model as a product (MacKenzie et al. 2005). A unique parameter estimate exists only for the product of the two parameters, the individual parameters have multiple parameter estimates. However in many models, such as the multi-state capture-recapture models in this paper, it is not obvious whether or not the parameters are identifiable. In Section S3.1 we discuss the methods used to investigate non-identifiability. Results of identifiability tests are presented in Section S3.2.

#### S3.1 Methods for Investigating Intrinsic Non-Identifiability

We can investigate intrinsic identifiability using symbolic algebra (Cole et al. 2010). This involves using a symbolic algebra package (Cacthpole et al. 2002), and here we used Maple. First we consider an exhaustive summary of a model, which is a vector of parameter combinations that provide a unique representation of that model. For the here presented mark-recapture-recovery model for trout, the probabilities of each possible capture history for any individual fish form an exhaustive summary. There are three possible observations at each time point (where a time point here is every two years): the fish is caught alive in the fish ladder, represented by 1, the fish is caught and reported dead by a fisher, represented by 2 and the fish is not seen, represented by 3. A possible history for 5 years of study is $h = 13112$, where the fish is first caught in the dam at time 1, is not caught at time 2, and then is caught in the dam at times 3 and 4 and finally caught dead by a fisher at time 5. For constant parameters this history has probability

$$\Pr(h = 13112) = S_1^2(1 - p_1)S_2p_2p_1(1 - S_1)\alpha_1 r.$$

where $S_n$ ($e^{(-(m^H+m_n^O))}$) is the survival probability of state $n$ (1 = above-dam spawners, 2 = below-dam spawners), $p_n$ the ladder usage probability of state $n$, and $\alpha_n$ the ratio of harvest- to total mortality ($\frac{m^H}{m^H+m_n^O}$) of state $n$. The exhaustive summary consisting of all possible histories is then

$$\kappa = \begin{bmatrix} \Pr(h = 11111) \\ \Pr(h = 31111) \\ \Pr(h = 13111) \\ \vdots \\ \Pr(h = 13112) \\ \vdots \end{bmatrix} = \begin{bmatrix} S_1^4p_1^4 \\ S_1^3p_1^3 \\ S_1^3S_2p_1^2(1 - p_1)p_2 \\ \vdots \\ s_1^2(1 - p_1)S_2p_2p_1(1 - S_1)\alpha_1 r \\ \vdots \end{bmatrix}.$$

The full exhaustive summary can be found in the Maple file `constant.mw`.

We can test whether a model is non-identifiable using the symbolic method, which involves forming a derivative matrix by differentiating each term in the exhaustive summary with respect to each parameter. If the rank of the derivative matrix is less than the number of parameters, the model is non-identifiable. If the rank of the derivative matrix is equal to the number of parameters then the model is identifiable. The deficiency of a model is the difference between the number of parameters and the rank. An identifiable model has deficiency 0; a non-identifiable model has deficiency greater than 0 (Cole et al. 2010).

This symbolic method is demonstrated for the exhaustive summary above in the Maple file `constant.mw`. There are 6 top-level parameters: $m^H, m_1^O, m_2^O, p_1, p_2, r$. However the rank of the derivative matrix is 5, so this model is non-identifiable. The deficiency of this model is $6 - 5 = 1$. The Maple code also provides information on which parameters are confounded. Specifically, we obtain a reparameterization of the model in terms of parameter combinations which result in an identifiable model. In this simple case, the estimable parameter combinations are: $rm^H$, $m^H + m_1^O$, $m^H + m_2^O$, $p_1$, $p_2$.

For more complex models Maple runs out of memory calculating the rank of the derivative matrix (Jiang et al. 2007, Hunter and Caswell 2009, Cole and Morgan 2010a, Cole 2012). It is possible to use a numerical method instead (see Jiang et al. 2007, for example), however this can give the wrong result about identifiability (Cole and Morgan 2010a). Alternatively a hybrid symbolic-numerical method can be used (Choquet and Cole 2012) or the symbolic method can be extended by finding simpler exhaustive summaries (Cole et al. 2010). Here we use both of these approaches, which are described below.

The hybrid symbolic-numerical method involves finding the exact derivative matrix, but then evaluating the rank of the derivative matrix at random points in the parameter space; Choquet and Cole (2012) recommend 5 sets of random points. The maximum of the 5 ranks is the rank for the model. For the constant model above the Maple code `constant.mw` also demonstrates how the hybrid symbolic-numerical method is applied. In this case all 5 random points in the parameter space have a rank of 5. This again leads to a deficiency of 1 and shows that the model is non-identifiable.

Alternatively, a simpler exhaustive summary can be found by reparameterising the original exhaustive summary. The method is explained in Cole et al. (2010) and Cole (2012). Suppose that for this model the transition matrix has entries $\psi_{j,k,t}$ at time points $t = 1, ..., T$ for $j = 1, 2$ and $k = 1, 2, 3, 4$ (where $j$ = state[t] and $k$ = state[t+1]). Note that $\psi_{j,4,t} = 1 - \psi_{j,1,t} - \psi_{j,2,t} - \psi_{j,3,t}$. In the Maple code `simplerexproof.mw` we show that a simpler exhaustive summary consists of the terms:

$$\begin{array}{ll} \psi_{1,1,t} & \text{for } t = 1,\ldots,T-1 \\ \psi_{1,2,t-1}\psi_{2,1,t} & \text{for } t = 2,\ldots,T-1 \\ \psi_{1,3,t}r_{t+1} & \text{for } t = 1,\ldots,T-1 \\ \frac{\psi_{1,2,t-1}\psi_{2,3,t}}{\psi_{1,3,t}} & \text{for } t = 2,\ldots,T-1 \\ \frac{\psi_{1,2,t-1}\psi_{2,2,t}}{\psi_{1,2,t}} & \text{for } t = 2,\ldots,T-1 \end{array}$$

The models we consider include covariates and random effects. In some cases identifiability results for models with covariates can be deduced from equivalents models without covariates (Cole and Morgan 2010b), however in this case it is simpler to use the simpler exhaustive summary directly. For time varying covariates, such as the average discharge during the fall when trout are expected to migrate downriver, this just involves changing the exhaustive summary terms to the form involving covariates. For individual covariates, such as the individual body size at spawning, each individual adds a new exhaustive summary term to the original exhaustive summary involving probabilities of each capture history. However to obtain general results it is sufficient to use the simpler exhaustive summary repeated 2 times, with different covariate values on each repeat.

For example, suppose harvest mortality ($m^H$) depends on individual covariate $x_{i,t}$, background mortality ($m_n^O$) depends on on individual covariate $x_{i,t}$ and time dependent covariate $y_t$, the probability of using a fish ladder ($p$) is dependent on individual covariate $x_{i,t}$ and time dependent covariate $z_t$ and the reporting probability ($r$) is constant. When $T = 5$ we show in Maple code `covariates.mw` that the model has rank 11, but there are 12 parameters, therefore the model has deficiency 1 and is not identifiable.

Similarly methods can be extended for models with random effects (Janzen et al. 2016; 2017). An exhaustive summary for models with random effects consists of the probabilities of each history conditional on the random effects, $\Pr(h = |\mathbf{b})$, and the probability density function of the random effects $f(\mathbf{b})$, where $\mathbf{b}$ represents the random effects. This can be simplified to the simpler exhaustive summary described above conditional on the random effects with the addition of $f(\mathbf{b})$. Using the extension Theorem from Cole et al. (2010) we can consider this in two parts. First we consider the conditional simpler exhaustive summary. In the Maple code `randomeffects.mw` we consider the same example as above but with time varying random effects on ($m^H$). The conditional simpler exhaustive summary results in a derivative matrix with rank 12. Then we consider $f(\mathbf{b})$, but because there is only one parameter. When only one parameter is added Cole et al. (2010) show that the extension Theorem application is trivial, as the rank of the required derivative matrix would always be 1. This is the case for the rank of the derivative matrix for $f(\mathbf{b})$. The rank for the whole model would then be $12 + 1 = 13$, and as there are 13 parameters the deficiency is 0 and the model thus identifiable.

#### S3.2 Intrinsic Identifiability Results

Table S3.1 gives the identifiability results for a selection of relevant models. We consider models where the harvest mortality rate ($m^H$) and the hazard mortality rate ($m^O$) are constant, dependent on appropriate covariates and dependent on random year effects. The reporting rate ($r$) is allowed to be constant or time dependent. In Table S3.1 the probability of using the fish ladder ($p$) is always assumed to be dependent on covariates, as there is no change in deficiency when the parameter is constant. We also consider including random year effects on ($p$). There are three covariates used, one is an individual, time-varying covariate ($x_{i,t}$, representing body size), and two are time dependent covariates ($y_t$ and $z_t$, representing river discharge in the fall and summer respectively).

The results show that to ensure identifiability, either $m^H$ and $m_n^O$ need to depend on different covariates or random effects have to be included on at least $m^H$ or $r$. This makes sense when considering the combinations of parameters that were not separately estimable in the constant model: $rm^H$, $m^H + m_1^O$, $m^H + m_2^O$. The parameter $m^H$ is contained in all three expressions; therefore, model identifiability is conditional on separating out $m^H$. Previous research has shown that in some cases, adding information through covariates can separate out non-identifiable parameters (see, for example, Cole and Morgan 2010b, Royle et al. 2012). Here, adding a covariate to $m^H$ resulted in an identifiable model as long as this covariate was not shared with the confounded $m_n^O$ parameters (column 5 in Table S3.1). Just like covariates, random effects can also aid in achieving identifiability by adding more information on parameters. For that reason, including random effects on $m^H$ results in identifiable models. Adding random effects on $r$ also allows separating $m^H$, thus making models identifiable. Adding the same random effect on $m_1^O$ and $m_2^O$, on the other hand, does not provide independent information on $m^H$ since $m^H$ is confounded with both $m_1^O$ and $m_2^O$ in the same way.

Table S3.1: Results of identifiability tests of models with different covariate and random effect structures. The first four columns specify the fixed effect model used for reporting rate $r$, harvest mortality $m^H$, background mortality $m_n^O$, and ladder usage probability $p$. $x_{i,t}$ represents an individual, time-varying covariate (*e.g.* body size) and $y_t$ and $z_t$ represent two different time-varying environmental covariates (*e.g.* discharge during two seasons). The remaining columns show the deficiency of the model under different random effect structures. Deficiency is calculated as the number of parameters minus the rank of the derivative matrix (found using the hybrid symbolic numerical method), and models where deficiency $> 0$ are not identifiable.

| | | | | **Deficiency** | | | | | |
|---|---|---|---|---|---|---|---|---|---|
| $\boldsymbol{r}$ | $\boldsymbol{m^H}$ | $\boldsymbol{m_n^O}$ | $\boldsymbol{p}$ | **No RE** | **RE on $\boldsymbol{m^H}$** | **RE on $\boldsymbol{m_n^O}$** | **RE on $\boldsymbol{m^H}$ & $\boldsymbol{m_n^O}$** | **RE on $\boldsymbol{m^H}$ & $\boldsymbol{m_n^O}$ & $\boldsymbol{p}$** | **RE on $\boldsymbol{r}$** |
| $t$ | $x_{i,t}$ | $x_{i,t}$, $y_t$ | $x_{i,t+1}$, $z_{t+1}$ | 1 | **0** | 1 | **0** | **0** | - |
| . | $x_{i,t}$ | $x_{i,t}$, $y_t$ | $x_{i,t+1}$, $z_{t+1}$ | 1 | **0** | 1 | **0** | **0** | **0** |
| $t$ | $x_{i,t}$ | $y_t$ | $x_{i,t+1}$, $z_{t+1}$ | **0** | **0** | **0** | **0** | **0** | - |
| . | $x_{i,t}$ | $y_t$ | $x_{i,t+1}$, $z_{t+1}$ | **0** | **0** | **0** | **0** | **0** | **0** |
| $t$ | $x_{i,t}$ | $x_{i,t}$ | $x_{i,t+1}$, $z_{t+1}$ | 1 | **0** | 1 | **0** | **0** | - |
| . | $x_{i,t}$ | $x_{i,t}$ | $x_{i,t+1}$, $z_{t+1}$ | 1 | **0** | 1 | **0** | **0** | **0** |
| $t$ | $x_{i,t}$ | . | $x_{i,t+1}$, $z_{t+1}$ | **0** | **0** | **0** | **0** | **0** | - |
| . | $x_{i,t}$ | . | $x_{i,t+1}$, $z_{t+1}$ | **0** | **0** | **0** | **0** | **0** | **0** |
| $t$ | . | $x_{i,t}$, $y_t$ | $x_{i,t+1}$, $z_{t+1}$ | 1 | **0** | 1 | **0** | **0** | - |
| . | . | $x_{i,t}$, $y_t$ | $x_{i,t+1}$, $z_{t+1}$ | 1 | **0** | 1 | **0** | **0** | **0** |
| $t$ | . | $y_t$ | $x_{i,t+1}$, $z_{t+1}$ | 1 | **0** | 1 | **0** | **0** | - |
| . | . | $y_t$ | $x_{i,t+1}$, $z_{t+1}$ | 1 | **0** | 1 | **0** | **0** | **0** |
| $t$ | . | $x_{i,t}$ | $x_{i,t+1}$, $z_{t+1}$ | 1 | **0** | 1 | **0** | **0** | - |
| . | . | $x_{i,t}$ | $x_{i,t+1}$, $z_{t+1}$ | 1 | **0** | 1 | **0** | **0** | **0** |
| $t$ | . | . | $x_{i,t+1}$, $z_{t+1}$ | 1 | **0** | 1 | **0** | **0** | - |
| . | . | . | $x_{i,t+1}$, $z_{t+1}$ | 1 | **0** | 1 | **0** | **0** | **0** |

##### S3.3 Testing for Weak Identifiability using Prior-Posterior Overlaps

In some cases, intrinsically identifiable models may be near-redundant, resulting in in parameter estimates with low precision and large influence of prior selection in Bayesian implementations. Garrett and Zeger (2000) investigate near-redundancy by examining the prior-posterior overlap. For latent class models (Garrett and Zeger 2000) and capture-recapture models (Gimenez et al. 2009) an overlap of 35% is commonly used to classify a model as weakly identifiable. In Figure S3.1 we display the prior-posterior overlap of all main parameters in our model. The only parameter with an overlap clearly exceeding 35% is $r_1$, the reporting rate over the very first 2-year interval. Since we know that the parameter is intrinsically identifiable (section S3.2) this indicates that the amount of data may be insufficient to precisely estimate this parameter. We assessed whether this potential weak identifiability in the first reporting rate could affect the estimation of the biological parameters in the model by comparing posterior distributions of the model with autocorrelated $r_t$ to those obtained under the assumption of an alternative reporting rate model. This alternative model, which assumed five period-specific mean reporting rates ($\mu_1^r$, $\mu_2^r$, $\mu_3^r$, $\mu_4^r$, $\mu_5^r$) and random variation around those, did not show any sign of non-identifiability (prior-posterior overlaps of all parameters ¡ 35%, Figure S3.2). Posterior distributions for biological parameters obtained from both models were basically identical (Figure S3.3), illustrating that the potential weak identifiability of the initial reporting rate $r_1$ did not affect biological predictions here and was thus unproblematic.

We note that the reason why we find a weakly identifiable parameter in the model with autocorrelated $r_t$ but not in the model with period-specific $r_t$ may well be related, again, to random effect. Intrinsic identifiability analysis (above) showed that random effects on at least $m^H$ or $r$ were prerequisite to obtaining an identifiable model. Autocorrelative models are very flexible, and here this resulted in lower residual variance ($\sigma^r$, Figure S3.3); as residual variance decreases, the random effects necessary for identifiability will approach 0, which may then lead to the model becoming less identifiable.

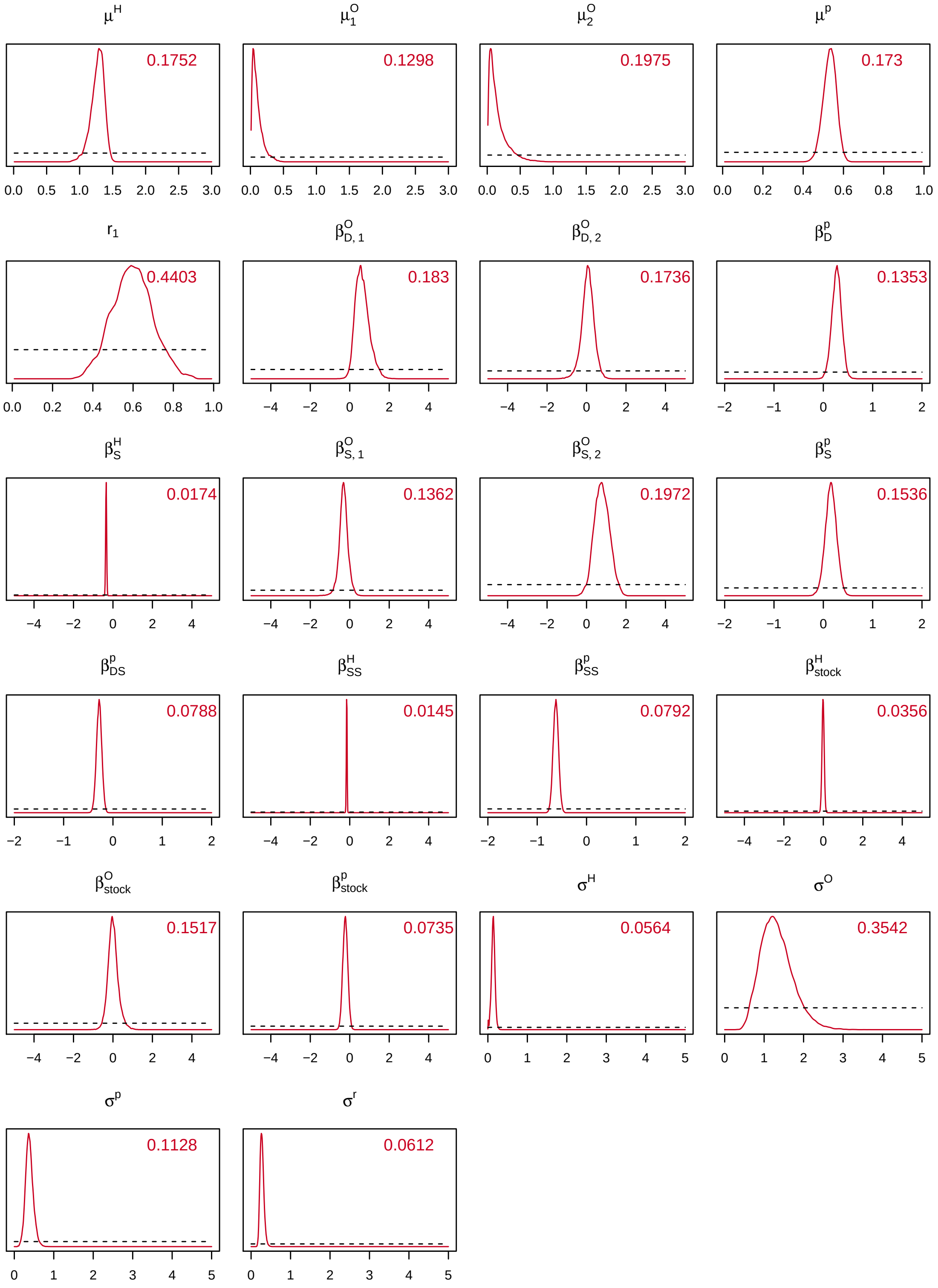

Figure S3.1: Densities of prior (dashed black line) and posterior (solid red line) distributions for all main parameters in the final model with autocorrelated reporting rates. The red numbers indicate the proportion overlap between prior and posterior.

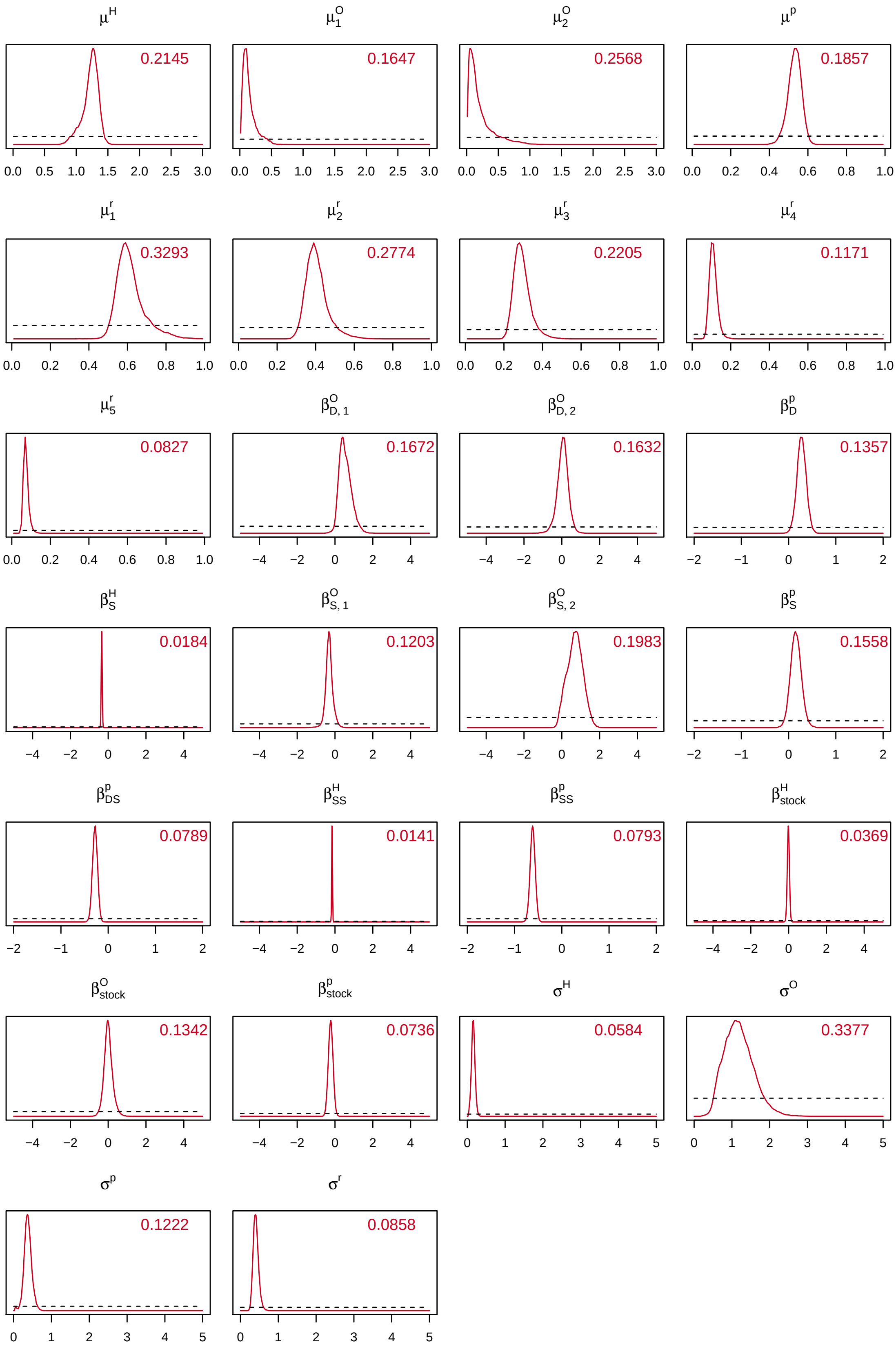

Figure S3.2: Densities of prior (dashed black line) and posterior (solid red line) distributions for all main parameters in the alternative model with period-specific mean reporting rates. The red numbers indicate the proportion overlap between prior and posterior.

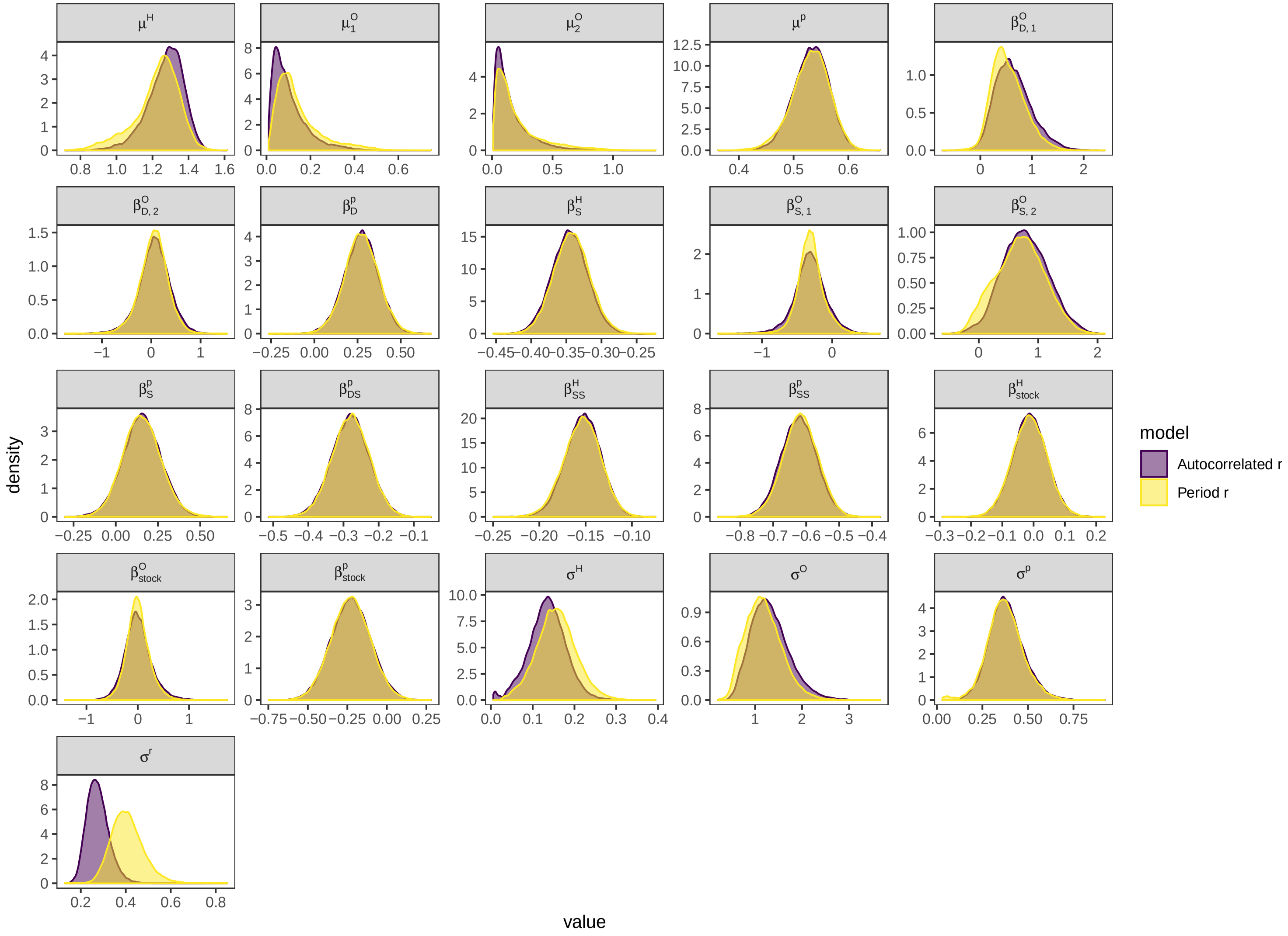

Figure S3.3: Posterior distributions for all main parameters estimated from the final model with autocorrelated reporting rates (purple) and the alternative model with period-specific mean reporting rates (yellow).

### S4 Posterior Predictive Checks

#### S4.1 Simulation of new data sets and extraction of test statistics

Assessing model fit using posterior predictive checks (PPCs) relies on the comparison of the real dataset to replicate datasets generated from the joint posterior of a Bayesian model (Conn et al. 2018). We selected 500 evenly spaced samples from our model's joint posterior distributions and simulated 10 replicate datasets for each of those 500 posterior samples (= 5000 simulated datasets). Specifically, we simulated capture histories for each individual, assuming initial capture in the same year and at the same body size as in the real data and calculating individual- and year-specific transition probabilities using the joint posterior sample of model parameters. Effects of river discharge and individual body size on transition probabilities were calculated using the same covariates employed during the model run. Random year effects were not re-sampled for data simulation and we used posterior samples of the estimated random effect levels to include among-year variation in predicted transition probabilities. Simulated datasets consequently contained the same number of individuals and years (divided into even- and odd-year spawners) as the real dataset.

We selected two general characteristics of our datasets for comparison: the number of individuals recaptured (both in the fish ladder and through harvest) and the size distribution of those individuals. When further describing size distribution through mean, median, and standard deviation, this resulted in the following eight general test statistics:

- Number of recaptures
- Mean size of recaptured individuals
- Median size of recaptured individuals
- Size standard deviation of recaptured individuals
- Number of harvests
- Mean size of harvested individuals
- Median size of harvested individuals
- Size standard deviation of harvested individuals

We then further specified these test statistics by separately considering distributions of marking size (MarkSize) and recapture size (RecapSize) of recaptured/harvested individuals[3], and evaluating all met-

[3]The rationale behind making this separation is that these two size metrics are influenced to a different degree by assumptions about growth. Contrasting model fit in the context of the two metrics may thus give some insights into the contribution of the growth model (Appendix S5) to overall model fit, or lack thereof. Note, however, that we evaluated recapture size metrics only for ladder recaptures. We did not do so for harvests, since harvest usually takes place before an individual has realized its full 2-year growth, and often within less than a year after spawning.

rics at time $t + 1$ (two years after marking) and at time $t + 2$ (four years after marking). An overview of the resulting 22 test statistics is provided in Table S4.1.

#### S4.2 Comparing real and simulated datasets

Following current recommendations (Conn et al. 2018), we used both visual tools and Bayesian p-values to assess the fit of our model. Specifically, we plotted representations of our test statistics derived from simulated datasets versus those derived from the real dataset, and calculated Bayesian p-values as the proportion of test statistics derived from simulated datasets that were higher than the test statistics derived from the real dataset. p-values surrounding 0.5 thus indicate good fit of the model to the data, while p-values at the extremes (0 or 1) provide evidence for lack of model fit or non-uniform p-value distribution (Besbeas and Morgan 2014). In the next section, we present the results of our model fit assessment for test statistics related to recapture numbers and recapture size distributions, at the level of the whole dataset and by marking cohort.

#### S4.3 Results: Model goodness-of-fit

PPCs indicated that overall and regarding the majority of evaluated test statistics, our model produced a decent fit to the data. Of the 22 Bayesian p-values we calculated for test statistics across the entire dataset, 16 (73%) fell between 0.1 and 0.9, and 13 (59%) fell between 0.2 and 0.8 (Table S4.1). p-Values varied substantially across different marking cohorts, but when averaged over all cohorts fell between 0.37 and 0.59 (Figure S4.1).

Our model performed well when predicting the numbers of individuals recaptured two and four years after marking (Figure S4.2a & b) and harvested within two years after marking (Figure S4.2c). For harvest between two and four years after marking, however, the model had a tendency to overestimate numbers (Figure S4.2d) when considering the whole dataset. The average of cohort-specific p-values, on the other hand, was 0.52 (Figure S4.1f). A closer look at cohort-specific model fit and p-values (Figure S4.3) revealed that overestimation of t+2 harvest numbers predominantly occurred in the first half of the time series and was most severe for the very first few marking cohorts. The model frequently also predicted lower than observed mean sizes (Table S4.1) for t+2 harvests, and investigating the entire size distributions revealed that this was due to a marked absence of relatively small individuals in t+2 harvests in the real data (as opposed to simulation, Figure S4.4).

The observed marking and recapture size distributions all fit well within the limits of what the model was able to predict (Figure S4.4), supporting an overall decent fit of the model also regarding size dynamics. Considering specific size metrics (and associated p-values) made some discrepancies between

Table S4.1: The 22 evaluated test statistics including Bayesian p-values calculated across entire datasets (all marking cohorts pooled).

| Test statistic | Capture type | Size measure | Time | p-value |
|---|---|---|---|---|
| recapNo | Recapture | MarkSize | t+1 | 0.2768 |
| sizeMean | Recapture | MarkSize | t+1 | 0.4818 |
| sizeMedian | Recapture | MarkSize | t+1 | 0.3846 |
| sizeSD | Recapture | MarkSize | t+1 | 0.0194 |
| recapNo | Recapture | MarkSize | t+2 | 0.4922 |
| sizeMean | Recapture | MarkSize | t+2 | 0.5182 |
| sizeMedian | Recapture | MarkSize | t+2 | 0.4760 |
| sizeSD | Recapture | RecapSize | t+2 | 0.5092 |
| sizeMean | Recapture | RecapSize | t+1 | 0.0002 |
| sizeMedian | Recapture | RecapSize | t+1 | 0.0000 |
| sizeSD | Recapture | RecapSize | t+1 | 0.6578 |
| sizeMean | Recapture | RecapSize | t+2 | 0.1866 |
| sizeMedian | Recapture | RecapSize | t+2 | 0.1658 |
| sizeSD | Recapture | RecapSize | t+2 | 0.9432 |
| recapNo | Harvest | MarkSize | t+1 | 0.2054 |
| sizeMean | Harvest | MarkSize | t+1 | 0.3236 |
| sizeMedian | Harvest | MarkSize | t+1 | 0.5564 |
| sizeSD | Harvest | MarkSize | t+1 | 0.8720 |
| recapNo | Harvest | MarkSize | t+2 | 0.9992 |
| sizeMean | Harvest | MarkSize | t+2 | 0.1418 |
| sizeMedian | Harvest | MarkSize | t+2 | 0.2040 |
| sizeSD | Harvest | MarkSize | t+2 | 0.9594 |

observed and predicted size distributions more obvious. Overall, the model performed well at predicting mean and median sizes of recaptures and harvests (Figures S4.5 & S4.6, Table S4.1). However, it consistently underestimated mean/median recapture size for recaptures four years after marking (t+2). Investigating the corresponding predicted and observed size distribution (Figure S4.4d) revealed that the main reason for the underestimation of mean/median size by the model lay in its failure to reproduce the non-normality in the observed size distribution, which was characterised by a surprisingly low number of relatively small individuals. Underestimation of mean/median size was also visible when considering cohort-specific p-values (Figure S4.1c) and – similar to the mismatch of t+2 harvest numbers – appeared to be largely driven by earlier years (Figure S4.7). Notably, the most extreme p-values in the beginning of the time series came from years 1966, 1967, 1969, 1971, 1973, and 1975. With the exception of 1966, all of these pertain to odd-year spawner, indicating that specific birth and/or spawning cohorts may contribute disproportionately to lack of fit. Standard deviation of size distributions was clearly underestimated by the model on some occasions (t+1 recapture marking size) and overestimated on others (t+2 recapture recapture size, t+2 harvest marking sizes, Figure S4.8). However, considering the entire size distributions (Figure S4.4) illustrated that over-/underestimation of size standard deviation by no means indicated severe bias in overall predictions of size distributions in recaptures and harvests. Unlike above-mentioned biases, the model's ability to predict size standard deviation was not subject to any temporal patterns.

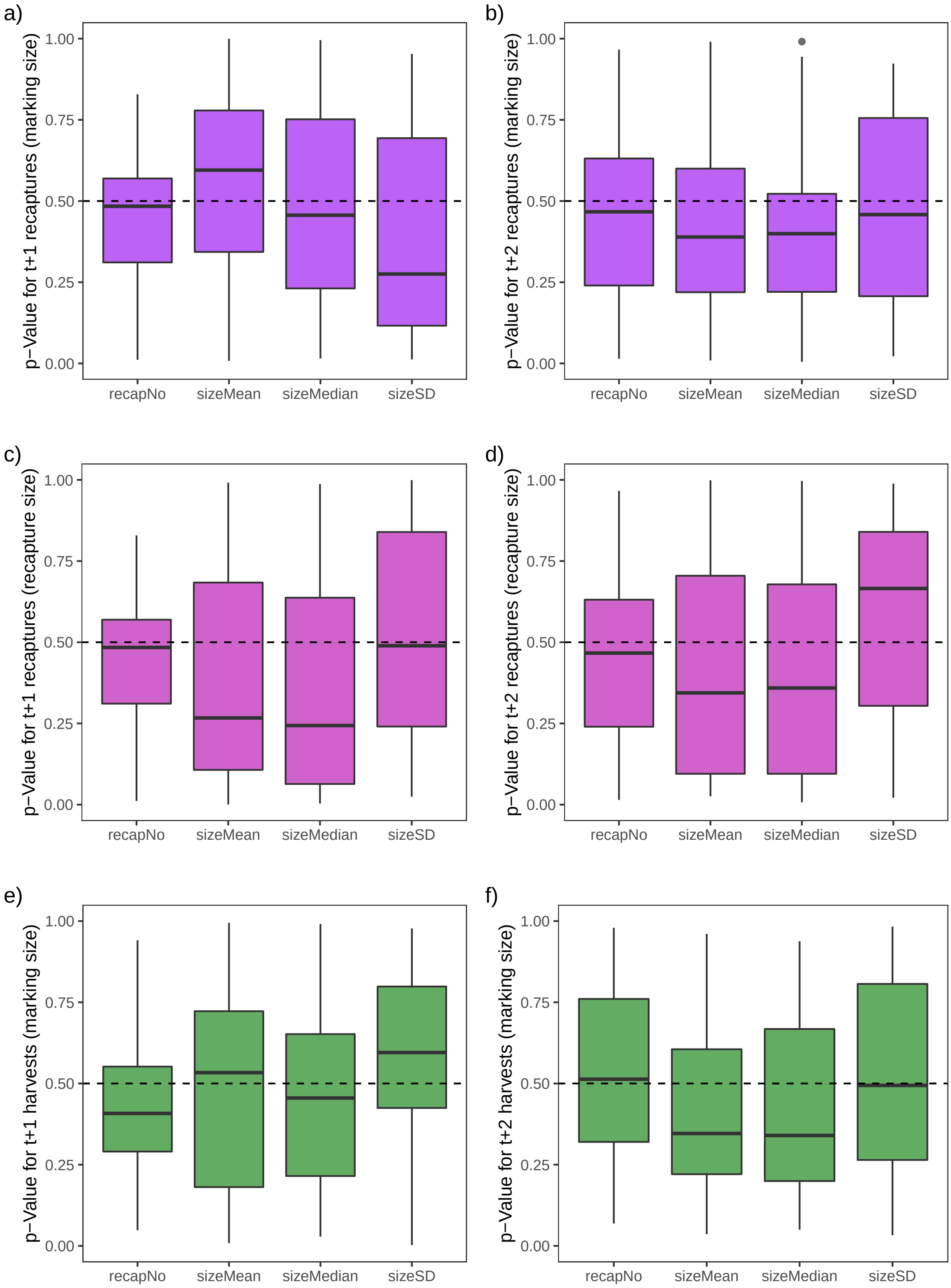

Figure S4.1: Distributions of marking cohort-specific p-Values for all evaluated test statistics two (t+1, left column) and four (t+2, right column) after marking.

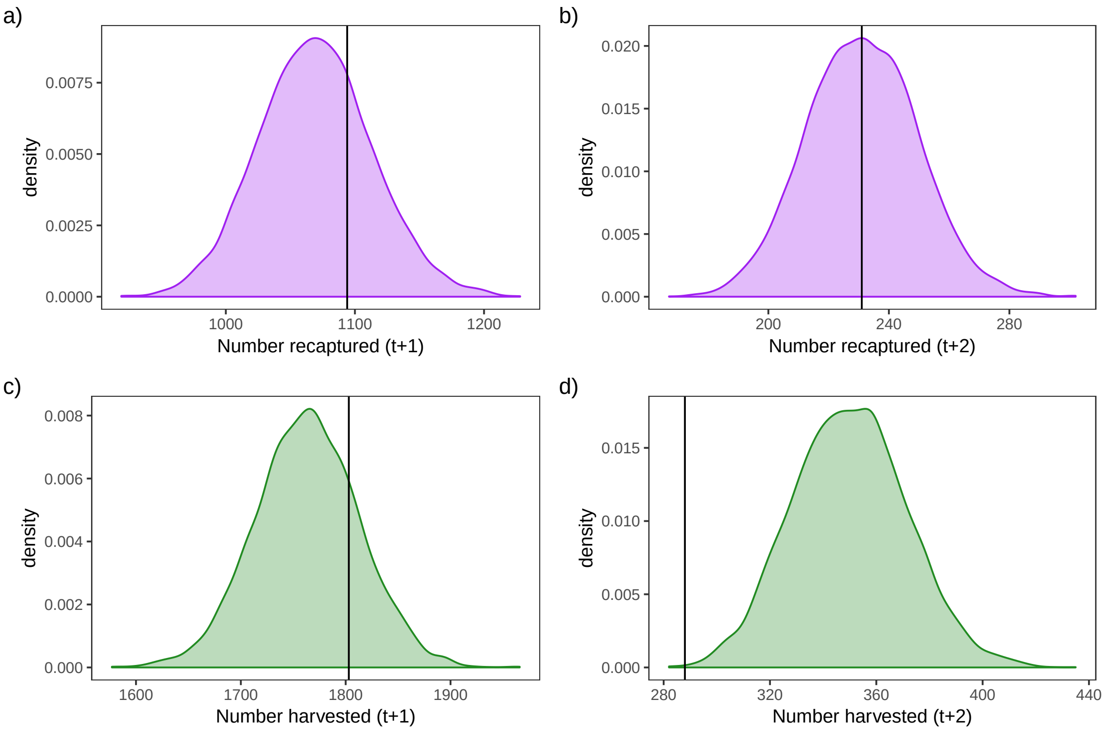

Figure S4.2: Distributions of the number of ladder recaptures and harvests two (t+1) and four years (t+2) after marking across the entire simulated datasets (all marking cohorts pooled). Black line marks the corresponding number recaptured/harvested in the real dataset.

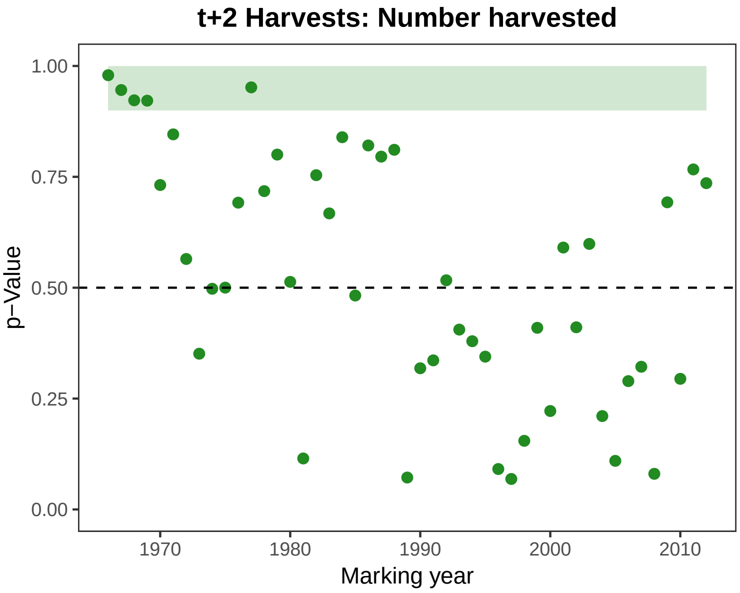

Figure S4.3: p-Values for the number of individuals harvested two to four years after marking across time (marking years). The shaded area highlights marking cohorts with potentially large influence on entire dataset p-value.

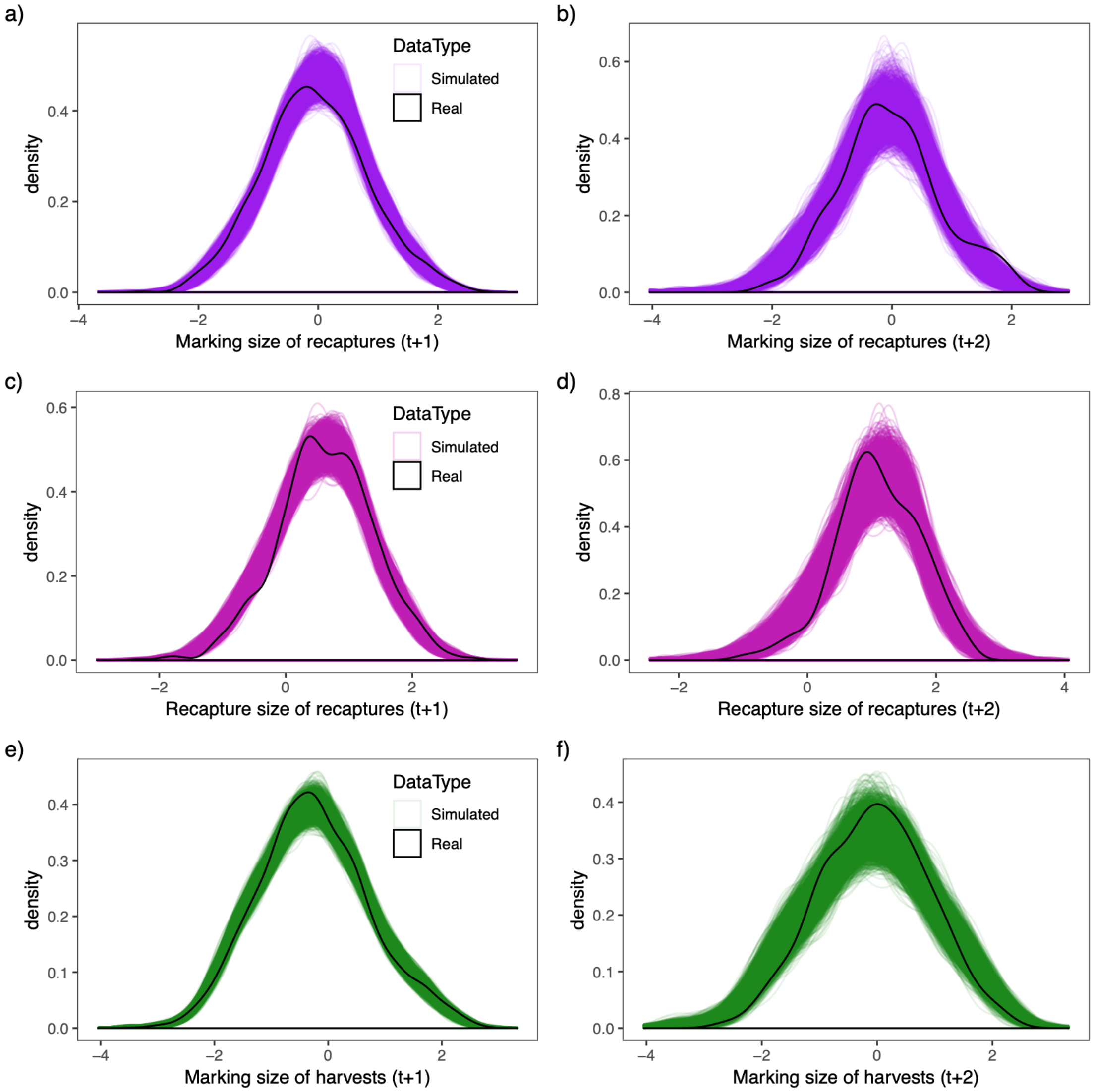

Figure S4.4: Distributions of marking size of ladder recaptures (top), recapture size of ladder recaptured (middle), and marking size of harvests (bottom) two (t+1) and four (t+2) years after marking. Each colored line represents the size distribution from one entire simulated dataset (N = 5000). The black line is the same distribution as found in the real dataset. All size measures are standardized.

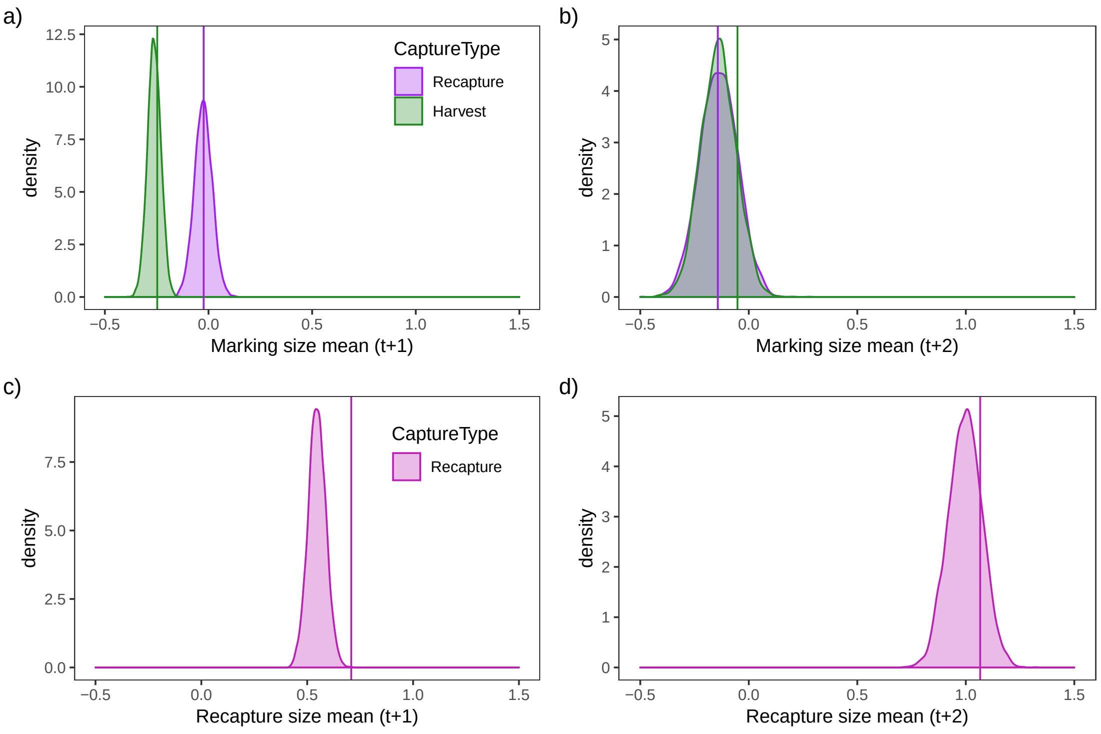

Figure S4.5: Distributions of the mean marking and recapture sizes during recaptures and harvests two (t+1) and four years (t+2) after marking, across the entire simulated datasets (all marking cohorts pooled). The solid line marks the corresponding mean size in the real dataset. Size here is standardized.

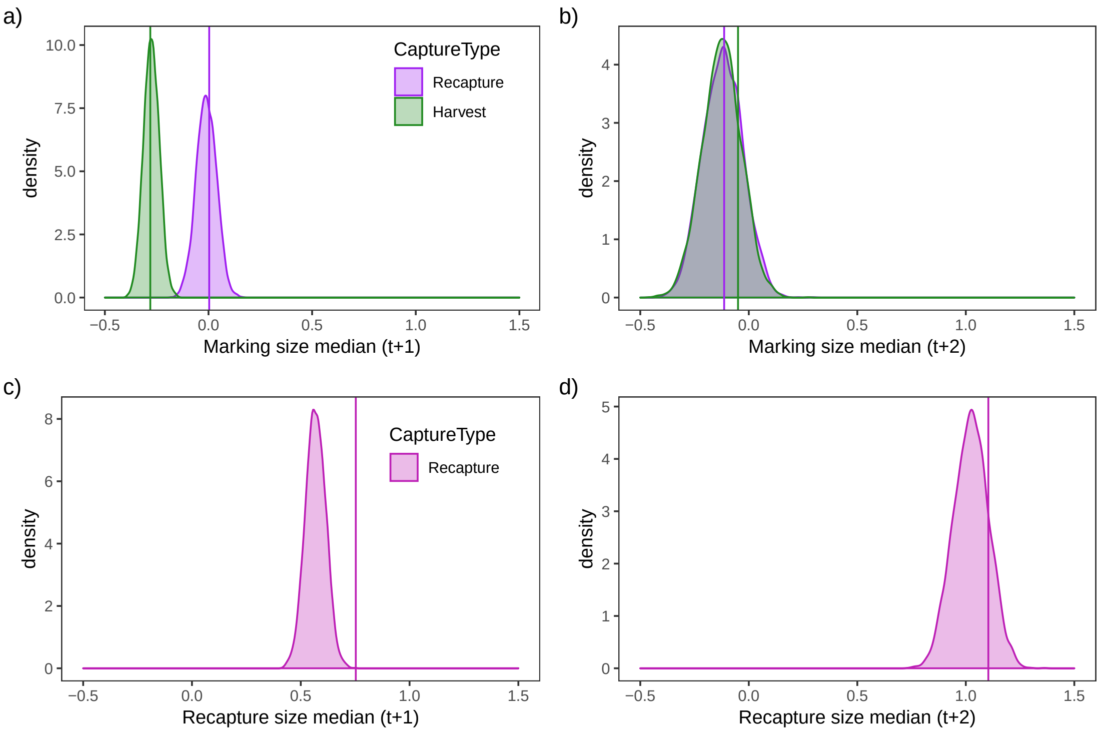

Figure S4.6: Distributions of the median marking and recapture sizes during recaptures and harvests two (t+1) and four years (t+2) after marking, across the entire simulated datasets (all marking cohorts pooled). The solid line marks the corresponding median size in the real dataset. Size here is standardized.

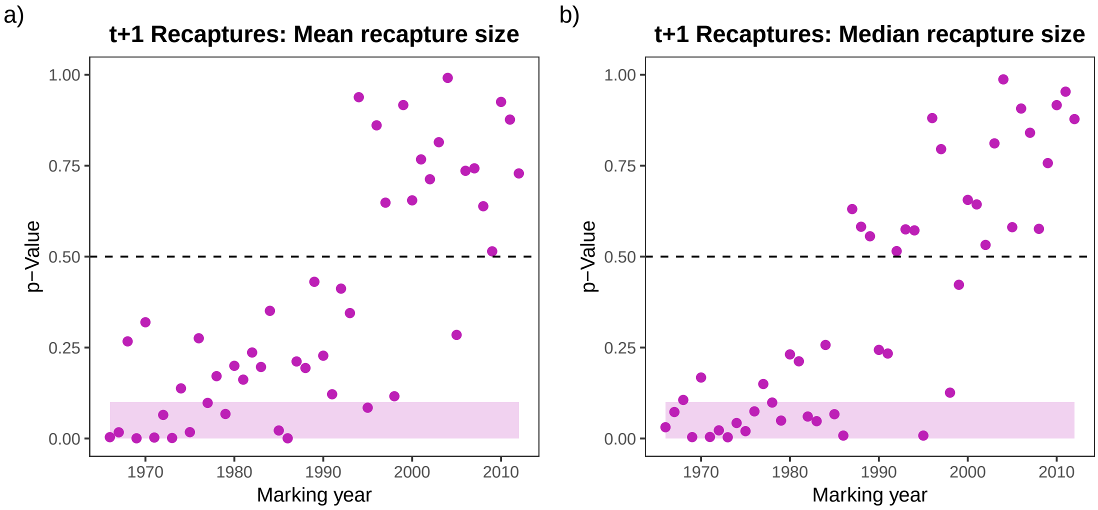

Figure S4.7: p-Values for the mean and median of standardized size of individuals recaptured two years after marking across time (marking years). The shaded areas highlight marking cohorts with potentially large influence on entire dataset p-value.

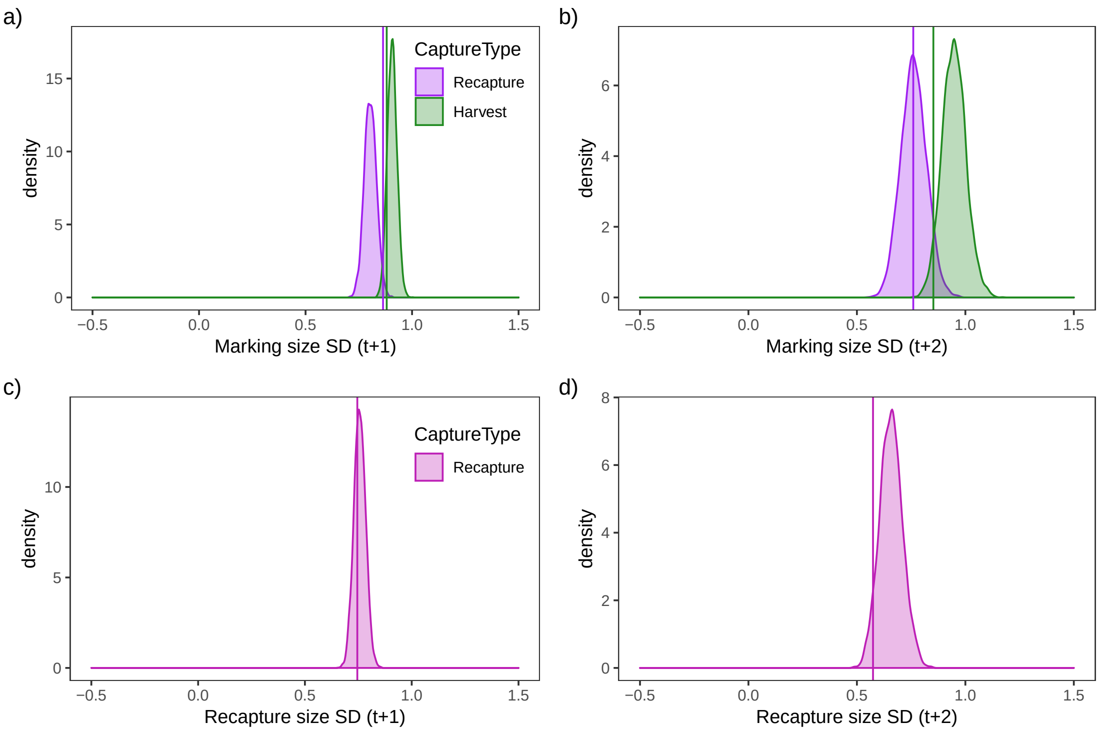

Figure S4.8: Distributions of the marking and recapture size standard deviation during recaptures and harvests two (t+1) and four years (t+2) after marking, across the entire simulated datasets (all marking cohorts pooled). The solid line marks the corresponding size standard deviation in the real dataset. Size here is standardized.

### S5 Growth Model and Size Extrapolation

#### S5.1 Re-fitting of the growth model

A subset of trout asceding the ladder had scale samples taken at capture, and the year rings on those scales were used to reconstruct individual growth histories (dataset described in Moe et al. 2019). Nater et al. (2018) analysed these data using a biphasic growth model which divided growth into a juvenile river period and an adult lake period while accounting for among-individual, among-year, and residual variation, as well as measurement error. Specifically, they modelled juvenile growth as a linear process and adult growth as following a von Bertalanffy curve such that $L_{i,t} = L_{i,t-1} + (L_{i,\infty} - L_{i,t-1})(1 - e^{-k_{i,t-1}})$, where $L_{i,t}$ is the length of individual $i$ in year $t$, $L_{i,\infty}$ is an individual's asymptotic length and $k_{i,t}$ its year-specific lake growth rate (Nater et al. 2018).

For the purpose of this study, we re-fit the model of Nater et al. (2018) (without environmental covariates) to an extended data set of 6,843 individual growth histories spanning the years 1952 to 2003. We further included potential differences between wild-hatched and hatchery-reared (= stocked) trout, and male and female trout, by including effects of origin and sex (if available) on asymptotic size and lake growth rate. The re-fitting of the growth model was done with JAGS 4.2.0 (Plummer 2003) via the `dclone` package (Sólymos 2010) in R 3.3.0 (R Core Team 2018), generating 3 chains of 50,000 iterations each (of which the first 20,000 were discarded as burn-in).

#### S5.2 Parameter estimates from extended growth model

The parameter estimates from the re-fitted growth analysis are reported in Table S5.1. With the extended dataset, we were unable to separately estimate individual variation in asymptotic size and therefore excluded this parameter from the analyses. New estimates of individual variation in lake growth rates were consequently higher, but otherwise posterior estimates closely resembled those reported previously (Nater et al. 2018). Additionally, we found that sex had larger effects on growth parameters than origin, and that male and/or stocked trout grew slower (lower lake growth rate) but towards a larger asymptotic size than female and/or wild-hatched trout.

Table S5.1: Posterior means, standard deviations (SD), 2.5% and 97.5% quantiles for parameters of the re-fitted growth model. RE is used as an abbreviation for random effect.

| **Parameter** | **Unit** | **Abbreviation** | **Mean** | **SD** | **2.5%** | **97.5%** |
|---|---|---|---|---|---|---|
| Baseline river growth rate | mm | h0 | 69.206 | 1.150 | 66.684 | 71.362 |
| Male effect on h | mm | betaM.h | 0.556 | 0.342 | -0.118 | 1.227 |
| Length at birth | mm | mu0 | 1.548 | 0.290 | 0.976 | 2.120 |
| Time trend on h | mm/year | betaYR | -0.224 | 0.038 | -0.297 | -0.147 |
| Average lake growth rate (wild) | - | k0[1] | 0.155 | 0.004 | 0.148 | 0.162 |
| Average lake growth rate (stocked) | - | k0[2] | 0.139 | 0.005 | 0.130 | 0.148 |
| Spawning effect on log(k) | - | betaS | -1.770 | 0.034 | -1.838 | -1.705 |
| Male effect on log(k) | - | betaM.k | -0.126 | 0.020 | -0.166 | -0.086 |
| Average asymptotic size (wild) | mm | mu_inf0[1] | 1295.570 | 11.525 | 1272.937 | 1318.251 |
| Average asymptotic size (stocked) | mm | mu_inf0[2] | 1435.445 | 25.177 | 1388.476 | 1487.319 |
| Male effect on mu_inf | - | betaM.mu | 157.966 | 18.880 | 121.537 | 194.764 |
| SD of individual REs on h | mm | sigmaR.i | 7.715 | 0.173 | 7.381 | 8.059 |
| SD of year REs on h | mm | sigmaR.t | 3.221 | 0.425 | 2.495 | 4.144 |
| SD of individual REs on log(k) | - | sigmaL.i.k | 0.154 | 0.002 | 0.094 | 0.159 |
| SD of year REs on log(k) | - | sigmaL.t.k | 0.117 | 0.013 | 0.094 | 0.146 |
| Residual process SD (river) | mm | sigmaR.R | 13.714 | 0.128 | 13.464 | 13.969 |
| Residual process SD (lake) | mm | sigmaL.R | 19.920 | 0.229 | 19.469 | 20.365 |
| Measurement error SD (river) | mm | sigmaR.M | 5.353 | 0.101 | 5.155 | 5.552 |
| Measurement error SD (lake) | mm | sigmaL.M | 15.914 | 0.183 | 15.554 | 16.273 |

##### S5.3 Extrapolating size for mark-recapture modelling

Whenever size covariate values were missing for a given individual in a given time-interval, we calculated them using the parameter estimates of the re-fitted biphasic growth model. All trout in the mark-recapture-recovery data are mature adults, so we used only the von Bertalanffy model (adult growth) and extrapolated length for two-year intervals (year $t-2$ to year $t$) as follows:

$$L_{i,t} = \underbrace{L_{i,t-2} + (L_{i,\infty} - L_{i,t-2})(1 - e^{-k_{N,i,t-2}})}_{L_{i,t-1}} + (L_{i,\infty} - L_{i,t-1})(1 - e^{-k_{S,i,t-1}})F$$

Here, $k_N$ and $k_S$ are used for lake growth rate in non-spawning and spawning years respectively. $L_{i,t-2}$ can be the measured length of a trout captured in the fish ladder, or an extrapolated value. $L_{i,t-1}$ is always an extrapolated value, as trout can be captured and measured in the ladder only every other year. For each individual, we used the origin- and sex-specific baseline $k$ and $L_\infty$. For the 343 individuals with unknown sex, we used parameter estimates from a growth model run without sex effects.

The re-fitted growth models also produced estimates of the specific growth random effect levels for the years 1966 to 2003 and for the 6,843 individuals contained in both the growth and the mark-recapture-recovery data. For these years and individuals, we also included the respective random effect levels for calculating missing values in the size covariates. For the remaining years and individuals, we set random effect levels to 0 instead.

$F$ in the above equation represents a correction factor. When comparing sizes predicted using the original growth model ($F = 1$) to sizes measured in the fish ladder at later recaptures, we found that the model consistently overestimated size (Figure S5.1, top-left). The degree of overestimation further scaled with growth increment, with more severe overestimation for the larger growth increments of smaller individuals (Figure S5.1, bottom-left). Plausible explanations for the model's overestimation of growth include violations of the assumption of no growth in fall/winter (measuring trout that have not completed their annual growth yet in the fish ladder) and bias in the data used for estimating lake growth parameters towards subadult trout whose growth patterns may differ from those of mature trout. Overestimating growth, and therefore the size covariate used in CMRR analyses, resulted in biased estimates of size-dependence in mortality and ladder usage and ultimately to poor fit of the CMRR model to the data (as indicated by posterior predictive checks, Appendix S4). We were able to greatly improve model fit by avoiding growth overestimation through the use of the correction factor $F$ while extrapolating size. To do so, we determined the optimal value for $F$ by minimizing average difference between observed and predicted sizes (Figure S5.2). We ultimately used $F = 0.63$, the optimized correction factor for extrapolation when both individual and year random effects are available (most of our extrapolated datapoints fell into this category). The resulting reduced growth model predicted sizes that fit well with observations (Figure S5.1, right panels).

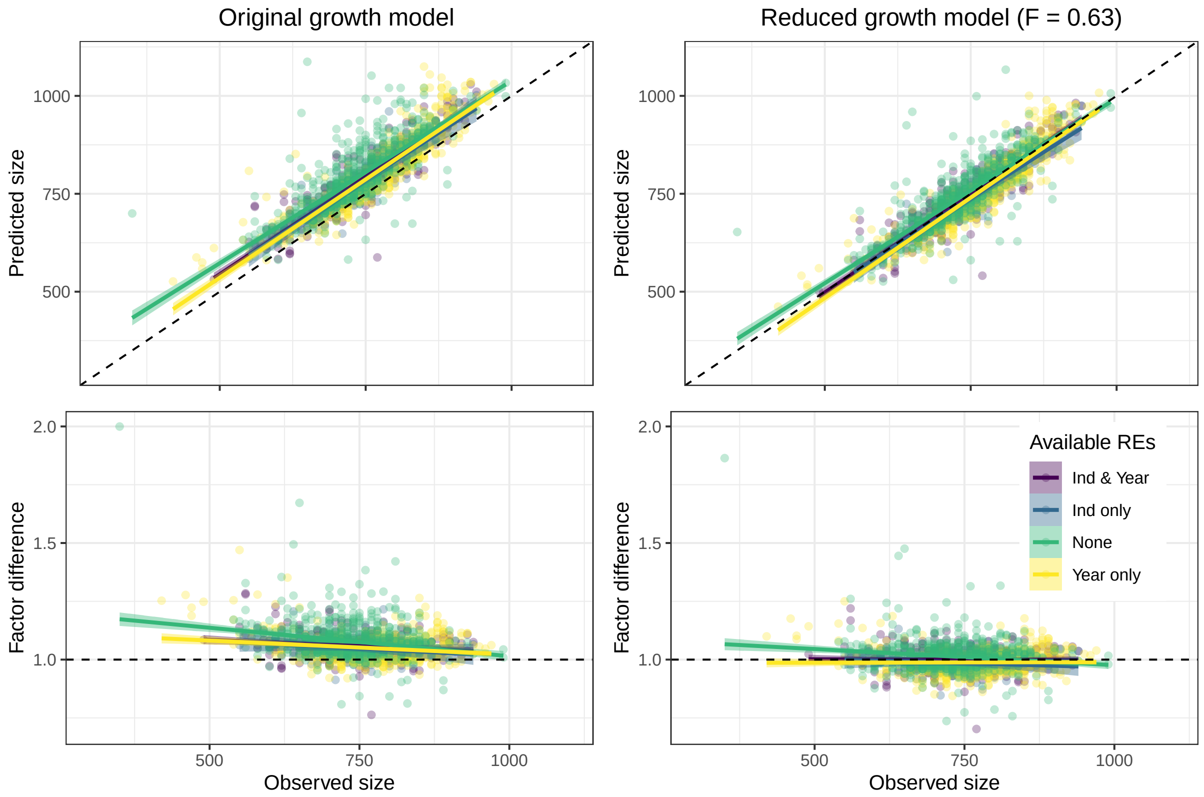

Figure S5.1: Top panels: comparison of observed sizes (measured in the fish ladder) and sizes predicted using the original growth model (left) and a reduced growth model with $F = 0.63$ (right). Bottom panels: relationship of observed sizes and factor difference of predicted sizes. Colors indicate which random effects were available for predicting sizes.

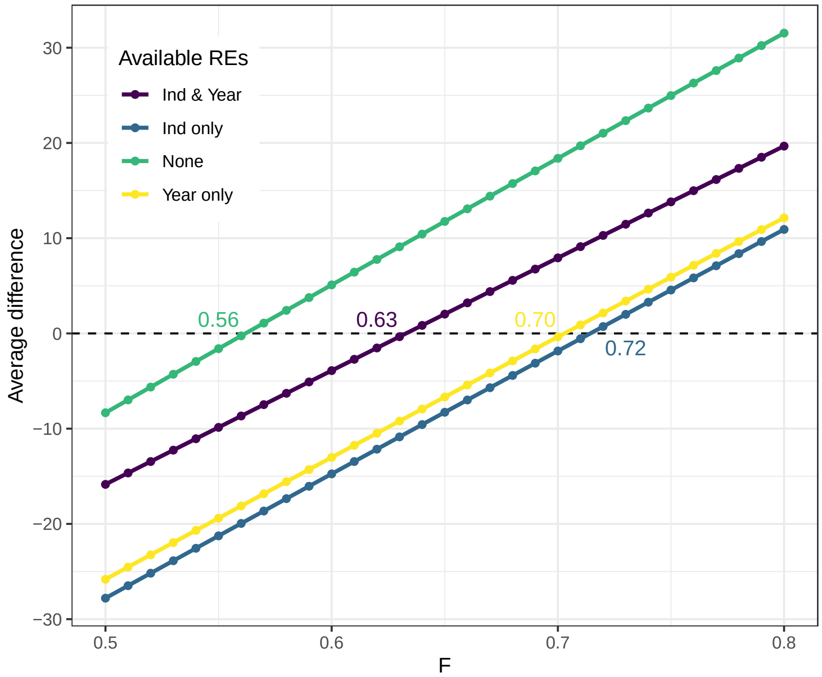

Figure S5.2: Average difference between observed sizes (measured in the fish ladder) and sizes predicted using the growth models with different values for the correction factor $F$. Numbers indicate the value of $F$ closest to 0 difference under different random effect availability (colors).

#### S6 Extended Model with Correlated Random Effects

We attempted to estimate the temporal correlation between harvest and background mortality hazard rates. To do so, we re-expressed the random effects on the hazard rates such that

$$\epsilon_t^H = \sigma_t^H * \xi_t^H, \qquad \xi_t^H \sim Normal(0,1)$$
$$\epsilon_t^O = \xi_t^O + \tau * \xi_t^H, \quad \xi_t^O \sim Normal(0,\sigma_t^O)$$

where $\sigma_t^H$ and $\sigma_t^O$ are the standard deviations for the random effects on harvest and background mortality hazard rates respectively. The scaling parameter $\tau$ can then be used to calculate the correlation between random effects as $C = \tau/\sqrt{(\sigma_t^O)^2 + \tau^2}$. We selected this parameterization to model the correlation between $\epsilon_t^H$ and $\epsilon_t^O$ since it induces the same marginal variances as the equivalent multivariate normal prior, but is computationally faster on account of forgoing the full multivariate normal distribution and associated matrix operations.

In the trout analysis, we found that the available data was insufficient to yield a precise estimate of the random effects correlation $C$. The resulting posterior distribution for $C$ (Figure S6.1) spanned the entire possible range from -1 to 1 and its median of 0.667 (95% CI [-0.622, 0.854] was indistinguishable from 0. Neither estimates of other parameters nor goodness-of-fit of the model were affected by the inclusion of random effect correlation.

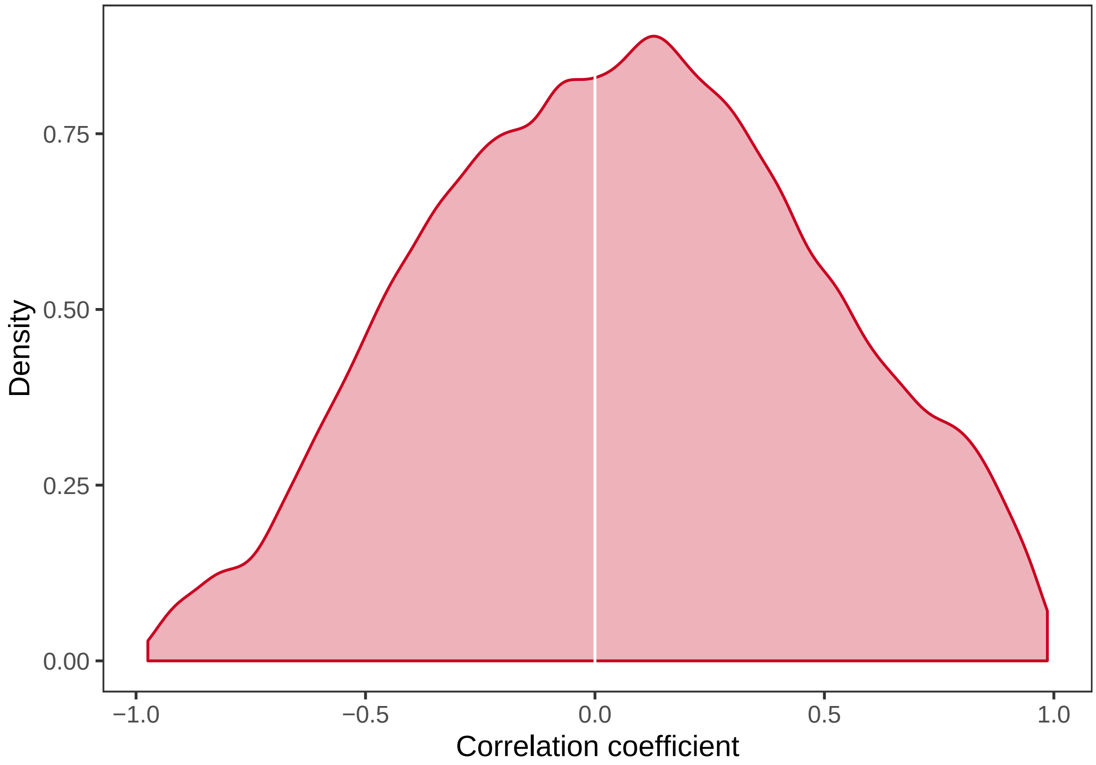

Figure S6.1: Posterior distributions of coefficient $C$ of the temporal correlation between harvest and background mortality hazard rates estimated by an extended model. White line marks $C = 0$.

#### References

Besbeas, P. and B. J. Morgan, 2014. Goodness-of-fit of integrated population models using calibrated simulation. *Methods in Ecology and Evolution* **5**:1373–1382.

Cacthpole, E. A., B. J. T. Morgan, and A. Viallefont, 2002. Solving problems in parameter redundancy using computer algebra. *Journal of Applied Statistic* **29**:625–636.

Choquet, R. and D. J. Cole, 2012. A hybrid symbolic-numerical method for determinig model structure. *Mathematical Biosciences* **236**:117–125.

Cole, D. J., 2012. Determining parameter redundancy of multi-state mark–recapture models for sea birds. *Journal of Ornithology* **152**:305–315.

Cole, D. J., B. J. Morgan, and D. Titterington, 2010. Determining the parametric structure of models. *Mathematical Biosciences* **228**:16–30.

Cole, D. J. and B. J. T. Morgan, 2010a. A note on determining parameter redundancy in age-dependent tag return models for estimating fishing mortality, natural mortality and selectivity. *Journal of Agricultural Biological and Environmental Statistics* **15**:431–434.

Cole, D. J. and B. J. T. Morgan, 2010b. Parameter redundancy with covariates. *Biometrika* **97**:1002–1005.

Conn, P. B., D. S. Johnson, P. J. Williams, S. R. Melin, and M. B. Hooten, 2018. A guide to bayesian model checking for ecologists. *Ecological Monographs* **88**:526–542.

de Valpine, P., D. Turek, C. J. Paciorek, C. Anderson-Bergman, D. T. Lang, and R. Bodik, 2017. Programming with models: writing statistical algorithms for general model structures with nimble. *Journal of Computational and Graphical Statistics* **26**:403–413.

Garrett, E. S. and S. L. Zeger, 2000. Latent class model diagnosis. *Biometrics* **56**:1055–1067.

Gimenez, O., B. J. Morgan, and S. P. Brooks, 2009. Weak identifiability in models for mark-recapture-recovery data. In *Modeling demographic processes in marked populations*, pages 1055–1067. Springer.

Hunter, C. M. and H. Caswell, 2009. Rank and redundancy of multistate mark-recapture models for seabird populations with unobservable states. In D. Thomson, E. Cooch, and M. Conroy, editors, *Modeling Demographic Processes in Marked Populations.*, pages 797–826. Springer series, New York.

Janzen, D., M. Jirstrand, M. J. Chappell, and N. D. Evans, 2016. Three novel approaches to structural identifiability analysis in mixed-effects models. *Computer Methods and Programs in Biomedicine* .

Janzen, D., M. Jirstrand, M. J. Chappell, and N. D. Evans, 2017. Extending existing structural identifiability analysis methods to mixed-effects models. *Mathematical Biosciences* **295**:1–10.

Jiang, H., K. H. Pollock, C. Brownie, J. E. Hightower, J. E. Hoenig, and W. S. Hearn, 2007. Age-dependent tag return models for estimating fishing mortality, natural mortality and selectivity. *Journal of Agricultural, Biological, and Environmental Statistics* **12**:177–194.

MacKenzie, D. I., J. D. Nichols, J. A. Royle, K. H. Pollock, L. L. Bailey, and J. E. Hines, 2005. Occupancy estimation and modeling: inferring patterns and dynamics of species occurrence. Academic Press.

Moe, S. J., C. R. Nater, A. Rustadbakken, L. A. Vøllestad, E. Lund, T. Qvenild, O. Hegge, and P. Aass, 2019. A 50-year series of mark-recapture data of large-sized brown trout (*Salmo trutta*) from Lake Mjøsa, Norway. *bioRxiv* doi:10.1101/544825.

Nater, C. R., A. Rustadbakken, T. Ergon, Ø. Langangen, S. J. Moe, Y. Vindenes, L. A. Vøllestad, and P. Aass, 2018. Individual heterogeneity and early life conditions shape growth in a freshwater top predator. *Ecology* **99**:1011–1017.

NIMBLE Development Team, 2018. NIMBLE: MCMC, particle filtering, and programmable hierarchical modeling. R package version 0.6-13.

Plummer, M., 2003. JAGS: A program for analysis of bayesian graphical models using Gibbs sampling. Version 4.2.0.

R Core Team, 2018. R: A Language and Environment for Statistical Computing. R Foundation for Statistical Computing, Vienna, Austria.

Royle, J. A., R. B. Chandler, C. Yackulic, and J. D. Nichols, 2012. Likelihood analysis of species occurrence probability from presence-only data for modelling species distributions. *Methods in Ecology and Evolution* **3**:545–554.

Sólymos, P., 2010. dclone: Data cloning in R. *The R Journal* **2**:29–37.

Turek, D., P. de Valpine, and C. J. Paciorek, 2016. Efficient Markov chain Monte Carlo sampling for hierarchical hidden Markov models. *Environmental and Ecological Statistics* **23**:549–564.

Turek, D., P. de Valpine, C. J. Paciorek, C. Anderson-Bergman, et al., 2017. Automated parameter blocking for efficient Markov chain Monte Carlo sampling. *Bayesian Analysis* **12**:465–490.
