## Supplementary material for "Size- and stage-dependence in cause-specific mortality of migratory brown trout": Maple code for intrinsic identifiability analyses: constant.pdf

```
> # This Maple code is for the constant example in the Supplementary material of the paper
> # Estimating cause- and size- specific mortality hazard rates using mark-recapture-recovery data
> # by Chloé Nater, Yngvild Vindenesa, Diana Cole, Øystein Langangena, S. Jannicke Moec,
      Daniel Turekd, L. Asbjørn Vøllestada,
  # and Torbjørn Ergona
> restart;
> with(LinearAlgebra) :
> # Derivative matrix proceedure
> Dmat :=proc(se, pars)
  local DD1, i, j;
  description "This procedure finds the derivative matrix for vector se with parameters pars";
  with(LinearAlgebra) :
  DD1 := Matrix(1 ..Dimension(pars), 1 ..Dimension(se)) :
  for i from 1 to Dimension(pars)  do
      for j from 1 to Dimension(se) do
          DD1[i, j] := diff(se[j], pars[i])
      end do
   end do;
  DD1;
   end proc:
> # Estimable combinations proceedure
> Estpar :=proc(DD1, pars, ret)
  local r, d, alphapre, alpha, PDE, FF, i, ans;
  description "Finds the estimable set of parameters for derivative matrix DD1. If ret = 1 returns
      alpha, PDEs, estimable parameter combinations. Otherwise returns estimable parameter
      combinations";
  with(LinearAlgebra) :
  r := Rank(DD1); d := Dimension(pars) - r :
  alphapre := NullSpace(Transpose(DD1)) : α := Matrix(d, Dimension(pars)) : PDE :=
      Vector(d) :
  FF := f(seq(pars[i], i = 1 ..Dimension(pars))) :
  for i from 1 to d do
        α[i, 1 ..Dimension(pars)] := alphapre[i] :
        PDE[i] := add(diff(FF, pars[j])·α[i, j], j = 1 ..Dimension(pars)) :
  end do:
   if ret = 1 then
           ans := ⟨pdsolve({seq(PDE[i] = 0, i = 1 ..d)}), {alpha}, {PDE}⟩ :
   else
          ans := pdsolve({seq(PDE[i] = 0, i = 1 ..d)}) :
   end if:
   ans :
  end proc:
> #Hybrid proceedure :
> Hybrid :=proc(D1, pars, minpars, maxpars, ret)
   local results, j, numpars, D1rand, ans, roll :

      description   "This procedure finds the rank and alpha for the hybrid-symbolic-numeric
      method. If ret = 1 returns full results. Otherwise returns model rank.";
  results := Matrix(5, 2) :
```

```
for j from 1 to 5 do
   roll := rand(minpars..maxpars) :
   numpars := seq(pars[i] = evalf(roll( )), i = 1..Dimension(pars)) :
   D1rand := eval(D1, {numpars});
   results[j, 1] := Rank( D1rand);
  results[j, 2] := NullSpace(Transpose(D1rand)) :
end do:
 if ret = 1 then
      ans := results :
else
     ans := max(results[1..5, 1]) :
end if:
ans :
end proc:
```

```
> # Set up Matrices:
> #Initial state matrix:
> Pit := ⟨1|0|0|0⟩;
```

$$Pit := \begin{bmatrix} 1 & 0 & 0 & 0 \end{bmatrix} \quad (1)$$

```
> # Transition matrix:
> Phit := ⟨⟨S[1]·p[1]|S[1]·(1 − p[1])|(1 − S[1])·alpha[1]|(1 − S[1])·(1 − alpha[1])⟩,
    ⟨S[2]·p[2]|S[2]·(1 − p[2])|(1 − S[2])·alpha[2]|(1 − S[2])·(1 − alpha[2])⟩, ⟨0|0|0
    |1⟩, ⟨0|0|0|1⟩⟩;
```

$$Phit := \begin{bmatrix} S_1 p_1 & S_1 (1 - p_1) & (1 - S_1)\,\alpha_1 & (1 - S_1)(1 - \alpha_1) \\ S_2 p_2 & S_2 (1 - p_2) & (1 - S_2)\,\alpha_2 & (1 - S_2)(1 - \alpha_2) \\ 0 & 0 & 0 & 1 \\ 0 & 0 & 0 & 1 \end{bmatrix} \quad (2)$$

```
> # Observation matrix:
> B := ⟨⟨1|0|0⟩, ⟨0|0|1⟩, ⟨0|lambda|1 − lambda⟩, ⟨0|0|1⟩⟩;
```

$$B := \begin{bmatrix} 1 & 0 & 0 \\ 0 & 0 & 1 \\ 0 & \lambda & 1 - \lambda \\ 0 & 0 & 1 \end{bmatrix} \quad (3)$$

```
> B := Transpose(B);
```

$$B := \begin{bmatrix} 1 & 0 & 0 & 0 \\ 0 & 0 & \lambda & 0 \\ 0 & 1 & 1 - \lambda & 1 \end{bmatrix} \quad (4)$$

```
> # First time matrices
> B0 := B :
> Phit0 := Phit :
>
> # T number of years (N +1 number of observation options, here 1,2 or 3)
```

```
> # Finds every possible capture history. Removes histories that are not possible e.g. 2 deaths
> T := 5 : N := 2 :
H2 := Matrix( (N + 1)^T, T) :
for i from 1 to (N + 1)^T do
    nextone := i - 1 :
    for j from 1 to T do
        H2[i, j] := nextone mod (N + 1) + 1 :
        nextone := trunc( nextone / (N + 1) ) :
    end do:
 end do:
H3 := Matrix(1, Dimension(H2)[2]) :
 for i from 1 to Dimension(H2)[1] do
    no2 := 0 :
    for j from 1 to Dimension(H2)[2] do
        if H2[i, j] = 2 then
            no2 := no2 + 1 :
        end if:
    end do:
    if no2 ≤ 1 then
        H3 := ⟨H3, H2[i, 1 ..Dimension(H2)[2]]⟩ :
    end if:
 end do:
H3 := H3[2 ..Dimension(H3)[1], 1 ..Dimension(H3)[2]] :
H := Matrix(1, Dimension(H3)[2]) :
for i from 1 to Dimension(H3)[1] do
     index2 := Dimension(H3)[2] + 1 :
     for j from 1 to Dimension(H3)[2] do
         if H3[i, j] = 2 then
             index2 := j :
         end if:
    end do:
     index1 := 0 :
    firstyes := 0 :
     for j from Dimension(H3)[2] by -1 to 1 do
         if firstyes = 0 then
             if H3[i, j] = 1 then
                 index1 := j :
                 firstyes := 1 :
             end if:
         end if:
    end do:
   if index1 > 0 then
    if index2 > index1 then
        H := ⟨H, H3[i, 1 ..Dimension(H3)[2]]⟩ :
   end if:
 end if:
 end do:
H := H[2 ..Dimension(H)[1], 1 ..Dimension(H)[2]] :
```

> # Finds the probability of each history and creates the exhaustive summary of probabilities, κ

```
> κ := Vector(Dimension(H)[1]) :
for j from 1 to Dimension(H)[1] do
    h := Row(H, j) :
    # e is first nonzero entry of h
   indi := 0 :
   for k from 1 to Dimension(H)[2] do
       if indi = 0 then
          if h[k] = 1 then
             e := k :
             indi := 1 :
          end if:
       end if:
   end do:
   Ph := Multiply(eval(Pit, t = e), DiagonalMatrix(Row(B0, h[e])));
   for i from e + 1 to T do
        if i − 1 = e then
          Ph := Multiply(Ph, Multiply(eval(Phit0, t = i − 1), DiagonalMatrix(Row(eval(B, t
    = i), h[i])))) :
       else
         Ph := Multiply(Ph, Multiply(eval(Phit, t = i − 1), DiagonalMatrix(Row(eval(B, t = i),
   h[i])))) :
       end if:
    end do:
    Ph := Multiply(Ph, Vector(Dimension(Ph), 1));
    κ[j] := Ph :
end do:
```

> # Evaluate the exhaustive summary at the parameterisation used for this model:

> $kappa2 := eval\left(\text{kappa}, \left\{S[1] = \exp(-(mH + mO[1])), S[2] = \exp(-(mH + mO[2])), \text{alpha}[1] = \frac{mH}{mH + mO[1]}, \text{alpha}[2] = \frac{mH}{mH + mO[2]}\right\}\right);$

$$\kappa 2 := \left[\begin{array}{c} \text{1 .. 57 } Vector_{column} \\ \text{Data Type: anything} \\ \text{Storage: rectangular} \\ \text{Order: Fortran\_order} \end{array}\right] \quad (5)$$

> $indets(\text{kappa})$

$$\{\lambda, S_1, S_2, \alpha_1, \alpha_2, p_1, p_2\} \quad (6)$$

> $indets(kappa2)$

$$\left\{\lambda, mH, mO_1, mO_2, p_1, p_2, e^{-mH - mO_1}, e^{-mH - mO_2}\right\} \quad (7)$$

> # Vector of parameters (note we use the parameter λ rather than r for reporting rate)

> $pars := \langle \lambda, mH, mO_1, mO_2, p_1, p_2 \rangle;$

$$pars := \begin{bmatrix} \lambda \\ mH \\ mO_1 \\ mO_2 \\ p_1 \\ p_2 \end{bmatrix} \quad \textbf{(8)}$$

> # D1 stores the derivative matrix which is calculated using proceedure Dmat

> $D1 := Dmat(kappa2, pars)$ :

> # r stores the model rank, which is calculated using Maple's intrinsic proceedure rank. d stores the deficency, the difference between the number of parameters and the rank. A deficency of one or more indicates the model is parameter redundant, so this model is parameter redundant.

> $r := Rank(D1); d := Dimension(pars) - r;$

$$r := 5$$

$$d := 1 \quad \textbf{(9)}$$

> # Hybrid Symbolic-Numerical Method

> $Hybrid(D1, pars, 0.0, 1.0, 1)$

**(10)**

$$
\left[\begin{array}{cc}
5 & \left\{\left[\begin{array}{c} -0.317416856268040 \\ 0.547493237994324 \\ -0.547493237730089 \\ -0.547493240458025 \\ 1.44851459182392\ 10^{-10} \\ -6.98539493725700\ 10^{-10} \end{array}\right]\right\} \\
5 & \left\{\left[\begin{array}{c} 0.292860175764446 \\ -0.552036508761466 \\ 0.552036509092366 \\ 0.552036505201226 \\ 2.93463370147742\ 10^{-10} \\ 1.56610334604372\ 10^{-10} \end{array}\right]\right\} \\
5 & \left\{\left[\begin{array}{c} 0.0585809492149168 \\ -0.576358765051083 \\ 0.576358767511340 \\ 0.576358757589992 \\ 7.16310395564949\ 10^{-11} \\ -2.68693052244871\ 10^{-9} \end{array}\right]\right\} \\
5 & \left\{\left[\begin{array}{c} -0.288114167587521 \\ 0.552868346597555 \\ -0.552868347226462 \\ -0.552868346354642 \\ -2.64376827155999\ 10^{-10} \\ -7.46512837790941\ 10^{-10} \end{array}\right]\right\} \\
5 & \left\{\left[\begin{array}{c} -0.206782149668881 \\ 0.564872003692989 \\ -0.564872001479667 \\ -0.564872006711731 \\ 3.66617588921086\ 10^{-10} \\ -2.34266698206422\ 10^{-10} \end{array}\right]\right\}
\end{array}\right] \tag{10}
$$

>

> # It is also possible to find estimable parameter combinations in non-identifiable models.

> *simplify*(*Estpar*(*D1*, *pars*, 0));

[illegible]

$$\left\{ f\left(\lambda, mH, mO_1, mO_2, p_1, p_2\right) = \_F1\left(\lambda\, mH, mH + mO_1, mH + mO_2, p_1, p_2\right) \right\} \quad \textbf{(11)}$$

> #The estimable parameter combinations are : $\lambda\, mH,\ mH + mO_1,\ mH + mO_2,\ p_1,\ p_2$

>
