## Supplementary material for "Size- and stage-dependence in cause-specific mortality of migratory brown trout": Maple code for intrinsic identifiability analyses: covariates.pdf

```
# This Maple code is for the covariate example in the Supplementary material of the paper
# Estimating cause- and size- specific mortality hazard rates using mark-recapture-recovery data
# by Chloé Nater, Yngvild Vindenesa, Diana Cole, Øystein Langangena, S. Jannicke Moec,
        Daniel Turekd, L. Asbjørn Vøllestada,
 and Torbjørn Ergona

restart;
with(LinearAlgebra) :
# Derivative matrix proceedure
Dmat := proc(se, pars)
local DD1, i, j;
description "This procedure finds the derivative matrix for vector se with parameters pars";
with(LinearAlgebra) :
DD1 := Matrix(1 ..Dimension(pars), 1 ..Dimension(se)) :
for i from 1 to Dimension(pars) do
    for j from 1 to Dimension(se) do
        DD1[i, j] := diff(se[j], pars[i])
    end do
 end do;
DD1;
 end proc:
# Estimable combinations proceedure
Estpar := proc(DD1, pars, ret)
local r, d, alphapre, alpha, PDE, FF, i, ans;
description "Finds the estimable set of parameters for derivative matrix DD1. If ret = 1 returns
    alpha, PDEs, estimable parameter combinations. Otherwise returns estimable parameter
    combinations";
with(LinearAlgebra) :
r := Rank(DD1); d := Dimension(pars) - r :
alphapre := NullSpace(Transpose(DD1)) : α := Matrix(d, Dimension(pars)) : PDE :=
    Vector(d) :
FF := f(seq(pars[i], i = 1 ..Dimension(pars))) :
for i from 1 to d do
    α[i, 1 ..Dimension(pars)] := alphapre[i] :
    PDE[i] := add(diff(FF, pars[j]) ·α[i, j], j = 1 ..Dimension(pars)) :
end do:
 if ret = 1 then
        ans := ⟨pdsolve({seq(PDE[i] = 0, i = 1 ..d)}), {alpha}, {PDE}⟩ :
 else
        ans := pdsolve({seq(PDE[i] = 0, i = 1 ..d)}) :
 end if:
 ans :
end proc:
#` Hybrid proceedure :
Hybrid := proc(D1, pars, minpars, maxpars, ret)
 local results, j, numpars, D1rand, ans, roll :

    description "This procedure finds the rank and alpha for the hybrid-symbolic-numeric
    method. If ret = 1 returns full results. Otherwise returns model rank.";
```

```
results := Matrix(5, 2) :
 for j from 1 to 5 do
     roll := rand(minpars ..maxpars) :
    numpars := seq(pars[i] = evalf(roll( )), i = 1 ..Dimension(pars)) :
    D1rand := eval(D1, {numpars});
    results[j, 1] := Rank( D1rand);
   results[j, 2] := NullSpace(Transpose(D1rand)) :
 end do:
  if ret = 1 then
          ans := results :
 else
         ans := max(results[1 ..5, 1]) :
 end if:
 ans :
 end proc:
> exsumtrout :=proc(A, T)
 local i, j, k, kappa, kappaindex;
description "Finds the exhaustive summary for the B model";
with(LinearAlgebra) :
 kappa := Vector(5·T − 9) :
 kappaindex := 1 :
 for i from 1 to T − 1 do
       kappa[kappaindex] := eval(A[1, 1], t = i) :
      kappaindex := kappaindex + 1;
 end do:
 for i from 2 to T − 1 do
       kappa[kappaindex] := eval(A[1, 2], t = i − 1) ·eval(A[2, 1], t = i) :
      kappaindex := kappaindex + 1;
 end do:
 for i from 1 to T − 1 do
       kappa[kappaindex] := eval(A[1, 3], t = i) ·lambda[i + 1] :
      kappaindex := kappaindex + 1;
 end do:
 for i from 2 to T − 1 do
       kappa[kappaindex] := (eval(A[1, 2], t = i − 1) ·eval(A[2, 3], t = i)) / (eval(A[1, 3], t = i)) :
      kappaindex := kappaindex + 1;
 end do:
 for i from 2 to T − 2 do
       kappa[kappaindex] := (eval(A[1, 2], t = i − 1) ·eval(A[2, 2], t = i)) / (eval(A[1, 2], t = i)) :
      kappaindex := kappaindex + 1;
 end do:

 kappa :
end proc:
>
>  # Setting up transition matrix:
> Phit := ⟨⟨S[t, 1]·p[t + 1]|S[t, 1]· (1 − p[t + 1])|(1 − S[t, 1]) ·alpha[t, 1]|(1 − S[t, 1]) · (1
       − alpha[t, 1])⟩, ⟨S[t, 2]·p[t + 1]|S[t, 2]· (1 − p[t + 1])|(1 − S[t, 2]) ·alpha[t, 2]|(1
       − S[t, 2]) · (1 − alpha[t, 2])⟩, ⟨0|0|0|1⟩, ⟨0|0|0|1⟩⟩;
```

$$Phit := \begin{bmatrix} S_{t,1}\,p_{t+1} & S_{t,1}\left(1-p_{t+1}\right) & \left(1-S_{t,1}\right)\alpha_{t,1} & \left(1-S_{t,1}\right)\left(1-\alpha_{t,1}\right) \\ S_{t,2}\,p_{t+1} & S_{t,2}\left(1-p_{t+1}\right) & \left(1-S_{t,2}\right)\alpha_{t,2} & \left(1-S_{t,2}\right)\left(1-\alpha_{t,2}\right) \\ 0 & 0 & 0 & 1 \\ 0 & 0 & 0 & 1 \end{bmatrix} \quad \textbf{(1)}$$

> # Changing transition matrix to parameterisation involving mH and MO etc. mH dependents on an individual covariate, mO depends on the same individual covariate as well as a time varying covariate, p depends on the same individual covariate and a different time varying covariate.

> # Note that the link function used does not effect the identifiability result, therefore rather than a log link, we use the simplier linear link function in calculation.

> $Phit2 := eval\Big(eval\Big(Phit, \Big\{S[t,1] = \exp(-mH[t] - mO[t,1]), S[t,2] = \exp(-mH[t] - mO[t,2]), \text{alpha}[t,1] = \frac{mH[t]}{mH[t] + mO[t,1]}, \text{alpha}[t,2] = \frac{mH[t]}{mH[t] + mO[t,2]}\Big\}\Big), \{mH[t] = bH[0] + bH[1]\cdot x1[ii,t], mO[t,1] = bM1[0] + bM1[1]\cdot x1[ii,t] + bM1[2]\cdot x2[t], mO[t,2] = bM2[0] + bM2[1]\cdot x1[ii,t] + bM2[2]\cdot x2[t], p[t+1] = bp[0] + bp[1]\cdot x1[ii,t+1] + bp[2]\cdot x3[t+1]\}\Big);$

$$Phit2 := \Big[\Big[ e^{-bH_1\,x1_{ii,t} - bM1_1\,x1_{ii,t} - bM1_2\,x2_t - bH_0 - bM1_0}\left(bp_1\,x1_{ii,t+1} + bp_2\,x3_{t+1} + bp_0\right), \quad \textbf{(2)}$$

$$e^{-bH_1\,x1_{ii,t} - bM1_1\,x1_{ii,t} - bM1_2\,x2_t - bH_0 - bM1_0}\left(-bp_1\,x1_{ii,t+1} - bp_2\,x3_{t+1} - bp_0 + 1\right),$$

$$\frac{\left(1 - e^{-bH_1\,x1_{ii,t} - bM1_1\,x1_{ii,t} - bM1_2\,x2_t - bH_0 - bM1_0}\right)\left(bH_1\,x1_{ii,t} + bH_0\right)}{bH_1\,x1_{ii,t} + bM1_1\,x1_{ii,t} + bM1_2\,x2_t + bH_0 + bM1_0}, \Big(1$$

$$- e^{-bH_1\,x1_{ii,t} - bM1_1\,x1_{ii,t} - bM1_2\,x2_t - bH_0 - bM1_0}\Big)\Big(1$$

$$- \frac{bH_1\,x1_{ii,t} + bH_0}{bH_1\,x1_{ii,t} + bM1_1\,x1_{ii,t} + bM1_2\,x2_t + bH_0 + bM1_0}\Big)\Big],$$

$$\Big[ e^{-bH_1\,x1_{ii,t} - bM2_1\,x1_{ii,t} - bM2_2\,x2_t - bH_0 - bM2_0}\left(bp_1\,x1_{ii,t+1} + bp_2\,x3_{t+1} + bp_0\right),$$

$$e^{-bH_1\,x1_{ii,t} - bM2_1\,x1_{ii,t} - bM2_2\,x2_t - bH_0 - bM2_0}\left(-bp_1\,x1_{ii,t+1} - bp_2\,x3_{t+1} - bp_0 + 1\right),$$

$$\frac{\left(1 - e^{-bH_1\,x1_{ii,t} - bM2_1\,x1_{ii,t} - bM2_2\,x2_t - bH_0 - bM2_0}\right)\left(bH_1\,x1_{ii,t} + bH_0\right)}{bH_1\,x1_{ii,t} + bM2_1\,x1_{ii,t} + bM2_2\,x2_t + bH_0 + bM2_0}, \Big(1$$

$$- e^{-bH_1 x1_{ii,t} - bM2_1 x1_{ii,t} - bM2_2 x2_t - bH_0 - bM2_0} \left( 1 - \frac{bH_1 x1_{ii,t} + bH_0}{bH_1 x1_{ii,t} + bM2_1 x1_{ii,t} + bM2_2 x2_t + bH_0 + bM2_0} \right) \Bigg],$$

$$\Big[ 0, 0, 0, 1 \Big],$$

$$\Big[ 0, 0, 0, 1 \Big]\Big]$$

```
> T := 5 :
> # Here we assume the reporting probability is constant. Below the reporting probability is
      represented by lambda rather than r.
> kappa := eval(convert(⟨eval(exsumtrout(Phit2, T), ii = 1),
         eval(exsumtrout(Phit2, T), ii = 2)⟩, Vector), {seq(lambda[i] = lambda, i = 2 ..T)})
      :
      indets(kappa, name)
```

$$\{\lambda, bH_0, bH_1, bM1_0, bM1_1, bM1_2, bM2_0, bM2_1, bM2_2, bp_0, bp_1, bp_2, x1_{1,1}, x1_{1,2}, x1_{1,3}, x1_{1,4}, x1_{1,5}, x1_{2,1}, x1_{2,2}, x1_{2,3}, x1_{2,4}, x1_{2,5}, x2_1, x2_2, x2_3, x2_4, x3_2, x3_3, x3_4, x3_5\} \quad \textbf{(3)}$$

```
> pars := ⟨λ, bH0, bH1, bM10, bM11, bM12, bM20, bM21, bM22, bp0, bp1, bp2⟩ :
> D1 := Dmat(kappa, pars) :
> indetpars := ⟨seq(indets(kappa)[i], i = 1 ..nops(indets(kappa, name)))⟩ :
> r := Hybrid(D1, indetpars, 0.0, 1.0, 0); d := Dimension(pars) − r;
```

$$r := 11$$

$$d := 1 \quad \textbf{(4)}$$

```
> T := 6 :
> # Here we assume the reporting probability is constant. Below the reporting probability is
      represented by lambda rather than r.
> kappa := eval(convert(⟨eval(exsumtrout(Phit2, T), ii = 1),
         eval(exsumtrout(Phit2, T), ii = 2)⟩, Vector), {seq(lambda[i] = lambda, i = 2 ..T)})
      :
      indets(kappa, name)
```

$$\{\lambda, bH_0, bH_1, bM1_0, bM1_1, bM1_2, bM2_0, bM2_1, bM2_2, bp_0, bp_1, bp_2, x1_{1,1}, x1_{1,2}, x1_{1,3}, x1_{1,4}, x1_{1,5}, x1_{1,6}, x1_{2,1}, x1_{2,2}, x1_{2,3}, x1_{2,4}, x1_{2,5}, x1_{2,6}, x2_1, x2_2, x2_3, x2_4, x2_5, x3_2, x3_3, x3_4, x3_5, x3_6\} \quad \textbf{(5)}$$

```
> pars := ⟨λ, bH0, bH1, bM10, bM11, bM12, bM20, bM21, bM22, bp0, bp1, bp2⟩ :
> D1 := Dmat(kappa, pars) :
> indetpars := ⟨seq(indets(kappa)[i], i = 1 ..nops(indets(kappa, name)))⟩ :
> r := Hybrid(D1, indetpars, 0.0, 1.0, 0); d := Dimension(pars) − r;
```

$$r := 11$$

$$d := 1 \quad \textbf{(6)}$$

>

> # Time dependent reporting probability

> $T := 5:$

> $\text{kappa} := convert(\langle eval(exsumtrout(Phit2, T), ii=1), eval(exsumtrout(Phit2, T), ii=2)\rangle, Vector):$
$indets(\text{kappa}, name)$

$$\{bH_0, bH_1, bM1_0, bM1_1, bM1_2, bM2_0, bM2_1, bM2_2, bp_0, bp_1, bp_2, \lambda_2, \lambda_3, \lambda_4, \lambda_5, x1_{1,1}, x1_{1,2}, x1_{1,3}, x1_{1,4}, x1_{1,5}, x1_{2,1}, x1_{2,2}, x1_{2,3}, x1_{2,4}, x1_{2,5}, x2_1, x2_2, x2_3, x2_4, x3_2, x3_3, x3_4, x3_5\} \tag{7}$$

> $pars := \langle bH_0, bH_1, bM1_0, bM1_1, bM1_2, bM2_0, bM2_1, bM2_2, bp_0, bp_1, bp_2, \lambda_2, \lambda_3, \lambda_4, \lambda_5\rangle :$

> $D1 := Dmat(\text{kappa}, pars):$

> $indetpars := \langle seq(indets(\text{kappa})[i], i=1..nops(indets(\text{kappa}, name)))\rangle :$

> $r := Hybrid(D1, indetpars, 0.0, 1.0, 0);\ d := Dimension(pars) - r;$

$$r := 14$$
$$d := 1 \tag{8}$$

> $T := 6:$

> $\text{kappa} := convert(\langle eval(exsumtrout(Phit2, T), ii=1), eval(exsumtrout(Phit2, T), ii=2)\rangle, Vector):$
$indets(\text{kappa}, name)$

$$\{bH_0, bH_1, bM1_0, bM1_1, bM1_2, bM2_0, bM2_1, bM2_2, bp_0, bp_1, bp_2, \lambda_2, \lambda_3, \lambda_4, \lambda_5, \lambda_6, x1_{1,1}, x1_{1,2}, x1_{1,3}, x1_{1,4}, x1_{1,5}, x1_{1,6}, x1_{2,1}, x1_{2,2}, x1_{2,3}, x1_{2,4}, x1_{2,5}, x1_{2,6}, x2_1, x2_2, x2_3, x2_4, x2_5, x3_2, x3_3, x3_4, x3_5, x3_6\} \tag{9}$$

> $pars := \langle bH_0, bH_1, bM1_0, bM1_1, bM1_2, bM2_0, bM2_1, bM2_2, bp_0, bp_1, bp_2, \lambda_2, \lambda_3, \lambda_4, \lambda_5, \lambda_6\rangle :$

> $D1 := Dmat(\text{kappa}, pars):$

> $indetpars := \langle seq(indets(\text{kappa})[i], i=1..nops(indets(\text{kappa}, name)))\rangle :$

> $r := Hybrid(D1, indetpars, 0.0, 1.0, 0);\ d := Dimension(pars) - r;$

$$r := 15$$
$$d := 1 \tag{10}$$

>

>
