## Supplementary material for "Size- and stage-dependence in cause-specific mortality of migratory brown trout": Maple code for intrinsic identifiability analyses: randomeffects.pdf

```
> # Parameter Redundancy in Trout CMRR model with time simpler exhautive summary
> # This Maple code is for the random effects example in the Supplementary material of the paper
> # Estimating cause- and size- specific mortality hazard rates using mark-recapture-recovery data
> # by Chloé Nater, Yngvild Vindenesa, Diana Cole, Øystein Langangena, S. Jannicke Moec,
      Daniel Turekd, L. Asbjørn Vøllestada,
  # and Torbjørn Ergona
> restart;
> with(LinearAlgebra) :
> # Derivative matrix proceedure
> Dmat :=proc(se, pars)
  local DD1, i, j;
  description "This procedure finds the derivative matrix for vector se with parameters pars";
  with(LinearAlgebra) :
  DD1 := Matrix(1 ..Dimension(pars), 1 ..Dimension(se)) :
  for i from 1 to Dimension(pars)  do
      for j from 1 to Dimension(se) do
          DD1[i, j] := diff(se[j], pars[i])
      end do
   end do;
  DD1;
   end proc:
> # Estimable combinations proceedure
> Estpar :=proc(DD1, pars, ret)
  local r, d, alphapre, alpha, PDE, FF, i, ans;
  description "Finds the estimable set of parameters for derivative matrix DD1. If ret = 1 returns
      alpha, PDEs, estimable parameter combinations. Otherwise returns estimable parameter
      combinations";
  with(LinearAlgebra) :
  r := Rank(DD1); d := Dimension(pars) - r :
  alphapre := NullSpace(Transpose(DD1)) : α := Matrix(d, Dimension(pars)) : PDE :=
      Vector(d) :
  FF := f(seq(pars[i], i = 1 ..Dimension(pars))) :
  for i from 1 to d do
        α[i, 1 ..Dimension(pars)] := alphapre[i] :
        PDE[i] := add(diff(FF, pars[j])·α[i, j], j = 1 ..Dimension(pars)) :
  end do:
   if ret = 1 then
          ans := ⟨pdsolve({seq(PDE[i] = 0, i = 1 ..d)}), {alpha}, {PDE}⟩ :
   else
          ans := pdsolve({seq(PDE[i] = 0, i = 1 ..d)}) :
   end if:
   ans :
  end proc:
> # Hybrid proceedure :
> Hybrid :=proc(D1, pars, minpars, maxpars, ret)
   local results, j, numpars, D1rand, ans, roll :

      description   "This procedure finds the rank and alpha for the hybrid-symbolic-numeric
      method. If ret = 1 returns full results. Otherwise returns model rank.";
```

```
results := Matrix(5, 2) :
for j from 1 to 5 do
    roll := rand(minpars ..maxpars) :
    numpars := seq(pars[i] = evalf(roll( )), i = 1 ..Dimension(pars)) :
    D1rand := eval(D1, {numpars});
    results[j, 1] := Rank( D1rand);
   results[j, 2] := NullSpace(Transpose(D1rand)) :
end do:
 if ret = 1 then
        ans := results :
else
        ans := max(results[1 ..5, 1]) :
end if:
ans :
end proc:
> exsumtrout :=proc(A, T)
local i, j, k, kappa, kappaindex;
description "Finds the exhaustive summary for the B model";
with(LinearAlgebra) :
kappa := Vector(5·T − 9) :
kappaindex := 1 :
for i from 1 to T − 1 do
     kappa[kappaindex] := eval(A[1, 1], t = i) :
     kappaindex := kappaindex + 1;
end do:
for i from 2 to T − 1 do
     kappa[kappaindex] := eval(A[1, 2], t = i − 1) ·eval(A[2, 1], t = i) :
     kappaindex := kappaindex + 1;
end do:
for i from 1 to T − 1 do
     kappa[kappaindex] := eval(A[1, 3], t = i) ·lambda[i + 1] :
     kappaindex := kappaindex + 1;
end do:
for i from 2 to T − 1 do
     kappa[kappaindex] := (eval(A[1, 2], t = i − 1) ·eval(A[2, 3], t = i)) / (eval(A[1, 3], t = i)) :
     kappaindex := kappaindex + 1;
end do:
for i from 2 to T − 2 do
     kappa[kappaindex] := (eval(A[1, 2], t = i − 1) ·eval(A[2, 2], t = i)) / (eval(A[1, 2], t = i)) :
     kappaindex := kappaindex + 1;
end do:

kappa :
end proc:
>
> # Parameter constraints (note lambda is r in the paper)
> mHconst := bH[0] + bH[1]·x1[ii, t] + refM[t] :
> mOconst1 := bM1[0] + bM1[1]·x1[ii, t] + bM1[2]·x2[t] :
> mOconst2 := bM2[0] + bM2[1]·x1[ii, t] + bM2[2]·x2[t] :
```

> $pconst1 := bp[0] + bp[1] \cdot x1[ii, t+1] + bp[2] \cdot x3[t+1]$ :

> $pconst2 := pconst1$ :

> $lambdaconst := \text{lambda}$ :

>

> # Transition Matrix.

> $Phit := \langle \langle S[t,1] \cdot p[t+1,1] | S[t,1] \cdot (1 - p[t+1,1]) | (1 - S[t,1]) \cdot \text{alpha}[t,1] | (1 - S[t,1]) \cdot (1 - \text{alpha}[t,1]) \rangle, \langle S[t,2] \cdot p[t+1,2] | S[t,2] \cdot (1 - p[t+1,2]) | (1 - S[t,2]) \cdot \text{alpha}[t,2] | (1 - S[t,2]) \cdot (1 - \text{alpha}[t,2]) \rangle, \langle 0|0|0|1 \rangle, \langle 0|0|0|1 \rangle \rangle$ :

> $Phit2 := eval\left( eval\left( Phit, \left\{ S[t,1] = \exp(-mH[t] - mO[t,1]), S[t,2] = \exp(-mH[t] - mO[t,2]), \text{alpha}[t,1] = \frac{mH[t]}{mH[t] + mO[t,1]}, \text{alpha}[t,2] = \frac{mH[t]}{mH[t] + mO[t,2]} \right\} \right), \{ mH[t] = mHconst, mO[t,1] = mOconst1, mO[t,2] = mOconst2, p[t+1,1] = pconst1, p[t+1,2] = pconst2 \} \right)$ :

>

> $T := 5$ :

>

> # Exhaustive summary consists of two repeats of the exhaustive summary with diffferent random effect values. Here we ignore the random effect commonent

> $kappae1 := eval(eval(eval(exsumtrout(Phit2, T), \{seq(\text{lambda}[iii] = lambdaconst, iii = 1..T)\}), \{seq(refH[i] = RandomMatrix(1, generator = 0.0..1.0)[1,1], i = 1..T), seq(refM[i] = RandomMatrix(1, generator = 0.0..1.0)[1,1], i = 1..T), seq(refP[i] = RandomMatrix(1, generator = 0.0..1.0)[1,1], i = 1..T), seq(refL[i] = RandomMatrix(1, generator = 0.0..1.0)[1,1], i = 1..T)\}), ii = 1)$ :

> $kappae2 := eval(eval(eval(exsumtrout(Phit2, T), \{seq(\text{lambda}[iii] = lambdaconst, iii = 1..T)\}), \{seq(refH[i] = RandomMatrix(1, generator = 0.0..1.0)[1,1], i = 1..T), seq(refM[i] = RandomMatrix(1, generator = 0.0..1.0)[1,1], i = 1..T), seq(refP[i] = RandomMatrix(1, generator = 0.0..1.0)[1,1], i = 1..T), seq(refL[i] = RandomMatrix(1, generator = 0.0..1.0)[1,1], i = 1..T)\}), ii = 2)$ :

> $\text{kappa} := convert(\langle kappae1, kappae2 \rangle, Vector)$ :

>

> $seq(indets(\text{kappa}, name)[i], i = 1..nops(indets(\text{kappa}, name)))$;

$$\lambda, bH_0, bH_1, bM1_0, bM1_1, bM1_2, bM2_0, bM2_1, bM2_2, bp_0, bp_1, bp_2, x1_{1,1}, x1_{1,2}, x1_{1,3}, x1_{1,4}, x1_{1,5}, x1_{2,1}, x1_{2,2}, x1_{2,3}, x1_{2,4}, x1_{2,5}, x2_1, x2_2, x2_3, x2_4, x3_2, x3_3, x3_4, x3_5 \quad \textbf{(1)}$$

> $pars := \langle \lambda, bH_0, bH_1, bM1_0, bM1_1, bM1_2, bM2_0, bM2_1, bM2_2, bp_0, bp_1, bp_2 \rangle$ :

> $D1 := Dmat(\text{kappa}, pars)$ :

>

> $indetpars := \langle seq(indets(\text{kappa}, name)[i], i = 1..nops(indets(\text{kappa}, name))) \rangle$ :

> $r := Hybrid(D1, indetpars, 0.0, 1.0, 0)$; $d := Dimension(pars) - r$;

$$r := 12$$

$$d := 0 \quad \textbf{(2)}$$

>

> # The exhaustive summary would then have this additional term for the random effect component

> $kappaex := \left\langle \frac{1}{\text{sigma}} \exp\left( -\frac{1}{2} \left( \frac{bb}{\text{sigma}} \right)^2 \right) \right\rangle$;

$$kappaex := \left[ \frac{e^{-\frac{bb^2}{2\sigma^2}}}{\sigma} \right] \tag{3}$$

> # It is obvious that sigma is identifiable from this term.

>

>
