## Supplementary material for "Size- and stage-dependence in cause-specific mortality of migratory brown trout": Maple code for intrinsic identifiability analyses: readme.docx

This Maple code corresponds to the files mentioned in the Section S3 in the supplementary material of the paper ‘Size- and stage-dependence in cause-specific mortality of migratory brown trout’ by Chloé R. Nater, Yngvild Vindenes, Per Aass, Diana Cole, Øystein Langangen, S. Jannicke Moe, Atle Rustadbakken, Daniel Turek, L. Asbjørn Vøllestad and Torbjørn Ergon

The Maple code is files ending with .mw. A pdf version of the code is also provided, which does not require Maple to read.

| Maple file | pdf file | Explanation |
| --- | --- | --- |
| constant.mw | constant.pdf | Maple code for checking identifiability in the constant model |
| covariates.mw | covariates.mw | Maple code for checking identifiability in models with covariates |
| randomeffect.mw | randomeffect.pdf | Maple code for checking identifiability in models with random effects |
| simplerexproof.mw | simplerexproof.pdf | Maple code that outlines the proof for the simpler exhaustive summary |
