## Supplementary material for "Size- and stage-dependence in cause-specific mortality of migratory brown trout": Maple code for intrinsic identifiability analyses: simplerexproof.pdf

```
> # This Maple code contains the proof of the simpler exhaustive summary
> # Estimating cause- and size- specific mortality hazard rates using mark-recapture-recovery data
> # by Chloé Nater, Yngvild Vindenesa, Diana Cole, Øystein Langangena, S. Jannicke Moec,
      Daniel Turekd, L. Asbjørn Vøllestada,
  # and Torbjørn Ergona
>
> restart;
> with(LinearAlgebra) :
> # Derivative matrix proceedure
> Dmat := proc(se, pars)
  local DD1, i, j;
  description "This procedure finds the derivative matrix for vector se with parameters pars";
  with(LinearAlgebra) :
  DD1 := Matrix(1 ..Dimension(pars), 1 ..Dimension(se)) :
  for i from 1 to Dimension(pars) do
      for j from 1 to Dimension(se) do
          DD1[i, j] := diff(se[j], pars[i])
      end do
   end do;
  DD1;
   end proc:
> # Estimable combinations proceedure
> Estpar := proc(DD1, pars, ret)
  local r, d, alphapre, alpha, PDE, FF, i, ans;
  description "Finds the estimable set of parameters for derivative matrix DD1. If ret = 1 returns
      alpha, PDEs, estimable parameter combinations. Otherwise returns estimable parameter
      combinations";
  with(LinearAlgebra) :
  r := Rank(DD1); d := Dimension(pars) - r :
  alphapre := NullSpace(Transpose(DD1)) : α := Matrix(d, Dimension(pars)) : PDE :=
      Vector(d) :
  FF := f(seq(pars[i], i = 1 ..Dimension(pars))) :
  for i from 1 to d do
        α[i, 1 ..Dimension(pars)] := alphapre[i] :
        PDE[i] := add(diff(FF, pars[j]) · α[i, j], j = 1 ..Dimension(pars)) :
  end do:
   if ret = 1 then
          ans := ⟨pdsolve({seq(PDE[i] = 0, i = 1 ..d)}), {alpha}, {PDE}⟩ :
   else
          ans := pdsolve({seq(PDE[i] = 0, i = 1 ..d)}) :
   end if:
   ans :
  end proc:
> # Hybrid proceedure :
> Hybrid := proc(D1, pars, minpars, maxpars, ret)
   local results, j, numpars, D1rand, ans, roll :

      description "This procedure finds the rank and alpha for the hybrid-symbolic-numeric
      method. If ret = 1 returns full results. Otherwise returns model rank.";
```

```
results := Matrix(5, 2) :
 for j from 1 to 5 do
     roll := rand(minpars ..maxpars) :
    numpars := seq(pars[i] = evalf(roll( )), i = 1 ..Dimension(pars)) :
    D1rand := eval(D1, {numpars});
    results[j, 1] := Rank( D1rand);
   results[j, 2] := NullSpace(Transpose(D1rand)) :
 end do:
  if ret = 1 then
         ans := results :
 else
         ans := max(results[1 ..5, 1]) :
 end if:
 ans :
 end proc:
> # Set up matrices:
> # Initial state matrix:
> Pit := ⟨1|0|0|0⟩;
```

$$Pit := \begin{bmatrix} 1 & 0 & 0 & 0 \end{bmatrix} \tag{1}$$

```
> # Transition matrix:
> Phit := ⟨⟨A[1, 1, t]|A[1, 2, t]|A[1, 3, t]|1 − A[1, 1, t]-A[1, 2, t] − A[1, 3, t]⟩, ⟨A[2, 1, t]|A[2,
      2, t]|A[2, 3, t]|1 -A[2, 1, t]-A[2, 2, t] − A[2, 3, t]⟩, ⟨0|0|0|1⟩, ⟨0|0|0|1⟩⟩;
```

$$Phit := \begin{bmatrix} A_{1,1,t} & A_{1,2,t} & A_{1,3,t} & 1 - A_{1,1,t} - A_{1,2,t} - A_{1,3,t} \\ A_{2,1,t} & A_{2,2,t} & A_{2,3,t} & 1 - A_{2,1,t} - A_{2,2,t} - A_{2,3,t} \\ 0 & 0 & 0 & 1 \\ 0 & 0 & 0 & 1 \end{bmatrix} \tag{2}$$

```
> # Observation matrix:
> B := ⟨⟨1|0|0⟩, ⟨0|0|1⟩, ⟨0|lambda[t]|1 − λ[t]⟩, ⟨0|0|1⟩⟩;
```

$$B := \begin{bmatrix} 1 & 0 & 0 \\ 0 & 0 & 1 \\ 0 & \lambda_t & 1 - \lambda_t \\ 0 & 0 & 1 \end{bmatrix} \tag{3}$$

```
> B := Transpose(B);
```

$$B := \begin{bmatrix} 1 & 0 & 0 & 0 \\ 0 & 0 & \lambda_t & 0 \\ 0 & 1 & 1 - \lambda_t & 1 \end{bmatrix} \tag{4}$$

```
> # First time matrices:
> B0 := B :
> Phit0 := Phit :
>
```

```
> # T number of years (N +1 number of observation options, here 1,2 or 3)
> # Finds every possible capture history. Removes histories that are not possible e.g. 2 deaths
> T := 3 : N := 2 :
  H2 := Matrix((N + 1)^T, T) :
  for i from 1 to (N + 1)^T do
     nextone := i - 1 :
     for j from 1 to T do
        H2[i, j] := nextone mod (N + 1) + 1 :
        nextone := trunc(nextone / (N + 1)) :
     end do:
   end do:
  H3 := Matrix(1, Dimension(H2)[2]) :
   for i from 1 to Dimension(H2)[1] do
     no2 := 0 :
     for j from 1 to Dimension(H2)[2] do
        if H2[i, j] = 2 then
           no2 := no2 + 1 :
        end if:
     end do:
     if no2 ≤ 1 then
        H3 := ⟨H3, H2[i, 1 ..Dimension(H2)[2]]⟩ :
    end if:
   end do:
  H3 := H3[2 ..Dimension(H3)[1], 1 ..Dimension(H3)[2]] :
  H := Matrix(1, Dimension(H3)[2]) :
  for i from 1 to Dimension(H3)[1] do
      index2 := Dimension(H3)[2] + 1 :
      for j from 1 to Dimension(H3)[2] do
         if H3[i, j] = 2 then
            index2 := j :
        end if:
     end do:
      index1 := 0 :
     firstyes := 0 :
      for j from Dimension(H3)[2] by -1 to 1 do
         if firstyes = 0 then
            if H3[i, j] = 1 then
               index1 := j :
               firstyes := 1 :
            end if:
         end if:
     end do:
    if index1 > 0 then
     if index2 > index1 then
        H := ⟨H, H3[i, 1 ..Dimension(H3)[2]]⟩ :
    end if:
   end if:
   end do:
  H := H[2 ..Dimension(H)[1], 1 ..Dimension(H)[2]] :
```

```
> # Finds the probability of each history and creates the exhaustive summary of probabilities, κ
> κ := Vector(Dimension(H)[1]) :
  for j from 1 to Dimension(H)[1] do
      h := Row(H, j) :
      # e is first nonzero entry of h
      indi := 0 :
      for k from 1 to Dimension(H)[2] do
          if indi = 0 then
              if h[k] = 1 then
                  e := k :
                  indi := 1 :
              end if:
          end if:
      end do:
      Ph := Multiply(eval(Pit, t = e), DiagonalMatrix(Row(B0, h[e])));
      for i from e + 1 to T do
          if i − 1 = e then
              Ph := Multiply(Ph, Multiply(eval(Phit0, t = i − 1), DiagonalMatrix(Row(eval(B, t
      = i), h[i])))) :
          else
              Ph := Multiply(Ph, Multiply(eval(Phit, t = i − 1), DiagonalMatrix(Row(eval(B, t = i),
      h[i])))) :
          end if:
      end do:
      Ph := Multiply(Ph, Vector(Dimension(Ph), 1));
      κ[j] := Ph :
  end do:
> kappa[1 ..10]
```

$$\begin{bmatrix} A_{1,1,1} A_{1,1,2} \\ A_{1,1,2} \\ A_{1,2,1} A_{2,1,2} \\ 1 \\ A_{1,1,1} A_{1,3,2} \lambda_3 \\ A_{1,3,2} \lambda_3 \\ A_{1,2,1} A_{2,3,2} \lambda_3 \\ A_{1,1,1} A_{1,2,2} + A_{1,1,1} A_{1,3,2} (1-\lambda_3) + A_{1,1,1} (1 - A_{1,1,2} - A_{1,2,2} - A_{1,3,2}) \\ A_{1,3,2} (1-\lambda_3) + 1 - A_{1,1,2} - A_{1,3,2} \\ A_{1,3,1} \lambda_2 \end{bmatrix} \quad (5)$$

```
> # Vector of parameters
> pars := ⟨seq(indets(kappa)[i], i = 1 ..nops(indets(kappa)))⟩ :
> # D1 stores the derivative matrix which is calculated using proceedure Dmat
```

> $D1 := Dmat(\text{kappa}, pars):$

> # r stores the model rank, which is calculated using Maple's intrinsic proceedure rank. d stores the deficency, the difference between the number of parameters and the rank. A deficency of one or more indicates the model is parameter redundant, so this model is parameter redundant.

> $r := Rank(D1); d := Dimension(pars) - r;$

$$r := 6$$
$$d := 5 \qquad (6)$$

>

> # Tries to find the estimable parameter combinations

> $Estpar(D1, pars, 0);$

$$\left\{f\left(A_{1,1,1}, A_{1,1,2}, A_{1,2,1}, A_{1,2,2}, A_{1,3,1}, A_{1,3,2}, A_{2,1,2}, A_{2,2,2}, A_{2,3,2}, \lambda_2, \lambda_3\right) = \_F1\left(A_{1,1,1}, A_{1,1,2}, A_{1,2,1} A_{2,1,2}, \frac{A_{1,2,1} A_{2,3,2}}{A_{1,3,2}}, A_{1,3,1} \lambda_2, A_{1,3,2} \lambda_3\right)\right\} \qquad (7)$$

> # *Reparameterisation*

> $s := \left\langle A_{1,1,1}, A_{1,1,2}, A_{1,2,1} A_{2,1,2}, A_{1,3,1} \lambda_2, A_{1,3,2} \lambda_3, \frac{A_{1,2,1} A_{2,3,2}}{A_{1,3,2}} \right\rangle;$

$$s := \begin{bmatrix} A_{1,1,1} \\ A_{1,1,2} \\ A_{1,2,1} A_{2,1,2} \\ A_{1,3,1} \lambda_2 \\ A_{1,3,2} \lambda_3 \\ \frac{A_{1,2,1} A_{2,3,2}}{A_{1,3,2}} \end{bmatrix} \qquad (8)$$

(9)

> $AA := solve\left(\{seq(s[i] = ss[i], i = 1..Dimension(s))\}, \{A[1,1,1], A[1,1,2], A_{2,1,2}, A_{1,3,1}, A_{1,3,2}, A_{2,3,2}\}\right)$

$$AA := \left\{A_{1,1,1} = ss_1, A_{1,1,2} = ss_2, A_{1,3,1} = \frac{ss_4}{\lambda_2}, A_{1,3,2} = \frac{ss_5}{\lambda_3}, A_{2,1,2} = \frac{ss_3}{A_{1,2,1}}, A_{2,3,2} = \frac{ss_6\, ss_5}{A_{1,2,1} \lambda_3}\right\} \qquad (10)$$

> $kappas := simplify(eval(\text{kappa}, AA));$

(11)

$$kappas := \begin{bmatrix} \text{1 .. 11 } Vector_{column} \\ \text{Data Type: anything} \\ \text{Storage: rectangular} \\ \text{Order: Fortran\_order} \end{bmatrix} \quad \textbf{(11)}$$

> $kappas[1..10]$

$$\begin{bmatrix} ss_1\, ss_2 \\ ss_2 \\ ss_3 \\ 1 \\ ss_1\, ss_5 \\ ss_5 \\ ss_6\, ss_5 \\ -ss_1\,(ss_2 + ss_5 - 1) \\ -ss_2 - ss_5 + 1 \\ ss_4 \end{bmatrix} \quad \textbf{(12)}$$

> $kappas[11]$

$$-ss_6\, ss_5 - ss_1 - ss_3 - ss_4 + 1 \quad \textbf{(13)}$$

> $parss := \langle seq(ss[i], i = 1..Dimension(s))\rangle :$

> $Ds := Dmat(kappas, parss) :$

> $r := Rank(Ds); d := Dimension(parss) - r;$

$$r := 6$$
$$d := 0 \quad \textbf{(14)}$$

> # A PLUR decomposition of Ds, and finding the determinat of u1
> $(pp, ll, u1, r1) := LUDecomposition(\ Ds, output = [\text{'P','L','U1','R'}]\ ) :$
> $DetU := Determinant(u1);$

$$DetU := ss_1\, ss_5\, ss_2 \quad \textbf{(15)}$$

> $u1$

**(16)**

$$\begin{bmatrix} ss_2 & 0 & 0 & ss_5 & 0 & 0 \\ 0 & 1 & 0 & -\frac{ss_1\, ss_5}{ss_2} & 0 & 0 \\ 0 & 0 & 1 & 0 & 0 & 0 \\ 0 & 0 & 0 & ss_1 & ss_6 & 0 \\ 0 & 0 & 0 & 0 & ss_5 & 0 \\ 0 & 0 & 0 & 0 & 0 & 1 \end{bmatrix} \quad \textbf{(16)}$$

> *r1*

$$\begin{bmatrix} \textit{6 x 11 Matrix} \\ \textit{Data Type: anything} \\ \textit{Storage: rectangular} \\ \textit{Order: Fortran\_order} \end{bmatrix} \quad \textbf{(17)}$$

> *r1*[1 ..6, 1 ..10]; *r1*[1 ..6, 11];

$$\begin{bmatrix} 1 & 0 & 0 & 0 & 0 & -\frac{ss_5}{ss_1\, ss_2} & 0 & -\frac{ss_2 - 1}{ss_2} & \frac{ss_5}{ss_1\, ss_2} & 0 \\ 0 & 1 & 0 & 0 & 0 & \frac{ss_5}{ss_2} & 0 & -\frac{ss_1}{ss_2} & -\frac{ss_5 + ss_2}{ss_2} & 0 \\ 0 & 0 & 1 & 0 & 0 & 0 & 0 & 0 & 0 & 0 \\ 0 & 0 & 0 & 0 & 1 & \frac{1}{ss_1} & 0 & -1 & -\frac{1}{ss_1} & 0 \\ 0 & 0 & 0 & 0 & 0 & 0 & 1 & 0 & 0 & 0 \\ 0 & 0 & 0 & 0 & 0 & 0 & 0 & 0 & 0 & 1 \end{bmatrix}$$

$$\begin{bmatrix} -\frac{1}{ss_2} \\ \frac{ss_1}{ss_2} \\ -1 \\ 0 \\ -1 \\ -1 \end{bmatrix} \quad \textbf{(18)}$$

> *ll*

$$\begin{bmatrix} 1 & 0 & 0 & 0 & 0 & 0 \\ \frac{ss_1}{ss_2} & 1 & 0 & 0 & 0 & 0 \\ 0 & 0 & 1 & 0 & 0 & 0 \\ 0 & 0 & 0 & 1 & 0 & 0 \\ 0 & 0 & 0 & 0 & 1 & 0 \\ 0 & 0 & 0 & 0 & 0 & 1 \end{bmatrix} \quad (19)$$

```
> # Only an issue if certain values of ss are 0
> s
```

$$\begin{bmatrix} A_{1,1,1} \\ A_{1,1,2} \\ A_{1,2,1} A_{2,1,2} \\ A_{1,3,1} \lambda_2 \\ A_{1,3,2} \lambda_3 \\ \frac{A_{1,2,1} A_{2,3,2}}{A_{1,3,2}} \end{bmatrix} \quad (20)$$

```
> # Extension
> T := 4 : N := 2 :
  H2 := Matrix((N+1)^T, T) :
  for i from 1 to (N+1)^T do
     nextone := i - 1 :
     for j from 1 to T do
        H2[i, j] := nextone mod (N+1) + 1 :
        nextone := trunc(nextone/(N+1)) :
     end do:
   end do:
  H3 := Matrix(1, Dimension(H2)[2]) :
   for i from 1 to Dimension(H2)[1] do
     no2 := 0 :
     for j from 1 to Dimension(H2)[2] do
        if H2[i, j] = 2 then
           no2 := no2 + 1 :
        end if:
     end do:
     if no2 ≤ 1 then
        H3 := ⟨H3, H2[i, 1 ..Dimension(H2)[2]]⟩ :
     end if:
   end do:
  H3 := H3[2 ..Dimension(H3)[1], 1 ..Dimension(H3)[2]] :
```

```
H := Matrix(1, Dimension(H3)[2]) :
for i from 1 to Dimension(H3)[1] do
    index2 := Dimension(H3)[2] + 1 :
    for j from 1 to Dimension(H3)[2] do
        if H3[i, j] = 2 then
            index2 := j :
        end if:
    end do:
    index1 := 0 :
    firstyes := 0 :
    for j from Dimension(H3)[2] by -1 to 1 do
        if firstyes = 0 then
            if H3[i, j] = 1 then
                index1 := j :
                firstyes := 1 :
            end if:
        end if:
    end do:
    if index1 > 0 then
    if index2 > index1 then
        H := ⟨H, H3[i, 1 ..Dimension(H3)[2]]⟩ :
    end if:
  end if:
  end do:
H := H[2 ..Dimension(H)[1], 1 ..Dimension(H)[2]] :
```

```
> # Finds the probability of each history and creates the exhaustive summary of probabilities, κ
```

```
> κ := Vector(Dimension(H)[1]) :
  for j from 1 to Dimension(H)[1] do
      h := Row(H, j) :
      # e is first nonzero entry of h
      indi := 0 :
      for k from 1 to Dimension(H)[2] do
          if indi = 0 then
              if h[k] = 1 then
                  e := k :
                  indi := 1 :
              end if:
          end if:
      end do:
      Ph := Multiply(eval(Pit, t = e), DiagonalMatrix(Row(B0, h[e])));
      for i from e + 1 to T do
          if i − 1 = e then
              Ph := Multiply(Ph, Multiply(eval(Phit0, t = i − 1), DiagonalMatrix(Row(eval(B, t
    = i), h[i])))) :
          else
              Ph := Multiply(Ph, Multiply(eval(Phit, t = i − 1), DiagonalMatrix(Row(eval(B, t = i),
    h[i])))) :
          end if:
      end do:
```

```
    Ph := Multiply(Ph, Vector(Dimension(Ph), 1));
    κ[j] := Ph :
  end do:
> kappa
```

$$\left[\begin{array}{c} \mathit{1 .. 26\ Vector}_{column} \\ \mathit{Data\ Type:\ anything} \\ \mathit{Storage:\ rectangular} \\ \mathit{Order:\ Fortran\_order} \end{array}\right] \quad (21)$$

```
> # Vector of parameters
> pars := ⟨seq(indets(kappa)[i], i = 1 ..nops(indets(kappa)))⟩ :
> # D1 stores the derivative matrix which is calculated using proceedure Dmat
> D1 := Dmat(kappa, pars) :
> # r stores the model rank, which is calculated using Maple's intrinsic proceedure rank. d stores the deficency, the difference between the number of parameters and the rank. A deficency of one or more indicates the model is parameter redundant, so this model is parameter redundant.
> r := Rank(D1); d := Dimension(pars) − r;
```

$$r := 11$$
$$d := 7 \quad (22)$$

```
>
> # Tries to find the estimable parameter combinations
> Estpar(D1, pars, 0);
```

$$\left\{f\left(A_{1,1,1}, A_{1,1,2}, A_{1,1,3}, A_{1,2,1}, A_{1,2,2}, A_{1,2,3}, A_{1,3,1}, A_{1,3,2}, A_{1,3,3}, A_{2,1,2}, A_{2,1,3}, A_{2,2,2}, A_{2,2,3}, A_{2,3,2}, A_{2,3,3}, \lambda_2, \lambda_3, \lambda_4\right) = \_F1\left(A_{1,1,1}, A_{1,1,2}, A_{1,1,3}, A_{1,2,1} A_{2,1,2}, A_{1,2,2} A_{2,1,3}, \frac{A_{1,2,1} A_{2,2,2}}{A_{1,2,2}}, \frac{A_{1,2,1} A_{2,3,2}}{A_{1,3,2}}, \frac{A_{1,2,2} A_{2,3,3}}{A_{1,3,3}}, A_{1,3,1} \lambda_2, A_{1,3,2} \lambda_3, A_{1,3,3} \lambda_4\right)\right\} \quad (23)$$

```
> # Reparameterisation
> s := ⟨...⟩ :
```

$$s := \left\langle A_{1,1,1}, A_{1,1,2}, A_{1,2,1} A_{2,1,2}, A_{1,3,1} \lambda_2, A_{1,3,2} \lambda_3, \frac{A_{1,2,1} A_{2,3,2}}{A_{1,3,2}}, A_{1,1,3}, A_{1,2,2} A_{2,1,3}, A_{1,3,3} \lambda_4, \frac{A_{1,2,2} A_{2,3,3}}{A_{1,3,3}}, \frac{A_{1,2,1} A_{2,2,2}}{A_{1,2,2}} \right\rangle :$$

(24)

```
> AA := solve({seq(s[i] = ss[i], i = 1 ..Dimension(s))}, {A[1, 1, 1], A[1, 1, 2], A[2,1,2], A[1,3,1], A[1,3,2], A[2,3,2], A[1,1,3], A[2,1,3], A[1,3,3], A[2,3,3], A[2,2,2]})
```

$$AA := \left\{A_{1,1,1} = ss_1, A_{1,1,2} = ss_2, A_{1,1,3} = ss_7, A_{1,3,1} = \frac{ss_4}{\lambda_2}, A_{1,3,2} = \frac{ss_5}{\lambda_3}, A_{1,3,3} = \frac{ss_9}{\lambda_4}, A_{2,1,2} = \frac{ss_3}{A_{1,2,1}}, A_{2,1,3} = \frac{ss_8}{A_{1,2,2}}, A_{2,2,2} = \frac{ss_{11} A_{1,2,2}}{A_{1,2,1}}, A_{2,3,2} = \frac{ss_6\, ss_5}{A_{1,2,1} \lambda_3}, A_{2,3,3} = \frac{ss_{10}\, ss_9}{A_{1,2,2} \lambda_4}\right\} \quad (25)$$

> $kappas2 := simplify(eval(\text{kappa}, AA)) : indets(kappas2)$

$$\{ss_1, ss_2, ss_3, ss_4, ss_5, ss_6, ss_7, ss_8, ss_9, ss_{10}, ss_{11}\} \quad (26)$$

> $kappas[1..9]; kappas[10..11]$

$$\left[\begin{array}{c} ss_1\, ss_2 \\ ss_2 \\ ss_3 \\ 1 \\ ss_1\, ss_5 \\ ss_5 \\ ss_6\, ss_5 \\ -ss_1\,(ss_2 + ss_5 - 1) \\ -ss_2 - ss_5 + 1 \end{array}\right]$$

$$\left[\begin{array}{c} ss_4 \\ -ss_6\, ss_5 - ss_1 - ss_3 - ss_4 + 1 \end{array}\right] \quad (27)$$

> $kappas2[1..10]$

$$\left[\begin{array}{c} ss_1\, ss_2\, ss_7 \\ ss_2\, ss_7 \\ ss_3\, ss_7 \\ ss_7 \\ ss_1\, ss_8 \\ ss_8 \\ ss_{11}\, ss_8 \\ 1 \\ ss_1\, ss_2\, ss_9 \\ ss_2\, ss_9 \end{array}\right] \quad (28)$$

> $kappas2[11..20]$

$$\begin{bmatrix} ss_3\, ss_9 \\ ss_9 \\ ss_1\, ss_{10}\, ss_9 \\ ss_{10}\, ss_9 \\ ss_{11}\, ss_{10}\, ss_9 \\ -ss_1\, ss_2\, (ss_7 + ss_9 - 1) \\ -ss_2\, (ss_7 + ss_9 - 1) \\ -ss_3\, (ss_7 + ss_9 - 1) \\ -ss_7 - ss_9 + 1 \\ ss_1\, ss_5 \end{bmatrix} \tag{29}$$

> $s[1..9];\ s[10..11]$

$$\begin{bmatrix} A_{1,1,1} \\ A_{1,1,2} \\ A_{1,2,1}\, A_{2,1,2} \\ A_{1,3,1}\, \lambda_2 \\ A_{1,3,2}\, \lambda_3 \\ \frac{A_{1,2,1}\, A_{2,3,2}}{A_{1,3,2}} \\ A_{1,1,3} \\ A_{1,2,2}\, A_{2,1,3} \\ A_{1,3,3}\, \lambda_4 \end{bmatrix}$$

$$\begin{bmatrix} \frac{A_{1,2,2}\, A_{2,3,3}}{A_{1,3,3}} \\ \frac{A_{1,2,1}\, A_{2,2,2}}{A_{1,2,2}} \end{bmatrix} \tag{30}$$

> $parss := \langle seq(ss[i], i = 1..Dimension(s))\rangle :$

> $Ds := Dmat(kappas, parss) :$

> $r := Rank(Ds);\ d := Dimension(parss) - r;$

$$r := 6$$

$$d := 5 \tag{31}$$

```
> # A PLUR decomposition of Ds, and finding the determinat of u1
(pp, ll, u1, r1) := LUDecomposition( Ds, output = [ 'P','L','U1','R' ] ) :
DetU := Determinant(u1);
```

$$DetU := ss_1\, ss_5\, ss_2 \tag{32}$$

```
>
```
