## Supplementary material for "Size- and stage-dependence in cause-specific mortality of migratory brown trout": Code for custom model implementation with NIMBLE: nimbleDHMM.R

```
require(nimble)

dtroutDHMM <- nimble::nimbleFunction(
    run = function(
        ## argument type declarations
        x = double(1), length = double(), origin = double(),
        mu.mH1 = double(), mu.mO1 = double(), mu.mO2 = double(), mu.p = double(),
        beta1.mO1 = double(), beta1.mO2 = double(), beta2.mO1 = double(), beta2.mO2 = double(), betaS.mO = double(),
        beta2.mH1 = double(), beta4.mH1 = double(), betaS.mH = double(),
        beta1.p = double(), beta2.p = double(), beta3.p = double(), beta4.p = double(), betaS.p = double(),
        size = double(1), epsilon.mH = double(1), epsilon.mO = double(1), epsilon.p = double(1), 
        discF = double(1), discS = double(1), r = double(1), log = double()) {
        
        ## calculate background and harvest mortality hazard rates,
        ## ladder usage probability, and survival probabilities
        mH1 <- exp(mu.mH1 + betaS.mH*origin + beta2.mH1*size + beta4.mH1*size^2 + epsilon.mH)
        mO1 <- exp(mu.mO1 + betaS.mO*origin + beta1.mO1*discF + beta2.mO1*size + epsilon.mO)
        mO2 <- exp(mu.mO2 + betaS.mO*origin + beta1.mO2*discF + beta2.mO2*size + epsilon.mO)
        p <- expit(mu.p + betaS.p*origin + beta1.p*discS + beta2.p*size + beta3.p*discS*size + beta4.p*size^2 + epsilon.p
        S1 <- exp(-(mH1 + mO1))
        alpha1 <- mH1 / (mH1 + mO1)
        S2 <- exp(-(mH1 + mO2))
        alpha2 <- mH1 / (mH1 + mO2)
        
        logL <- 0                             ## initialize log-likelihood
        pi <- numeric(4, init = FALSE)         
        pi[1] <- S1[1]*p[2]                   ## calculate probability of each 
        pi[2] <- S1[1]*(1-p[2])               ## unobserved state, conditional
        pi[3] <- (1-S1[1])*alpha1[1]          ## on first observation of spawning
        pi[4] <- (1-S1[1])*(1-alpha1[1])
        for(t in 2:length) {                  ## iterate over remaining observations
            Zpi <- pi
            if(x[t] == 1) { Zpi[2] <- 0       ## observed and seen alive:
                            Zpi[3] <- 0       ## update conditional distribution of unobserved
                            Zpi[4] <- 0 }     ## states, given this observation
            if(x[t] == 2) { Zpi[1] <- 0       ## harvested and reported:
                            Zpi[2] <- 0       ## update conditional distribution
                            Zpi[3] <- pi[3] * r[t]
                            Zpi[4] <- 0 }
            if(x[t] == 3) { Zpi[1] <- 0       ## not seen or reported
                            Zpi[3] <- pi[3] * (1-r[t]) }
            sumZpi <- sum(Zpi)
            logL <- logL + log(sumZpi)        ## log-likelihood contribution of observed state
            if(t != length) {                 ## distribution of unobserved state for next time period
                pi[1] <- Zpi[1]*S1[t]*p[t+1]            + Zpi[2]*S2[t]*p[t+1]
                pi[2] <- Zpi[1]*S1[t]*(1-p[t+1])        + Zpi[2]*S2[t]*(1-p[t+1])
                pi[3] <- Zpi[1]*(1-S1[t])*alpha1[t]     + Zpi[2]*(1-S2[t])*alpha2[t]
                pi[4] <- Zpi[1]*(1-S1[t])*(1-alpha1[t]) + Zpi[2]*(1-S2[t])*(1-alpha2[t])+Zpi[3]+Zpi[4]
                pi <- pi / sumZpi             ## normalize
            }
        }
        returnType(double())
        if(log) return(logL) else return(exp(logL))    ## return log-likelihood
    }
)

rtroutDHMM <- nimble::nimbleFunction(
    run = function(
        n = integer(), length = double(), origin = double(),
        mu.mH1 = double(), mu.mO1 = double(), mu.mO2 = double(), mu.p = double(),
        beta1.mO1 = double(), beta1.mO2 = double(), beta2.mO1 = double(), beta2.mO2 = double(), betaS.mO = double(), 
        beta2.mH1 = double(), beta4.mH1 = double(), betaS.mH = double(),
        beta1.p = double(), beta2.p = double(), beta3.p = double(), beta4.p = double(), betaS.p = double(),
        size = double(1), epsilon.mH = double(1), epsilon.mO = double(1), epsilon.p = double(1), 
        discF = double(1), discS = double(1), r = double(1)) {
        x <- rep(1, length)
        returnType(double(1))
        return(x)
    }
)

nimble::registerDistributions(list(
    dtroutDHMM = list(
        BUGSdist = 'dtroutDHMM(length, origin, mu.mH1, mu.mO1, mu.mO2, mu.p, beta1.mO1, beta1.mO2, beta2.mO1, beta2.mO2, betaS.mO, beta2.mH1, beta4.mH1, betaS.mH, beta1.p, beta2.p, beta3.p, beta4.p, betaS.p, size, epsilon.mH, epsilon.mO, epsilon.p, discF, discS, r)',
        types = c('value = double(1)', 'size = double(1)', 'epsilon.mH = double(1)', 'epsilon.mO = double(1)', 'epsilon.p = double(1)', 'discF = double(1)', 'discS = double(1)', 'r = double(1)'),
        discrete = TRUE
    )
))

trout.code.DHMM <- nimbleCode({

	  #---------------#
	  # SHARED PRIORS # 
	  #---------------#
    
      Mu.mH1 ~ dunif(0.01, 3)
      mu.mH1 <- log(Mu.mH1)
      
      Mu.mO1 ~ dunif(0.01, 3)
      mu.mO1 <- log(Mu.mO1)
      
      Mu.mO2 ~ dunif(0.01, 3)
      mu.mO2 <- log(Mu.mO2)
    
      Mu.p ~ dunif(0.01, 0.99)
      mu.p <- log(Mu.p/(1-Mu.p))
      
	  Mu.r1 ~ dunif(0.01, 0.99)
    
      beta1.p ~ dunif(-2, 2)
      beta2.p ~ dunif(-2, 2)
      beta3.p ~ dunif(-2, 2)
      beta4.p ~ dunif(-2, 2)
      betaS.p ~ dunif(-2, 2)

      beta2.mH1 ~ dunif(-5, 5)
      beta4.mH1 ~ dunif(-5, 5)
      betaS.mH ~ dunif(-5, 5)
      
      beta1.mO1 ~ dunif(-5, 5)
      beta2.mO1 ~ dunif(-5, 5)
      beta1.mO2 ~ dunif(-5, 5)
      beta2.mO2 ~ dunif(-5, 5)
      betaS.mO ~ dunif(-5, 5)

      sigma.mH ~ dunif(0, 5)
      sigma.mO ~ dunif(0, 5)
      sigma.p ~ dunif(0, 5)
      sigma.r ~ dunif(0, 5)
      
      #-----------------------------------------#
      # ANALYSIS OF DATA FOR EVEN-YEAR SPAWNERS #
      #-----------------------------------------#
      
	  for(t in 1:(n.occasions.even-1)){

        epsilon.mH.even[t] ~ dnorm(0, sd = sigma.mH)
        epsilon.mO.even[t] ~ dnorm(0, sd = sigma.mO)
        epsilon.p.even[t+1] ~ dnorm(0, sd = sigma.p) 
        epsilon.r.even[t+1] ~ dnorm(0, sd = sigma.r) 

        logit(r.even[t+1]) <- logit(r.odd[t]) + epsilon.r.even[t+1]       
        
      }
    
      epsilon.mH.even[n.occasions.even] <- 0
      epsilon.mO.even[n.occasions.even] <- 0
      epsilon.p.even[1] <- 0
      epsilon.r.even[1] <- 0
      r.even[1] <- Mu.r1
      

      for(i in 1:nind.even) {
        y.even[i, first.even[i]:last.even[i]] ~ dtroutDHMM(length = last.even[i]-first.even[i]+1, origin=even.origin[i],
                                               	mu.mH1=mu.mH1, mu.mO1=mu.mO1, mu.mO2=mu.mO2, mu.p=mu.p,
                                               	beta1.mO1=beta1.mO1, beta1.mO2=beta1.mO2, 
                                               	beta2.mO1=beta2.mO1, beta2.mO2=beta2.mO2, betaS.mO=betaS.mO, 
                                                beta2.mH1=beta2.mH1, beta4.mH1=beta4.mH1, betaS.mH=betaS.mH,
                                                beta1.p=beta1.p, beta2.p=beta2.p, beta3.p=beta3.p, beta4.p=beta4.p, betaS.p=betaS.p,
                                                size=even.size[i,first.even[i]:last.even[i]],
                                                epsilon.mH=epsilon.mH.even[first.even[i]:last.even[i]],
                                                epsilon.mO=epsilon.mO.even[first.even[i]:last.even[i]],
                                                epsilon.p=epsilon.p.even[first.even[i]:last.even[i]],
                                                discF=even.discF[first.even[i]:last.even[i]], 
                                                discS=even.discS[first.even[i]:last.even[i]], 
                                                r=r.even[first.even[i]:last.even[i]])
      }
         
      
      #----------------------------------------#
      # ANALYSIS OF DATA FOR ODD-YEAR SPAWNERS #
      #----------------------------------------#
       		
       		
	  for(t in 1:(n.occasions.odd-1)){

        epsilon.mH.odd[t] ~ dnorm(0, sd = sigma.mH)
        epsilon.mO.odd[t] ~ dnorm(0, sd = sigma.mO) 
        epsilon.p.odd[t+1] ~ dnorm(0, sd = sigma.p)      
        epsilon.r.odd[t+1] ~ dnorm(0, sd = sigma.r) 

        logit(r.odd[t+1]) <- logit(r.even[t+1]) + epsilon.r.odd[t+1]  
               
      }
      
      epsilon.mH.odd[n.occasions.odd] <- 0
      epsilon.mO.odd[n.occasions.odd] <- 0
      epsilon.p.odd[1] <- 0
      
      epsilon.r.odd[1] ~ dnorm(0, sd = sigma.r)
      logit(r.odd[1]) <- logit(r.even[1]) + epsilon.r.odd[1]
 
      
      for(i in 1:nind.odd) {
        y.odd[i, first.odd[i]:last.odd[i]] ~ dtroutDHMM(length = last.odd[i]-first.odd[i]+1, origin = odd.origin[i],
                                               	mu.mH1=mu.mH1, mu.mO1=mu.mO1, mu.mO2=mu.mO2, mu.p=mu.p,
                                               	beta1.mO1=beta1.mO1, beta1.mO2=beta1.mO2, 
                                               	beta2.mO1=beta2.mO1, beta2.mO2=beta2.mO2, betaS.mO=betaS.mO, 
                                                beta2.mH1=beta2.mH1, beta4.mH1=beta4.mH1, betaS.mH=betaS.mH,
                                                beta1.p=beta1.p, beta2.p=beta2.p, beta3.p=beta3.p, beta4.p=beta4.p, betaS.p=betaS.p,
                                                size=odd.size[i,first.odd[i]:last.odd[i]],
                                                epsilon.mH=epsilon.mH.odd[first.odd[i]:last.odd[i]],
                                                epsilon.mO=epsilon.mO.odd[first.odd[i]:last.odd[i]],
                                                epsilon.p=epsilon.p.odd[first.odd[i]:last.odd[i]],
                                                discF=odd.discF[first.odd[i]:last.odd[i]], 
                                                discS=odd.discS[first.odd[i]:last.odd[i]], 
                                                r=r.odd[first.odd[i]:last.odd[i]])
      } 
})
```
