## Supplementary material for "Size- and stage-dependence in cause-specific mortality of migratory brown trout": Code for custom model implementation with NIMBLE: readme.pdf

This R code corresponds to the file mentioned in Appendix S2 in the supplementary material of the paper ‘Size- and stage-dependence in cause-specific mortality of migratory brown trout’ by Chloé R. Nater, Yngvild Vindenes, Per Aass, Diana Cole,Øystein Langangen, S. Jannicke Moe, Atle Rustadbakken, DanielTurek, L. Asbjørn Vøllestad and Torbjørn Ergon

| R file | Explanation |
|---|---|
| nimbleDHMM.R | R code for specifying the custom distribution for NIMBLE and implementing it for the trout model. For detailed descriptions of all variables used in the code, refer to Appendix S2 of the article. |
